# High-variance phenome database reveals important roles of WD40 proteins in the plant pathogenic fungus *Fusarium graminearum*

**DOI:** 10.64898/2026.04.19.719521

**Authors:** Soyoung Choi, Nahyun Lee, Jiyeun Park, Hosung Jeon, Sieun Kim, Jung-Eun Kim, Jiyoung Shin, Heeji Moon, Kyunghun Min, Yejin Choi, Aram Hwangbo, Hun Kim, Gyung Ja Choi, Yin-Won Lee, Dae-Geun Song, Hokyoung Son

## Abstract

- WD40 is a highly conserved protein domain in eukaryotes that functions as a versatile platform for protein-protein interactions and participates in diverse biological processes.
- We performed a genome-wide functional analysis of WD40 domain-containing proteins in *Fusarium graminearum*, a phytopathogenic fungus that causes severe yield losses and mycotoxin contamination in major cereal crops.
- Comprehensive phenotypic profiling of 119 WD40 gene deletion mutants across 22 phenotypic traits established a systematic WD40 phenome dataset, revealing the broad functional involvement of WD40 proteins and a strong correlation between sexual reproduction and virulence.
- Protein interaction analyses of selected WD40 proteins revealed diverse WD40- mediated interaction patterns and provided further insights into WD40-mediated protein interactions and their roles in protein complex formation. This study provides a foundation for further characterization of WD40 proteins in filamentous fungi.

## Introduction

WD40 is a protein domain named after a conserved region containing a tryptophan-aspartic acid (WD) dipeptide at the C-terminus of each repeat (Neer *et al*., 1994; Smith *et al*., 1999). Each WD40 repeat contributes to the formation of a four-stranded antiparallel β-sheet blade, and multiple blades assemble into a characteristic β-propeller structure that provides a stable platform for protein interactions (Wall *et al*., 1995; Lambright *et al*., 1996; Sondek *et al*., 1996). WD40 is one of the most abundant protein families in eukaryotes (Wang *et al*., 2015). There are 743 WD40-coding genes in wheat (*Triticum aestivum* L.) (Hu *et al*., 2018), 200 in rice (Ouyang *et al*., 2012), 237 in *Arabidopsis thaliana* (van Nocker & Ludwig, 2003), and 262 in humans (Zou *et al*., 2016).

The WD40 family is a large and diverse protein family that plays versatile roles in protein–protein interactions (PPIs). WD40 proteins act as scaffolds for protein binding, thereby promoting the formation of protein complexes involved in diverse cellular processes, including signal transduction and gene expression regulation (Xu & Min, 2011; Wu *et al*., 2012; Zhang & Zhang, 2015). However, despite their functional importance, the characterization of WD40 family proteins remains challenging. WD40 proteins frequently interact with multiple binding partners, and the structural diversity of WD40 domains contributes to the formation of distinct interaction networks. Therefore, defining the biological functions and interaction partners of individual WD40 proteins remains an important challenge.

*Fusarium graminearum* is an important plant pathogenic fungus that causes Fusarium head blight (FHB) in wheat and barley, as well as ear rot in maize (Windels, 2000; Desjardins, 2006). *Fusarium* infection causes severe loss in yield and contamination with mycotoxins, such as zearalenone (ZEA) and deoxynivalenol (DON), which pose substantial risks to human and animal health (Windels, 2000; Goswami & Kistler, 2004; Desjardins & Proctor, 2007). To develop effective strategies for controlling FHB, extensive genetic and biochemical studies have been conducted to understand fungal development, stress responses, mycotoxin production, and pathogenicity in *F. graminearum*.

Genome-wide deletion mutant collections have served as powerful resources for systematically investigating gene functions and their contributions to biological processes. In *F. graminearum*, several genome-wide functional studies targeting specific gene families, including transcription factors (Son *et al*., 2011b), kinases (Wang *et al*., 2011), and phosphatases (Yun *et al*., 2015), have provided valuable insights into the genetic regulation of fungal development, stress responses, and pathogenicity. Given that WD40 proteins primarily function as scaffolds for protein interactions, integrating systematic phenotypic analysis with protein interaction studies provides an effective approach for understanding their functional diversity. However, despite their biological importance, a genome-wide functional investigation of WD40 proteins has not yet been conducted in filamentous fungi, including fungal pathogens.

In this study, we identified 157 genes encoding WD40 domain-containing proteins and generated a genome-wide deletion mutant library for non-essential WD40 genes in *F. graminearum*. Comprehensive phenotypic profiling of these mutants was performed across 22 phenotypic traits, including vegetative growth, sexual and asexual reproduction, pathogenicity, mycotoxin production, and responses to various stress agents. Based on this genome-wide phenome dataset, we analyzed relationships among WD40-associated phenotypes and further characterized selected WD40 proteins through protein interaction analyses. This study provides a comprehensive overview of the functional characteristics of WD40 proteins in *F. graminearum*, establishing a foundation for further investigation of their molecular functions in filamentous fungi.

## Materials and Methods

### Fungal strains and culture conditions

*F. graminearum* Schwabe wild-type strain Z-3639 was used as the parental strain throughout this study (Bowden & Leslie, 1999). Fungal strains were stored at −80 °C as mycelial suspensions in 20% glycerol solution. Culture media were prepared as described in the *Fusarium* laboratory manual (Leslie & Summerell, 2006). All fungal strains were grown at 25 °C.

### Identification of WD40 genes in *F. graminearum*

Genes encoding WD40 domain-containing proteins were identified from the FungiDB database (fungidb.org) (Stajich *et al*., 2012), with InterPro IDs used as a reference. The FungiDB database was used to analyze the chromosomal location of WD40 genes and obtain sequence information. To confirm the presence of WD40 motifs in the extracted WD40 protein sequences, we used the WD40-repeat Protein Structure Predictor (www.wdspdb.com/wdsp). To identify orthologs of *F. graminearum* WD40 proteins in *S. cerevisiae*, BLASTp analysis was performed using annotated information from the *Saccharomyces* Genome Database (www.yeastgenome.org). Hits with an *E*-value cut-off of <0.0001 were retained. The distribution of WD40 genes across chromosomes was mapped with MapGene2Crom (MG2C) (Chao *et al*., 2021), and the architecture of WD40 genes was visualized using IBS software (Illustrator for Biological Sequences) (Xie *et al*., 2022).

### Phylogenetic analysis and classification of putative WD40 genes

The amino acid sequences of WD40 proteins were aligned using ClustalW and MUSCLE algorithms (Thompson *et al*., 1994; Edgar, 2004). An unrooted neighbor-joining phylogenetic tree was constructed using MEGA11 (Molecular Evolutionary Genetics Analysis) software, with 1,000 bootstrap replicates for statistical support (Tamura *et al*., 2021). WD40 proteins were classified using the OmicsBox software (www.biobam.com/omicsbox). The finalized phylogenetic tree was visualized using iTOL (itol.embl.de) (Letunic & Bork, 2019).

### GO annotation and KEGG pathway enrichment analysis

GO IDs were identified using OmicsBox with default settings to identify biological functions of WD40 proteins (Götz *et al*., 2008). Functional annotation data of WD40 proteins were submitted to WEGO 2.0 (Web Gene Ontology Annotation Plot, wego.genomics.cn) (Ye *et al*., 2018) to generate GO classification histograms of WD40 proteins. In addition, KEGG enrichment analysis of WD40 genes was performed using KOBAS v3.0 (kobas.cbi.pku.edu.cn/) to obtain the KEGG IDs of WD40 proteins. The resulting graphs were generated using the R program.

### DNA extraction and Southern blot hybridization

Three-day-old mycelia grown in 5 mL of CM broth were lyophilized to extract genomic DNA using the CTAB method (Leslie & Summerell, 2006). Restriction endonuclease digestion and agarose gel electrophoresis were performed as previously described (Sambrook & Russell, 2001). Southern blot analysis was performed using the North2South Biotin Random Prime Labeling Kit and North2South Chemiluminescent Hybridization and Detection Kit (Thermo Scientific, Middlesex County, MA, USA). The ChemiDoc XRS+ Gel Imaging System (Bio-Rad, Hercules, CA, USA) was used to detect specific bands.

### Generation of WD40 gene deletion mutants and GFP-tagged Atg8 strain

WD40 genes were deleted using the split-marker method based on homologous recombination (Son *et al*., 2011a). The geneticin resistance gene cassette (*GEN*) was amplified from the pII99 plasmid, and 5′ and 3′ flanking regions of the target genes were amplified from wild-type genomic DNA. Three fragments were fused by double-joint PCR (Yu *et al*., 2004). PCR products were extracted using the MEGAquick-spin Plus Total Fragment DNA Purification Kit (Intron Biotech, Seongnam, Republic of Korea). Purified products were transformed into protoplasts of the wild-type strain.

To generate N-terminally GFP-tagged Atg8, the hygromycin resistance-GFP cassette (*HYG*-*GFP*), including the *Neurospora crassa* isocitrate lyase (ICL) promoter, was amplified from the pIGPAPA plasmid (Horwitz *et al*., 1999) and fused with the corresponding ATG8- targeting fragments to enable insertion immediately upstream of the ATG8 start codon. The resulting construct was introduced into protoplasts of the wild-type strain using the transformation procedure described above.

The transformants were screened using diagnostic PCR, and mutants were confirmed by Southern blot hybridization using either flanking region probes or ORF-specific probes. At least two independent mutants were obtained for each gene deletion. The oligonucleotides used were synthesized by Bioneer (Daejeon, Republic of Korea) (Table S1).

### Genetic manipulation for complementation and overexpression strains

For complementation of Δ*fgwd101* and Δ*fgwd133*, the ORF and native promoter of each gene were amplified from wild-type genomic DNA using PrimeSTAR (Takara Bio, Otsu, Japan). The resulting constructs and XhoI-digested pDL2 (Zhou *et al*., 2011) were co-transformed into the yeast strain PJ69-4A using the Alkali-Cation Yeast Transformation Kit (MP Biomedicals, Irvine, CA, USA). The vectors obtained from yeast transformants were transformed into *Escherichia coli* DH10B. Plasmid DNA was extracted with the DNA-spin Plasmid DNA Purification Kit (Intron Biotech) and verified by sequencing (Macrogen, Seoul, Republic of Korea).

To produce large quantities of protein, overexpression strains for *FgWD101* or *FgWD133* were generated. The ORF of each gene was amplified and co-transformed into the yeast strain PJ69-4A with XhoI-digested pDL2. *RP27-FgWD101-GFP* and *RP27-FgWD133- GFP* fusion vectors were obtained from the yeast transformants. The subsequent steps were performed as described above. The resulting fusion vectors were used for transforming the corresponding deletion mutants.

### Phenotypic analysis of vegetative growth, conidiation, and sexual development

For phenotypic analysis, at least two independent strains were used for each gene. Colony morphology was assayed on complete medium (CM) 4 d after inoculation. Conidiation assays were conducted by inoculating fresh mycelial blocks from each strain into 5 mL carboxymethyl cellulose (CMC) medium and incubating for 5 d on a rotary shaker (200 rpm). The number of conidia was determined using a hemocytometer (Paul Marienfeld, Lauda-Königshofen, Germany). Each experiment was repeated three times.

For sexual development analyses, each strain was inoculated on carrot agar medium, and after 5 d of incubation, aerial mycelia were removed with 0.4 mL of 2.5% sterile Tween 60 (Leslie & Summerell, 2006). The cultures were incubated under near-UV light conditions (wavelength: 365 nm; Sankyo Denki, Tokyo, Japan) for 7–9 d. The formation and maturation of perithecia were observed using a Zeiss SteREO Lumar.V12 microscope (Carl Zeiss, Oberkochen, Germany). Ascospore discharge was assessed by observing discharged ascospores on the medium lid. The morphology of rosettes, asci, and ascospores was examined using a Leica DM6 B microscope (Leica Microsystems, Wetzlar, Germany).

For outcrosses, the female strain was fertilized with 1 mL of conidial suspension (10^6^ conidia mL^−1^) from the male strain. Fertilization was performed 5 d after inoculating the strains on carrot agar. The Δ*mat1-1*/pIGPAPA*-*GFP strain was used to examine 1:1 segregation of ascospores in each ascus (Lee *et al*., 2003).

### Virulence test

The point inoculation method was employed to assess fungal virulence (Son *et al*., 2011b). Conidia were collected and resuspended in 0.01% Tween-20 at a concentration of 10^5^ conidia mL^−1^. For mutants showing reduced conidiation, larger volumes of CMC cultures were prepared to obtain sufficient numbers of conidia, and the conidial concentration was adjusted to the same level as that used for the wild-type strain prior to inoculation. Then, 10 μL of conidial suspension was injected into the central spikelet of a wheat head at the early anthesis stage (cv. Eunpamil). For mutants that completely failed to produce conidia, mycelial agar blocks were directly placed onto wheat heads to evaluate virulence. Inoculated wheat heads were covered with plastic bags to maintain humidity for 3 d. The number of spikelets exhibiting disease symptoms was evaluated 14 d after inoculation.

For the coleoptile virulence test, approximately 2 mm of the upper part of 3-d-old wheat seedlings was trimmed and inoculated with 2 μL of conidial suspension of each strain (10^6^ conidia mL^−1^). The inoculated seedlings were placed in a humid chamber, and disease symptoms were examined 7 d post-inoculation (dpi).

For the cellophane penetration assay, autoclaved cellophane sheets were used on CM plates. To determine if a strain penetrated the cellophane membrane, strains were cultured on CM with a cellophane membrane covering the surface. After 2 d of incubation, the cellophane membrane was removed from each plate. The plates were incubated for 2 d and photographed. All experiments were repeated three times independently.

### Mycotoxin analysis

To assess mycotoxin production, each strain was cultured on 1.5 g of sterilized rice substrate under dark conditions for 3 weeks. The rice culture was harvested and ground. Mycotoxins were extracted by vigorously mixing the ground rice culture with 6 mL of 84% acetonitrile (ACN) for 30 min at 25°C on a rotary shaker (200 rpm), resulting in phase separation. The upper layer was filtered through a 0.45-µm syringe filter (Merck Millipore, Burlington, MA, USA). Reverse-phase HPLC analysis was performed using a Prominence HPLC system (Shimadzu, Kyoto, Japan) equipped with a C18 column, with minor modifications (Kim *et al*., 2005; Ok *et al*., 2018). Zearalenone (ZEA) was detected using a mobile phase consisting of 70% aqueous methanol, at a flow rate of 1 mL min^−1^. Deoxynivalenol (DON) was detected with a mobile phase containing 10% aqueous ACN, at a flow rate of 1 mL min^−1^. The following gradient elution program was used for DON detection: 10% ACN was maintained for 11 min; increased linearly to 30% ACN at 12 min; decreased linearly from 30% ACN at 12 min to 10% ACN at 18 min; maintained for 17 min to re-equilibrate the column; and the next sample was injected, resulting in a total runtime of 35 min. Both ZEA and DON were detected at a wavelength of 235 nm using the diode-array detection.

### Stress tolerance analysis

Stress responses were examined using 96-well plates under various stress conditions, with the concentration of each stress agent adjusted to the half-maximal effective concentration (EC_50_) of the wild-type (Choi *et al*., 2023). Conidia were collected from CMC cultures, filtered using Miracloth (Merck Millipore), and resuspended in CM broth. All strains were normalized to a concentration of 10^5^ conidia mL^−1^. In 96-well plates, 100 μL of conidial suspension was mixed with an equal volume of CM supplemented with various stress agents: oxidative stress (0.31 mM hydrogen peroxide [H_2_O_2_]), osmotic stress (1 M sorbitol and 1 M NaCl), cell wall stress (0.1 mg mL^−1^ sodium dodecyl sulfate [SDS]), fungicide stress (0.05 μg mL^−1^ fludioxonil, 125 μg mL^−1^ iprodione, and 0.625 μg mL^−1^ tebuconazole), endoplasmic reticulum stress (2 μg mL^−1^ tunicamycin and 1.25 mM dithiothreitol [DTT]), and pH stress (pH 2 and 12). After inoculation, the plates were incubated in a humid plastic box for 48 h. OD600 was measured using a Synergy HTX microplate reader (BioTek, Winooski, VT, USA). The obtained values were adjusted by subtracting the average absorbance of the medium. The results were normalized with growth in CM, and log2 fold change was calculated based on the average growth. The heat map was generated using Morpheus software (software.broadinstitute.org/morpheus).

### Phenotypic correlation analysis

Phenotypic correlation analysis was performed using quantified phenotypic data from WD40 deletion mutants. For consistency across phenotypic traits, sexual reproduction-related phenotypes were converted into directional scores. Pairwise correlations between phenotypic traits were calculated using Spearman’s rank correlation coefficient. Correlation matrices were generated and visualized as heat maps using Python packages (version 3.11.15), including pandas, SciPy, and seaborn.

### Western blotting and immunoprecipitation

Fresh mycelia were harvested from 3-d-old CM culture, finely ground with liquid nitrogen, and suspended in 1 mL extraction buffer containing protease inhibitors (1 mM phenylmethyl sulfonyl fluoride, 20 μg mL^−1^ leupeptin, 2 μg mL^−1^ pepstatin A, and 20 μg mL^−1^ aprotinin). For analysis of autophagic activity, mycelia were cultured in CM for 24 h and then transferred to nitrogen starvation medium (MM-N) as previously described (Zheng *et al*., 2018) for 2 h before protein extraction. The samples were sonicated using an ultrasonic homogenizer, and cell lysates were collected by centrifugation at 16,000 ***g*** for 20 min at 4°C.

For immunoprecipitation (IP), GFP-tagged proteins were pulled down using anti-GFP monoclonal antibody-conjugated magnetic beads (Code No. D153-11, MBL, Tokyo, Japan). For co-immunoprecipitation (co-IP), strains expressing both GFP- and FLAG-tagged proteins were generated, and protein complexes were immunoprecipitated using anti-GFP magnetic beads. The immunoprecipitated proteins were analyzed by western blotting.

Proteins were separated using SDS-PAGE and transferred to nitrocellulose membranes using the Trans-Blot Turbo Transfer System (Bio-Rad, Hercules, CA, USA). The membranes were blocked with 7% blotting grade blocker in TBS-T buffer. Primary antibodies against GFP (Cat# ab290, Abcam, Cambridge, UK), FLAG (Cat# A00170, GenScript), and GAPDH (Cat# A19056, ABclonal) were used for immunoblotting. Goat anti-rabbit IgG-HRP antibody (Cat# ab6721, Abcam) was used as the secondary antibody. Signals were detected using Amersham ECL Select Western Blotting Detection Reagent (Cytiva, Marlborough, MA, USA) and photographed using ChemiDoc XRS+ Gel Imaging System (Bio-Rad).

### Affinity purification mass spectrometry (AP-MS)

Protein samples were separated using 4%–20% precast SDS-PAGE gels and stained with Coomassie Brilliant Blue (Cat# 4561095, Bio-Rad). The corresponding deletion mutants were used as negative controls to identify non-specific proteins bound to anti-GFP mAb-conjugated magnetic beads. After excluding non-specific bands detected in the negative controls, bands of interest were excised and digested with trypsin. Tryptic peptides were analyzed using a Q Exactive system (Thermo Scientific) (Kim *et al*., 2018).

### Yeast two-hybrid assay

Yeast Two-hybrid (Y2H) assay was performed using Matchmaker Gold Yeast Two-Hybrid System (Takara Bio, CA, USA), according to the manufacturer’s instructions. The indicated full-length or truncated coding sequences of candidate interacting proteins were amplified from cDNA of the wild-type strain Z-3639 and cloned into the corresponding bait and prey vectors. WD40 protein-encoding sequences, including *FgWD101*, *FgWD133*, Atg2, and the truncated structure of *FgWD101*, were cloned into pGADT7 as prey constructs. The sequences encoding their putative interacting partners, including Fgwd072, septin proteins, and *FgWD048* variants, were inserted into pGBKT7 as baits. The bait and prey constructs were co-transformed into *S. cerevisiae* strain Y2HGold. Transformants were selected on SD medium lacking leucine and tryptophan (SD/−Leu/−Trp), and protein–protein interactions were assessed on SD lacking histidine, leucine, and tryptophan (SD/−His/−Leu/−Trp).

### Prediction of protein interaction interfaces

Protein complex structures were predicted using AlphaFold2-Multimer via the ColabFold platform with the amino acid sequences of the corresponding protein pairs as input (Evans *et al*., 2022; Mirdita *et al*., 2022). Default prediction parameters were used. Putative interface residues between the two proteins were identified using the Interfaces tool in UCSF ChimeraX version 1.9 (Meng *et al*., 2023), excluding solvent molecules and using the default interface analysis parameters.

## Results

### Identification and classification of WD40 genes in *F. graminearum*

To identify WD40 genes in *F. graminearum*, we searched for WD40 repeat-containing domains and WD40 domain-encoding genes in FungiDB (fungidb.org) (Stajich *et al*., 2012) based on the InterPro database. Duplicates of InterPro IDs were removed, and 157 putative WD40-encoding genes were identified (Table S2). For convenience, we designated the identified genes as *FgWD001*–*FgWD157* (*F. graminearum* WD40 gene No.), and genes named in previous studies were referred to collectively. To verify the gene list, we used WDSP (WD40-repeat protein Structure Predictor) for search-based domain analysis. The *F. graminearum* genome contains 157 putative WD40 genes, which represent 1.1% of the whole *F. graminearum* protein-coding genes (King *et al*., 2015). Most WD40 genes (152/157) have not been functionally characterized in *F. graminearum*.

Subsequently, we identified 203 WD40 domains comprising a total of 1,254 WD40 repeats. Among the 157 WD40 proteins, 107 contained 6 to 8 repeats, indicating that approximately 68% of these proteins maintain the typical structure of the WD40 protein domain. Additionally, 33 WD40 proteins were found to contain more than 10 repeats, suggesting potential structural or functional divergence within this protein family.

To examine the evolutionary relationships of WD40 proteins in *F. graminearum*, we constructed a phylogenetic tree using the 157 identified WD40 proteins (Fig. 1a). All WD40 protein sequences were subjected to clustering analysis based on sequence similarity. The results showed that in *F. graminearum*, WD40 proteins are divided into 6 major clusters containing 9, 9, 7, 28, 53, and 51 proteins, respectively (Table S2), illustrating the complexity and differentiation of the *F. graminearum* WD40 protein family. Moreover, some chromosomes had a relatively high density of WD40 genes: chromosome 1 contained 61 genes. Chromosomes 1–4 comprised 38.85%, 17.19%, 21.01%, and 22.92% WD40 genes, respectively (Fig. 1b, c).

**Fig. 1.**
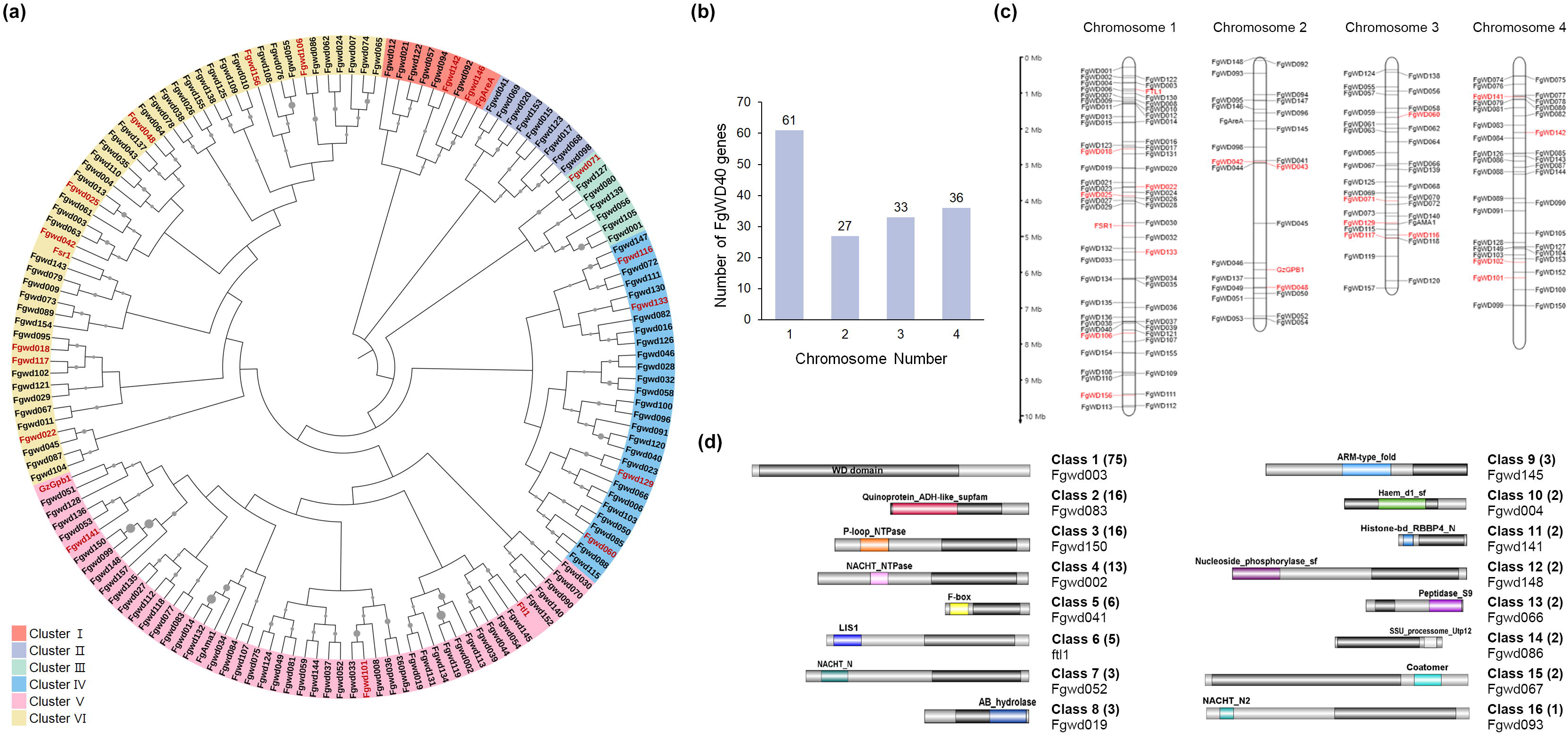
Genome-wide distribution and classification of WD40 genes in *Fusarium graminearum*. **(a)** Phylogenetic classification of 157 WD40 proteins in *F*. *graminearum*. WD40 proteins were grouped into six clusters (clusters I–VI), indicated by different colors. Core WD40 proteins, defined in this study as those involved in both pathogenicity and sexual reproduction, are highlighted in red. An unrooted phylogenetic tree was constructed using the neighbor-joining method in MEGA11 with 1,000 bootstrap replicates. Gray circles at the nodes indicate bootstrap support, with circle size proportional to the bootstrap value. **(b)** Number of WD40 genes on each chromosome of *F*. *graminearum*. **(c)** Chromosomal distribution of WD40 genes in *F. graminearum*. The chromosome map was generated using MapGene2Chrom (MG2C), with chromosome numbers indicated at the top. Core WD40 genes are highlighted in red. **(d)** Schematic representation of the domain architecture of *F. graminearum* WD40 proteins, classified into 16 groups. One representative protein from each group is shown. WD40 domains are shown as black boxes, and additional domains are indicated by different colors and labeled accordingly. The figure was generated using IBS (Illustrator for Biological Sequences).

Based on their domain architectures, the WD40 genes were classified into 16 distinct classes (Fig. 1d). Class 1 comprised 75 WD40 proteins that contained only WD40 domains. The remaining proteins, which harbored additional domains, were grouped into Classes 2 through 16. For example, 16 WD40 proteins with the quinoprotein alcohol dehydrogenase-like superfamily domain were categorized as Class 2, while 13 proteins containing the NACHT nucleoside triphosphatase domain were placed in Class 4. Domains that occurred only once, with the exception of the NWD2 domain, were not assigned to specific classes, as genes with such unique domain architectures were not grouped into separate classes. The domain-based classification closely aligned with the clustering of WD40 proteins, reflecting a similar pattern in the organization of classes.

### Functional enrichment analysis

To explore the potential function of *F. graminearum* WD40 proteins, we performed Gene Ontology (GO) distribution analysis. GO analysis classified WD40 genes into three categories: molecular function, cellular component, and biological process (Table S3) and WD40 were extensively distributed across all three categories (Fig. S1a). Particularly, many WD40 proteins were predicted to be involved in binding (molecular function), cell and organelle (cellular component), and cellular and metabolic processes (biological process) (Table S3).

The WD40 proteins of *F. graminearum* were also classified using Kyoto Encyclopedia of Genes and Genomes (KEGG) pathway enrichment analysis. In total, 18 KEGG pathways were enriched for the selected WD40 proteins (Table S4 and Fig. S1b). Ubiquitin-mediated proteolysis (fgr04120), ribosome biogenesis in eukaryotes (fgr03008), RNA transport (fgr03013), and autophagy (fgr04138) were significantly enriched. The results of functional enrichment analyses predicted that the WD40 proteins of *F. graminearum* participate in several biological processes and interact with multiple interaction partners. Furthermore, these findings are consistent with the well-established functions of the WD40 protein family and provide a foundation for further characterization of conserved genes in *F. graminearum*.

### Construction of the WD40 gene deletion mutant library

To establish a resource for systematic functional characterization of WD40 genes, we generated a WD40 gene deletion mutant library. Among the identified 157 WD40 genes, five genes were excluded due to our inability to construct amplicons. In total, we successfully obtained deletion mutants for 119 WD40 genes, with at least two independent strains for each gene (Fig. S2). Despite more than four independent deletion attempts, knockout mutants could not be obtained for 33 WD40 genes, suggesting that these genes are putatively essential for viability or normal growth. Among the 33 genes, 29 were homologous to WD40 genes reported to be essential for growth in *S. cerevisiae* (Table S2).

### Phenotypic profiling of WD40 gene deletion mutants

We analyzed phenotypic differences between the 119 WD40 gene deletion mutants and the wild-type strain under various developmental and stress conditions. The phenotypic traits examined included vegetative growth, asexual and sexual development, virulence, production of the mycotoxins deoxynivalenol (DON) and zearalenone (ZEA).

Vegetative growth was examined by cultivating WD40 deletion mutants on CM. Based on their relative growth rates compared with the wild-type strain, the mutants were classified into six growth groups (Fig. 2a): group 6, increased growth compared with the wild-type strain; group 5, 81–100%; group 4, 61–80%; group 3, 41–60%; group 2, 21–40%; and group 1, 0–20% of the wild-type growth rate. A total of 27 mutants exhibited reduced vegetative growth and were classified into groups 1–4, whereas 85 mutants belonged to group 5 and displayed growth comparable to the wild-type strain. Representative mutants displaying altered vegetative growth phenotypes (groups 1–4 and 6) are shown in Fig. 2b. In addition, five WD40 deletion mutants exhibited abnormal hyphal morphology, including bulbous and irregular hyphal tips (Fig. 2c).

**Fig. 2.**
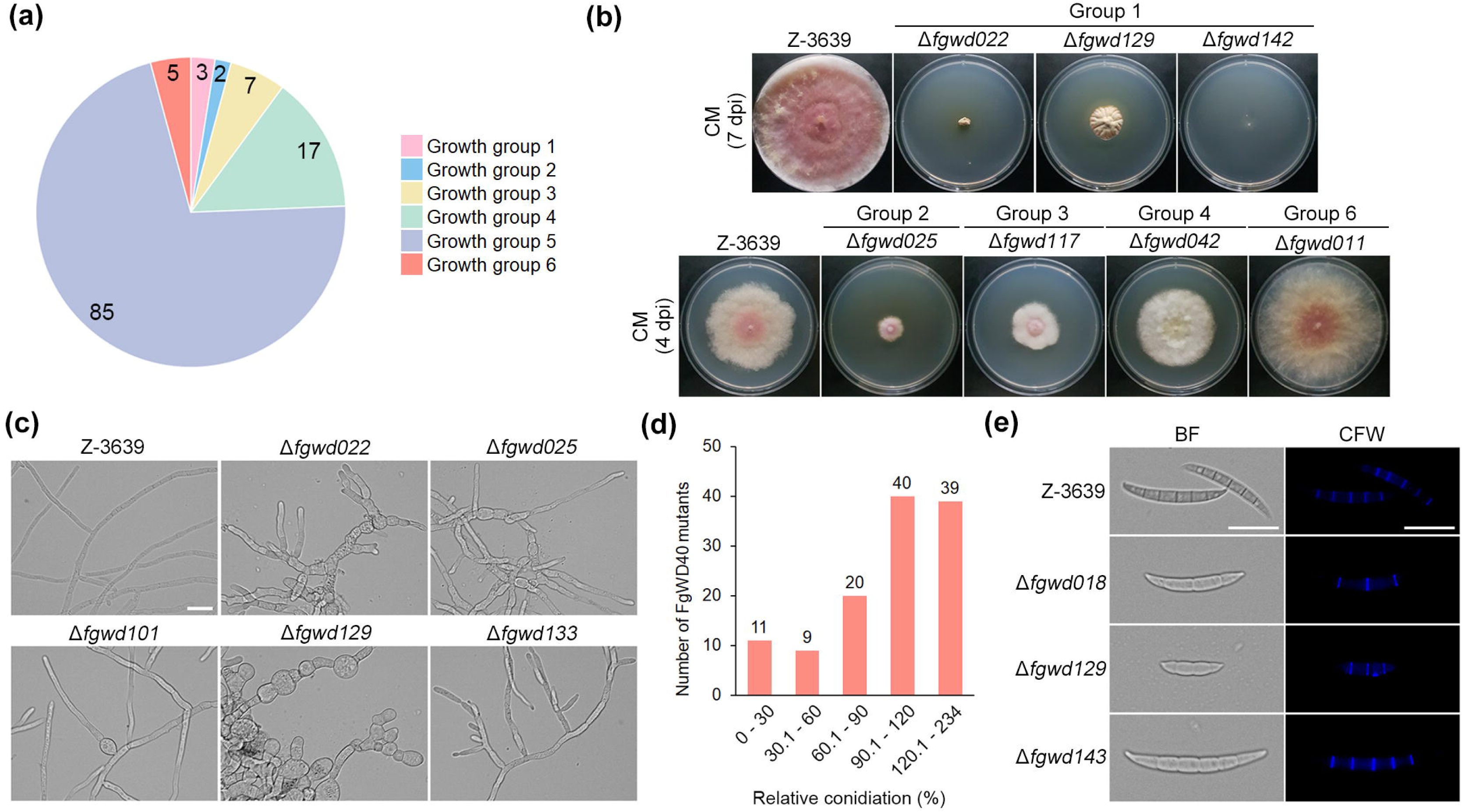
Vegetative growth and conidiation phenotypes of WD40 deletion mutants in *F. graminearum*. **(a)** Classification of WD40 gene deletion mutants into six vegetative growth groups based on their relative growth rates on complete medium (CM) compared with the wild-type strain. **(b)** Representative WD40 gene deletion mutants exhibiting altered vegetative growth phenotypes (groups 1–4 and 6) on CM. Images of group 1 mutants were captured at 7 days post-inoculation (dpi), whereas those of groups 2–4 and 6 were captured at 4 dpi. **(c)** Mycelial morphology of WD40 gene deletion mutants exhibiting abnormal hyphal morphology. Scale bar = 20 µm. **(d)** Number of WD40 gene deletion mutants in each conidiation group based on their relative conidial production compared with the wild-type strain. **(e)** Conidial morphology of WD40 gene deletion mutants exhibiting altered conidial morphology. Bright-field (BF) and calcofluor white (CFW)-stained images were captured after 3 days of incubation in carboxymethyl cellulose medium. Scale bar = 20 µm.

Conidial production was assessed to investigate the role of WD40 genes in asexual development. Based on relative conidiation levels compared with the wild-type strain, WD40 deletion mutants were classified into five groups (Fig. 2d). The conidiation groups were defined according to relative conidial production, ranging from group 1 (0–30%) to group 5 (120.1–234%), with intermediate groups representing 30.1–60% (group 2), 60.1–90% (group 3), and 90.1–120% (group 4) of wild-type conidiation. Based on this classification, 40 mutants were classified into group 4, which included the wild-type conidiation level range, whereas 39 mutants exhibited increased conidial production and were classified into group 5. The remaining mutants showed various degrees of reduced conidial production, with 20, 9, and 11 mutants classified into groups 3, 2, and 1, respectively. In addition to changes in conidial production, several WD40 deletion mutants displayed altered conidial morphology. The Δ*fgwd018* and Δ*fgwd129* mutants produced shorter conidia with fewer septa (2–3 septa), whereas Δ*fgwd143* produced longer conidia compared with the wild-type strain, which typically produced conidia with 5–6 septa (Fig. 2e).

Sexual reproduction was evaluated by examining perithecium formation, internal maturation, ascospore morphology, and forcible ascospore discharge. Based on these phenotypes, WD40 deletion mutants were classified into four sexual development (SD) groups (Fig. 3a). A total of 85 mutants showed normal sexual development (SD group 1), whereas 34 mutants exhibited defects in sexual reproduction. In SD group 2, seven WD40 mutants produced normal-shaped perithecia containing asci and ascospores but displayed defects in ascospore morphology and/or forcible ascospore discharge (Fig. 3b and Fig. S3a). Among them, Δ*fgwd094* and Δ*fgwd139* formed normal melanized perithecia with eight ascospores per ascus, but showed impaired ascospore discharge. Δ*fgwd044* produced abnormal ascospores with fewer septa compared with the wild-type strain, whereas Δ*Fgama1* and Δ*fgwd118* formed irregular shaped ascospores. In addition, Δ*fgwd125* and Δ*fgwd143* produced fewer ascospores per ascus and exhibited defects in ascospore discharge. Five mutants classified into SD group 3 exhibited impaired perithecium maturation and internal development, producing immature perithecia with few or no intact ascospores (Fig. 3c and Fig. S3b). Twenty-two WD40 mutants belonging to SD group 4 failed to produce perithecia, resulting in self-sterility. To determine whether these defects were associated with male or female fertility, self-sterile mutants were outcrossed as either male or female parents with the Δ*mat1-1*/pIGPAPA-GFP strain (Fig. 3d–3f and Fig. S4). Except for Δ*fgwd142*, which was excluded from the fertility analysis due to its severe growth defect, all tested self-sterile mutants retained male fertility, whereas their female fertility varied. Among them, seven mutants retained both male and female fertility, four mutants produced immature perithecia when used as female parents, and ten mutants exhibited complete female sterility with no mature perithecium formation.

**Fig. 3.**
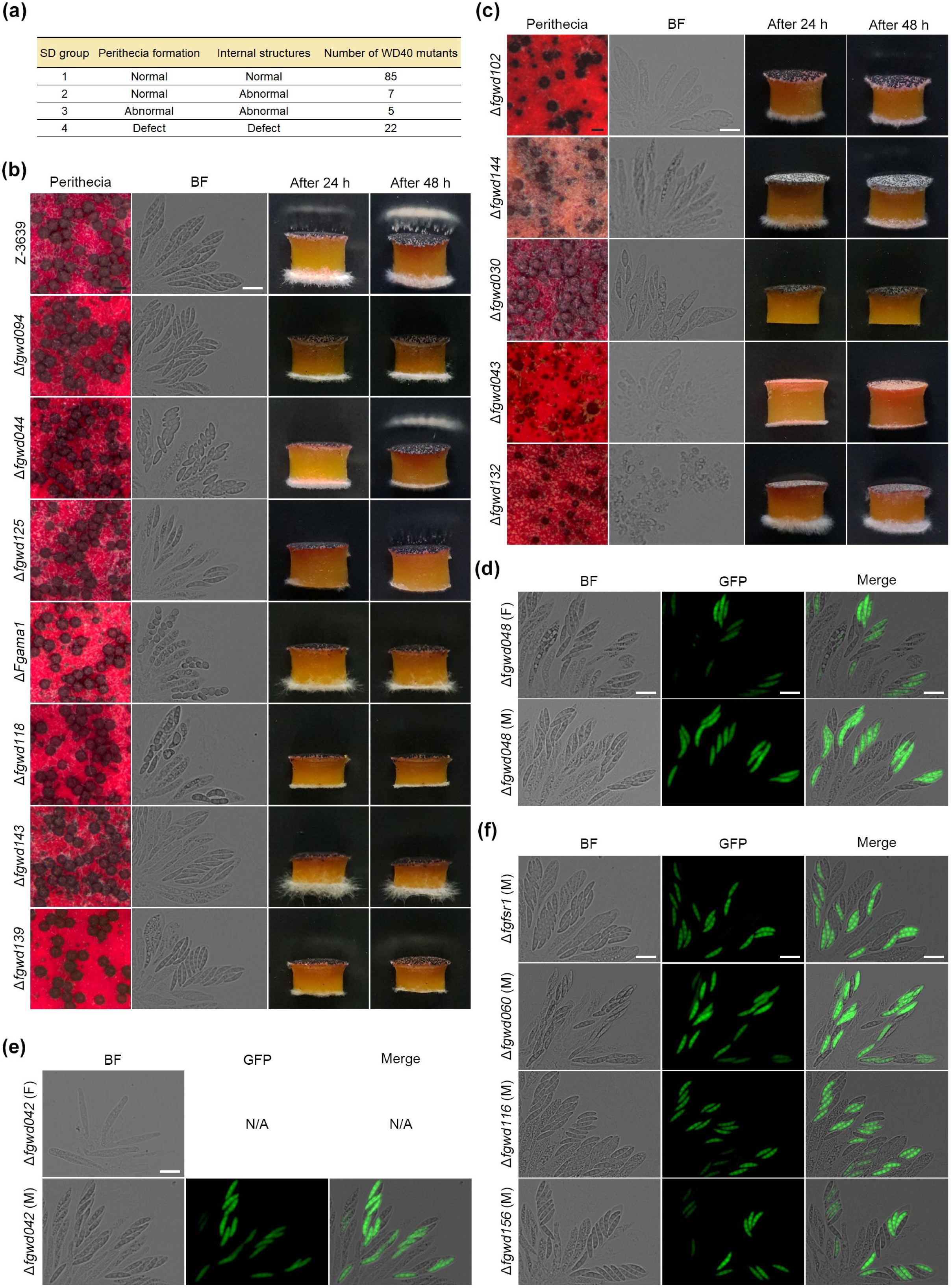
Sexual development phenotypes of WD40 gene deletion mutants in *F. graminearum*. **(a)** Classification of sexual development (SD) groups based on perithecial formation and internal structures. “Normal” indicates a wild-type phenotype. “Abnormal” indicates reduced numbers or altered morphology of perithecia or internal structures, whereas “Defect” indicates the absence of perithecia or internal structures. **(b, c)** Self-fertility of WD40 gene deletion mutants belonging to SD group 2 and 3. Perithecial formation, rosette asci, and forcible ascospore discharge were examined 7–9 days after sexual induction. Forcible ascospore discharge was assessed at 24 and 48 h. White cloudy material represents discharged ascospores. SD group 2 is shown in (b), and SD group 3 is shown in (c). Scale bars: black = 200 µm; white = 20 µm. **(d**–**f)** Representative male and female fertility phenotypes of self-sterile WD40 gene deletion mutants belonging to SD group 4. Each mutant was crossed as either the male (M) or female (F) parent with the Δ*mat1-1*/pIGPAPA- GFP strain. Male and female fertility was evaluated by examining ascus formation and the expected 1:1 segregation of green fluorescent protein (GFP)-positive and GFP-negative ascospores. Representative mutants with normal male and female fertility are shown in (d), those exhibiting impaired female fertility are shown in (e), and those exhibiting female-specific sterility are shown in (f). The remaining mutants are presented in Fig. S4. BF, bright field; N/A, not applicable. Scale bar = 20 µm.

Virulence of WD40 deletion mutants was assessed by point inoculation of wheat heads. Based on disease progression, the mutants were classified into three virulence groups. Overall, 37 mutants exhibited reduced virulence compared with the wild-type strain. Twenty-four mutants in virulence group 1 showed severe defects in disease progression and only colonized the inoculated spikelets, with limited spread to adjacent spikelets (Fig. S5a). In virulence group 2, 13 WD40 mutants were able to spread to adjacent spikelets; however, their virulence was reduced compared with the wild-type strain (Fig. S5b). The remaining 82 mutants were classified into virulence group 3, which showed wild-type level disease progression. Representative disease symptoms of each virulence group are shown in Fig. 4a. Notably, mutants exhibiting significantly reduced vegetative growth (growth groups 1–3) were enriched in virulence group 1, suggesting a close association between vegetative growth and disease progression in *F. graminearum* (Fig. 4b). To further investigate the reduced disease progression observed in WD40 mutants belonging to virulence group 1, penetration ability was first examined using cellophane membrane assay and *in planta* penetration assays (Fig. 4c). Among these mutants, four WD deletion mutants failed to penetrate the cellophane membrane. We further examined whether these penetration defects affected fungal spread during host infection by generating cytosolic GFP-expressing strains of the wild-type strain and the four mutants (Fig. 4d). At 6 days after inoculation, GFP signals from the wild-type strain were observed in adjacent spikelets through rachis nodes, whereas GFP signals from the four mutants were restricted to the inoculated spikelets, indicating impaired fungal spread through rachis nodes *in planta*.

**Fig. 4.**
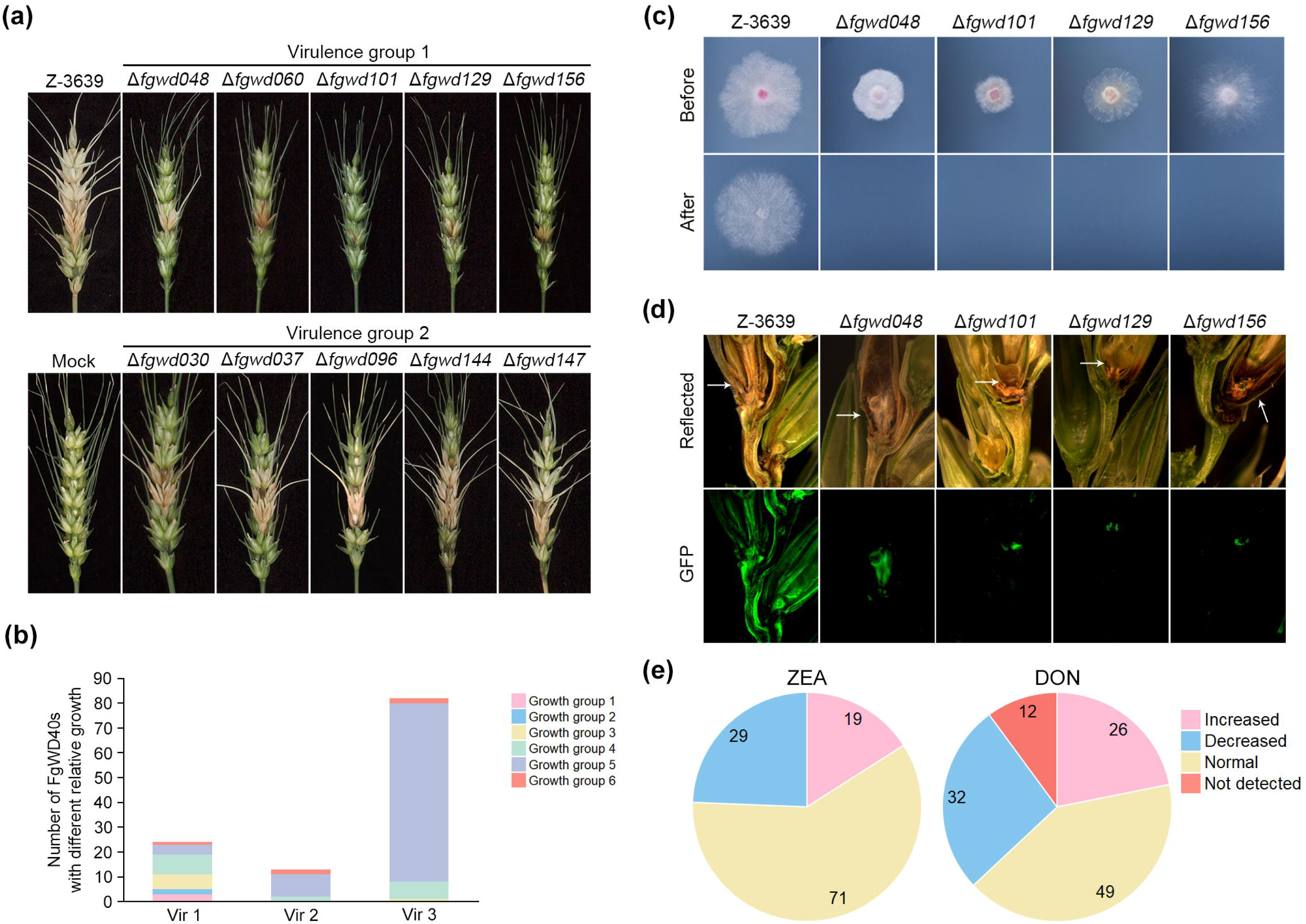
Virulence and mycotoxin production phenotypes of WD40 gene deletion mutants in *F. graminearum*. **(a)** Representative virulence phenotypes of WD40 gene deletion mutants belonging to virulence groups 1 and 2. **(b)** Number of WD40 gene deletion mutants in each virulence group. Colors indicate the vegetative growth groups (groups 1–6) of the corresponding mutants. Vir 1–3 denote virulence groups 1–3. **(c)** Cellophane penetration assay of the wild-type strain and the four WD40 gene deletion mutants that failed to penetrate the cellophane membrane. Colonies were grown on complete medium (CM) overlaid with a cellophane membrane for 2 days (Before). After removal of the membrane, plates were incubated for an additional 2 days (After). **(d)** Longitudinal sections of wheat heads inoculated with cytosolic GFP-expressing wild-type and the four WD40 gene deletion mutants exhibiting defective penetration. Wheat heads were harvested at 6 days after inoculation and longitudinally sectioned. GFP fluorescence indicates fungal colonization and spread from the inoculated spikelet. Arrowheads indicate the inoculated spikelets. “Reflected” indicates reflected-light images. **(e)** Pie charts showing the distribution of WD40 gene deletion mutants according to their relative zearalenone (ZEA) and deoxynivalenol (DON) production. Numbers indicate the number of mutants in each category.

Production of the mycotoxins ZEA and DON was analyzed in WD40 deletion mutants. Altered ZEA production was observed in 48 mutants, with 19 mutants showing increased production and 29 mutants showing reduced production compared with the wild-type strain (Fig. 4e). In addition, 38 mutants exhibited altered DON production, including 26 mutants with increased accumulation and 12 mutants with reduced production. All 12 mutants with reduced DON production also belonged to virulence group 1 and SD group 4.

### Comprehensive phenotypic summary and phenotypic correlation observed among WD40 proteins

In addition to developmental and pathogenicity-related phenotypes, we evaluated the stress responses of WD40 deletion mutants under various environmental and chemical stress conditions. Stress tolerance was assessed under conditions affecting multiple cellular processes, including cell wall integrity, osmotic stress response, oxidative stress response, endoplasmic reticulum stress response, and antifungal drug sensitivity.

Together with the stress response data, we generated a comprehensive genome-wide phenome dataset for 119 non-essential WD40 deletion mutants, covering vegetative growth, asexual and sexual development, virulence, mycotoxin production, stress tolerance, and antifungal drug responses (Table S5). To provide an overview of the phenotypic contribution of WD40 proteins, all phenotypic data were integrated and visualized as a heatmap (Fig. 5). This integrated phenotypic landscape revealed that several WD40 deletion mutants exhibited defects across multiple biological processes. To further investigate relationships among these phenotypic traits, we performed Spearman correlation analysis using the integrated dataset (Fig. S6). The analysis revealed strong associations between vegetative growth and sexual reproduction, as well as between virulence and sexual reproduction (Spearman correlation coefficient > 0.5), indicating that these phenotypic alterations frequently occurred together among WD40 deletion mutants. Because sexual reproduction and virulence represent distinct biological processes in the fungal life cycle, the strong association between these traits suggests that a subset of WD40 proteins contributes to both processes. Based on this observation, we defined WD40 genes required for both sexual reproduction and virulence as core WD40 genes associated with fungal development and pathogenicity.

**Fig. 5.**
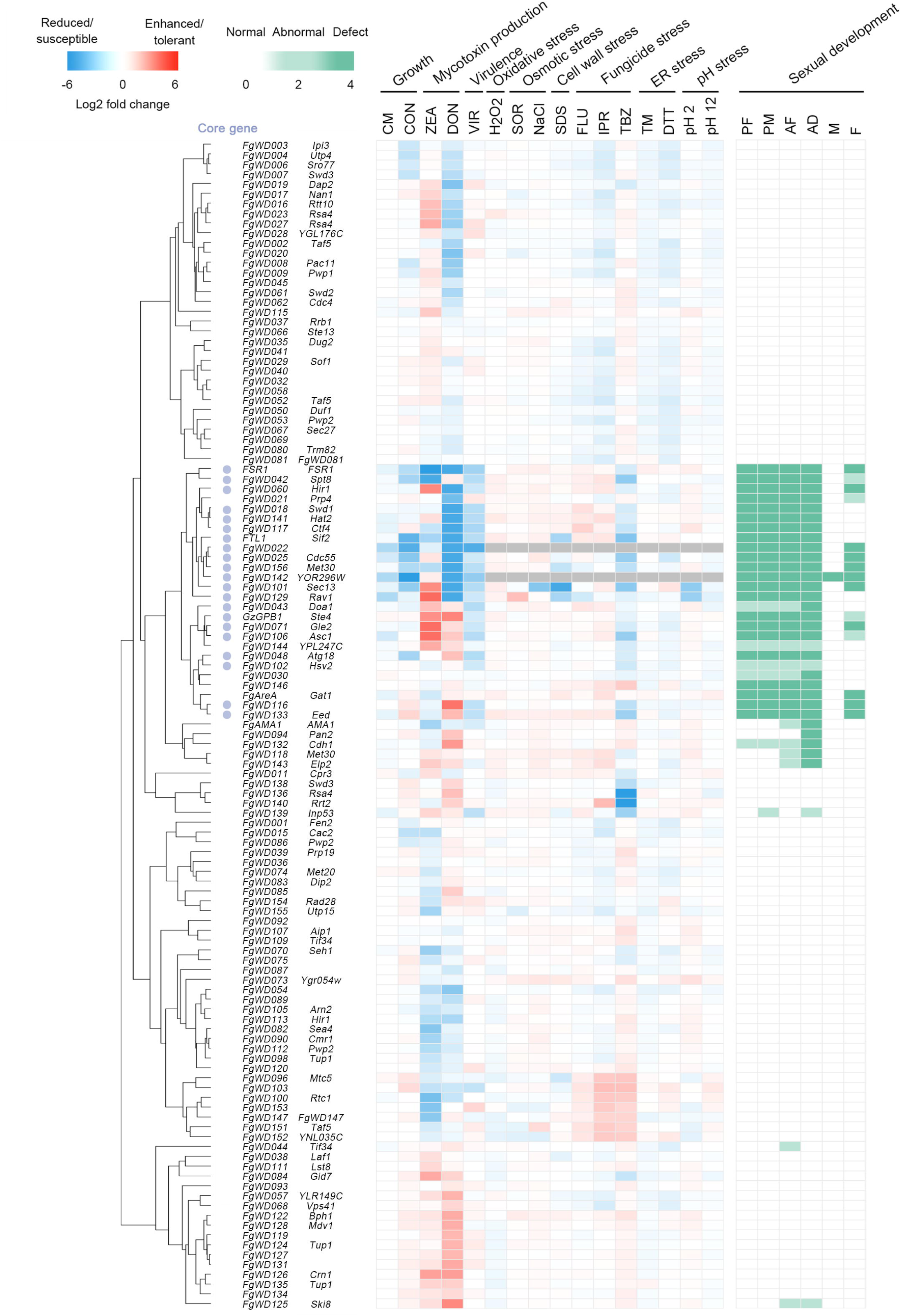
Phenotypic clustering of non-essential WD40 gene deletion mutants in *F. graminearum*. Hierarchical clustering of 119 WD40 gene deletion mutants based on phenotypic profiles was performed using Pearson’s correlation in the Morpheus software. Log_2_-transformed fold changes relative to the wild-type strain were used for hierarchical clustering and heat map generation. Blue dots indicate WD40 genes designated as core genes in this study. In the heat map, blue indicates reduced phenotypes or increased chemical susceptibility, whereas red indicates enhanced phenotypes or increased chemical tolerance. The right panel summarizes the phenotypes associated with sexual development, where white, light green, and dark green indicate wild-type, abnormal, and defective phenotypes, respectively. The complete phenotypic dataset used for hierarchical clustering and heat map generation is provided in Table S5. Abbreviations used in the heat map: CM, complete medium; CON, conidiation; ZEA, zearalenone; DON, deoxynivalenol; VIR, virulence; H_2_O_2_, 0.31 mM hydrogen peroxide; SOR, 1 M sorbitol; NaCl, 1 M NaCl; SDS, 0.1 mg mL^−1^ sodium dodecyl sulfate; FLU, 0.05 μg mL^−1^ fludioxonil; IPR, 125 μg mL^−1^ iprodione; TBZ, 0.625 μg mL^−1^ tebuconazole; TM, 2 μg mL^−1^ tunicamycin; DTT, 1.25 mM dithiothreitol; PF, perithecia formation; PM, perithecia maturation; AF, ascospore formation; AD, ascospore discharge; M, male fertility; F, female fertility.

### Identification of interaction networks associated with core WD40 proteins

To investigate the molecular networks associated with core WD40 proteins, we performed affinity purification-mass spectrometry (AP-MS) analysis. Because WD40 proteins primarily function as scaffolds for protein-protein interactions, identification of their interacting partners is essential for understanding their functional roles. Among the core WD40 proteins associated with fungal development and virulence, Fgwd101 and Fgwd133 were selected as representative candidates because their orthologs, Sec13 and EED, respectively, have been extensively characterized in model organisms. Given the limited information available regarding WD40 protein interaction networks in *F. graminearum*, analyzing conserved WD40 proteins provided a reference framework to validate our interaction-based approach and to investigate conserved and potentially species-specific protein associations.

To perform AP-MS analysis, we generated GFP-tagged complementation and overexpression strains of *FgWD101* and *FgWD133*. For *FgWD101*, immunoprecipitation of Fgwd101-GFP was confirmed by western blot analysis, detecting a protein band corresponding to the expected size of approximately 63 kDa (Fig. S7a). AP-MS analysis identified 33 candidate interacting proteins enriched in the Fgwd101-GFP samples compared to the control (log2 fold change > 1) (Table S6). Among these candidates, Fgwd072 was detected in both *FgWD101* complementation and overexpression strains and showed the highest enrichment frequency. STRING-based network (Szklarczyk *et al*., 2023) predicted a specific interaction between Fgwd101 and Fgwd072 (Fig. 6a), whereas other identified candidates mainly clustered with ribosomal proteins, which were considered background-associated proteins in the AP-MS dataset. The interaction between Fgwd101 and Fgwd072 was further confirmed by co-immunoprecipitation (co-IP) and yeast two-hybrid (Y2H) assay (Fig. 6b, 6c).

**Fig. 6.**
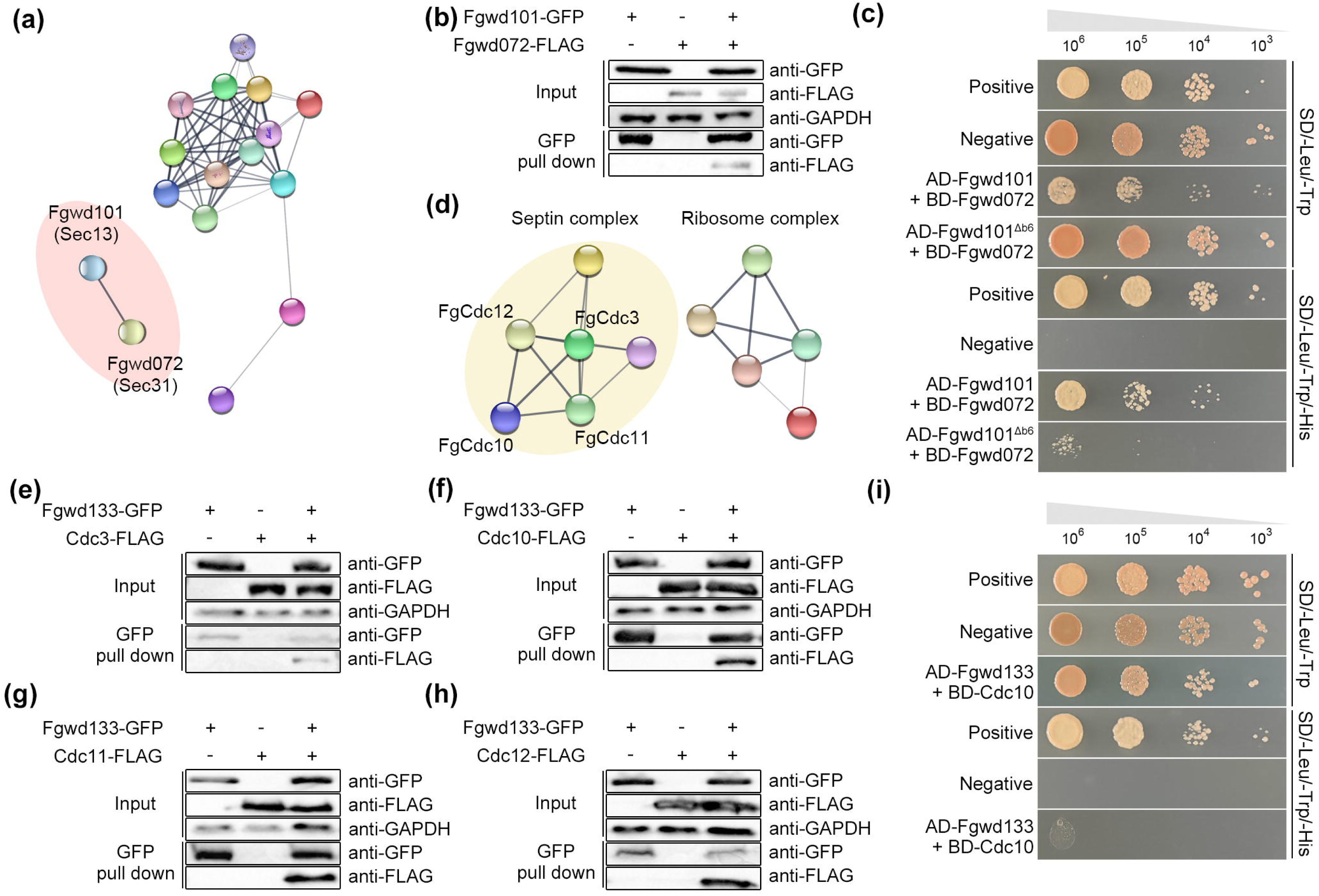
Validation of protein interaction networks associated with Fgwd101 and Fgwd133 in *F. graminearum*. **(a)** STRING network of Fgwd101-associated proteins identified by affinity purification-mass spectrometry (AP-MS). Fgwd072 was selected for interaction validation. **(b)** Co-immunoprecipitation (co-IP) analysis confirming the interaction between Fgwd101 and Fgwd072. Total protein extracts (Input) and protein immunoprecipitated using anti-GFP magnetic beads (GFP pull down) were detected with anti-GFP or anti-FLAG antibodies. GAPDH served as a loading control for the input samples. **(c)** Yeast two-hybrid (Y2H) assay confirming the interaction between Fgwd101 and Fgwd072. **(d)** STRING network of Fgwd133-associated proteins identified by AP-MS. Septin complex proteins were selected for interaction validation. **(e–h)** Co-IP analysis confirming the association between Fgwd133 and septin complex proteins. Total protein extracts (Input) and proteins immunoprecipitated using anti-GFP magnetic beads (GFP pull down) were detected with anti-GFP or anti-FLAG antibodies. GAPDH served as a loading control for the input samples. (e) Cdc3; (f) Cdc10; (g) Cdc11; (h) Cdc12. **(i)** Y2H assay examining the interaction between Fgwd133 and Cdc10. AD, activation domain; BD, DNA-binding domain; SD, synthetic dropout medium.

Using the same approach, we investigated the protein interaction network associated with Fgwd133. Immunoprecipitation of GFP-tagged Fgwd133 was confirmed by western blot analysis, detecting a protein band corresponding to the expected size of approximately 86 kDa (Fig. S7b). AP-MS analysis identified candidate Fgwd133-associated proteins using three independent strains expressing GFP-tagged Fgwd133 (Table S7). STRING-based network analysis showed that Fgwd133-associated proteins were mainly divided into ribosome-associated proteins and septin complex proteins (Fig. 6d). The septin complex cluster included Cdc3, Cdc10, Cdc11, and Cdc12, which were selected for further validation as potential Fgwd133-associated proteins. Co-IP analysis confirmed the association between Fgwd133 and septin proteins (Fig. 6e-6h). However, in the Y2H assay, Fgwd133 exhibited only a weak direct interaction with Cdc10, which became detectable after one additional day of incubation (Fig. 6i, S7c). No direct interactions were observed with the other septin proteins (Fig. S7d). These results suggest that Fgwd133 weakly and directly interacts with Cdc10, whereas its association with the other septin components is likely mediated through the septin protein complex rather than through individual direct interactions.

### WD40 domains are required for the biological functions of core WD40 proteins

We next investigated whether WD40 domain-mediated protein interactions contribute to the biological functions of core WD40 proteins. To this end, we generated strains expressing proteins lacking specific WD40 blades in the corresponding deletion mutant backgrounds. To select WD40 blades likely to play important roles in WD40-mediated protein-protein interactions, we used previously identified interacting proteins to guide AlphaFold prediction of interaction interfaces.

For Fgwd101, AlphaFold structural prediction identified blade 6 as part of the predicted interaction interface with the previously identified interacting protein Fgwd072 (Fig. 7a). We therefore generated a strain expressing a blade 6-deleted version of *FgWD101* (*FgWD101*^Δb6^) and first examined whether deletion of this blade affected its interaction with Fgwd072. Co-IP analysis showed that the association between *FgWD101*^Δb6^ and Fgwd072 was no longer detectable, and Y2H analysis further demonstrated a marked reduction in the direct interaction between Fgwd101^Δb6^ and Fgwd072 compared with the full-length Fgwd101 protein (Fig. 7b, 6c). The ability of Fgwd101^Δb6^ to complement the defects of the Δ*fgwd101* mutant was then compared with that of the full-length complementation strain (*FgWD101*c). The *FgWD101*c strain substantially complemented defects in vegetative growth and virulence and restored sexual reproduction compared with the Δ*fgwd101* mutant (Fig. 7c, 7d). In contrast, the *FgWD101*^Δb6^ strain failed to complement defects in vegetative growth, sexual reproduction, or virulence and exhibited phenotypes indistinguishable from those of the Δ*fgwd101* mutant.

**Fig. 7.**
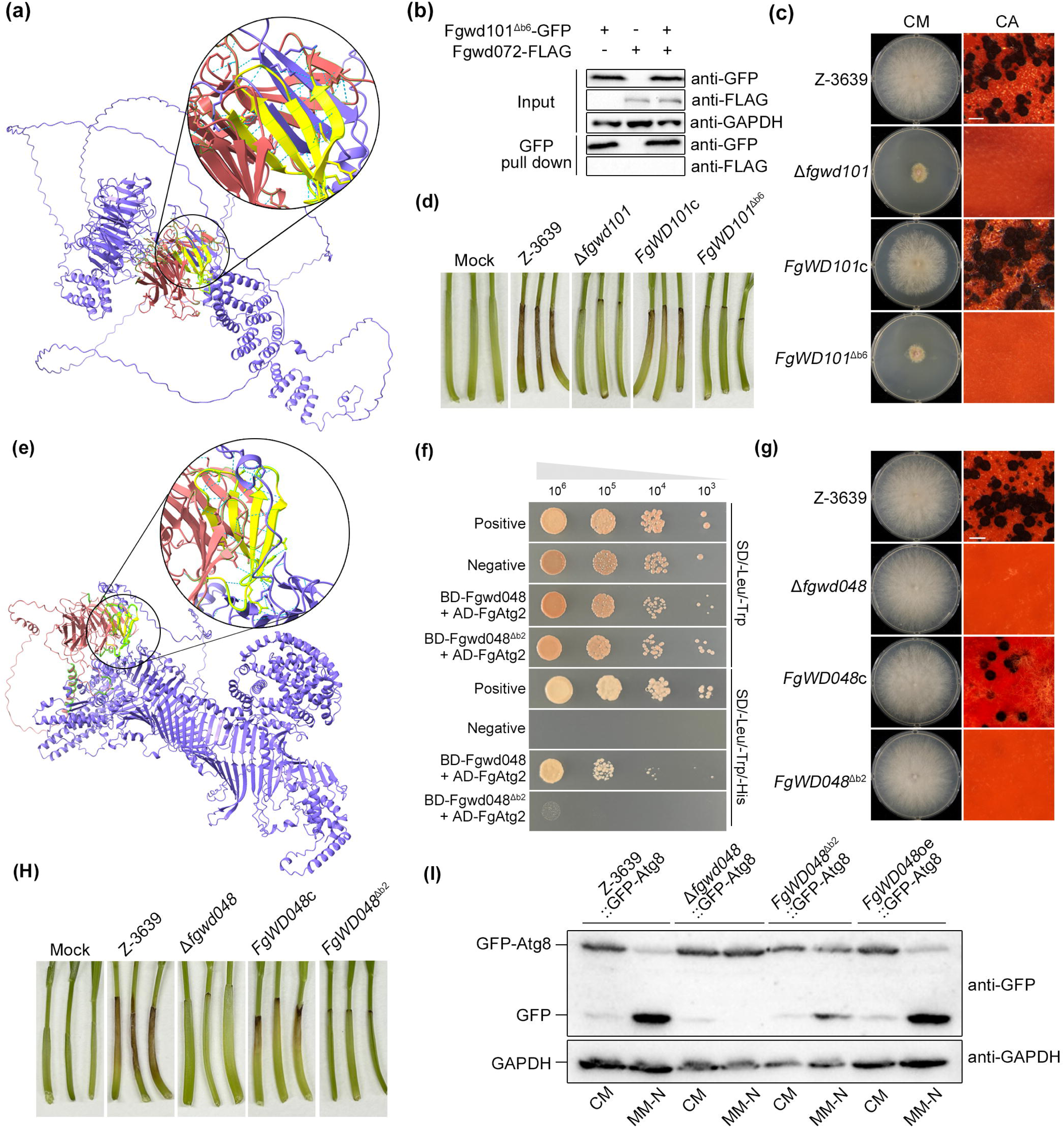
Functional analysis of predicted interaction-associated WD40 blades in Fgwd101 and Fgwd048 of *F. graminearum*. **(a)** AlphaFold-predicted interaction interface between Fgwd101 (red) and Fgwd072 (purple). The predicted interaction-associated WD40 blade (blade 6) of Fgwd101 is highlighted in yellow. **(b)** Co-immunoprecipitation (co-IP) analysis examining the association between *FgWD101*^Δb6^ and Fgwd072. Total protein extracts (Input) and proteins immunoprecipitated using anti-GFP magnetic beads (GFP pull down) were detected with anti-GFP or anti-FLAG antibodies. GAPDH served as a loading control for the input samples. **(c)** Vegetative growth and sexual reproduction of the wild-type strain (Z-3639), Δ*fgwd101*, *FgWD101*c, and *FgWD101*^Δb6^. CM, complete medium; CA, carrot agar. Scale bar = 200 µm. **(d)** Virulence assay of the indicated strains using wheat coleoptiles. **(e)** AlphaFold-predicted interaction interface between Fgwd048 (red) and Atg2 (purple). The predicted interaction-associated WD40 blade (blade 2) of Fgwd048 is highlighted in yellow. **(f)** Yeast two-hybrid (Y2H) assay examining the interaction between Fgwd048 and Atg2. AD, activation domain; BD, DNA-binding domain. **(g)** Vegetative growth and sexual reproduction of the wild-type strain, Δ*fgwd048*, *FgWD048*c, and *FgWD048*^Δb2^. CM, complete medium; CA, carrot agar. Scale bar = 200 µm. **(h)** Virulence assay of the indicated strains using wheat coleoptiles. **(i)** Immunoblot analysis of GFP-Atg8 processing in the indicated strains to evaluate autophagic activity. GFP-Atg8 and free GFP were detected using an anti-GFP antibody, and GAPDH served as a loading control. MM-N, nitrogen-free minimal medium.

We also attempted to generate a blade-deleted *FgWD133* strain by introducing a construct lacking the WD40 blade predicted to contribute to the interaction interface with Cdc10. However, transformants carrying the blade-deleted construct could not be obtained. Therefore, we selected Fgwd048, another core WD40 protein previously characterized as an autophagy-related protein in *F. graminearum*, to further investigate the contribution of WD40 domains to protein function. Based on the previously reported interaction between Fgwd048 and Atg2 (Obara *et al*., 2008), we selected blade 2 for functional analysis. AlphaFold structural prediction identified blade 2 as part of the predicted interaction interface between Fgwd048 and Atg2 (Fig. 7e). We therefore generated a strain expressing a blade 2-deleted version of Fgwd048 (*FgWD048*^Δb2^). Y2H analysis showed that deletion of blade 2 markedly weakened the interaction between Fgwd048 and Atg2 (Fig. 7f). We next examined whether *FgWD048*^Δb2^ retained the biological functions of Fgwd048. Compared with the full-length complementation strain (*FgWD048*c), *FgWD048*^Δb2^ failed to rescue defects in sexual reproduction and virulence, exhibiting phenotypes comparable to those of the Δ*fgwd048* mutant (Fig. 7g, 7h). Since Fgwd048 has previously been shown to function in autophagy through Atg8 processing, we further investigated whether deletion of blade 2 affected this activity. GFP-Atg8 was expressed in each strain, and Atg8 processing was evaluated by immunoblot analysis following nitrogen starvation. In the full length *FgWD048*-expressing strains, transfer to MM-N for 2 h resulted in efficient Atg8 processing, as indicated by the accumulation of free GFP. In contrast, GFP-Atg8 processing was markedly reduced in *FgWD048*^Δb2^, although not completely abolished (Fig. 7i).

Together, these findings indicate that disruption of the predicted interaction blades compromises both WD40-mediated protein-protein interactions and the biological functions of the examined WD40 proteins.

## Discussion

WD40 domain-containing proteins participate in diverse biological processes through protein-protein interactions, but the functional contribution of individual WD40 proteins remains largely unexplored in filamentous fungi. In this study, we identified 157 putative WD40 genes in *F. graminearum* and generated a deletion mutant library containing 119 non-essential WD40 genes. Comprehensive phenotypic profiling under 22 different conditions, including fungal development, pathogenicity, mycotoxin production, and various stress responses, allowed us to establish a genome-wide WD40 phenome dataset. This systematic resource provides a foundation for understanding the functional diversity of WD40 proteins and exploring WD40-mediated regulatory networks in *F. graminearum*.

Among the 33 putative essential WD40 genes identified in this study, 29 had essential orthologs in *S. cerevisiae*, suggesting that a subset of WD40 proteins retains evolutionarily conserved functions required for fundamental cellular processes. However, this conservation was not observed for all WD40 proteins, as several orthologs of essential WD40 genes in *S. cerevisiae* were successfully deleted in *F. graminearum,* with some mutants displaying alterations in specific biological processes. These observations suggest that, although some WD40 proteins maintain conserved roles across fungal species, others may have acquired species-specific functions. Given that WD40 proteins function as scaffolds for protein-protein interactions, these differences may be associated with changes in their regulatory roles or interaction networks.

In addition to characterizing the contribution of individual WD40 proteins, the genome-wide phenome dataset allowed us to investigate relationships among phenotypic traits affected by WD40 gene disruption. Spearman correlation analysis of the WD40 deletion mutant phenome revealed a relatively strong positive correlation between sexual reproduction and virulence. Sexual reproduction and pathogenicity represent distinct stages of the *F. graminearum* life cycle and involve different developmental processes (Guenther & Trail, 2005; Leplat *et al*., 2013). Sexual reproduction occurs primarily on plant residues and involves the differentiation of specialized reproductive structures, including perithecium development and ascospore formation, which contribute to survival and inoculum production. In contrast, pathogenicity occurs during host infection and relies on invasive hyphal growth and colonization within host tissues. Interestingly, a positive correlation between these distinct biological processes has also been observed in a genome-wide transcription factor mutant library of *F. graminearum*, in which defects in virulence were strongly correlated with impaired perithecium formation and maturation (Son *et al*., 2011b). These observations suggest that the correlation between sexual reproduction and virulence is not specific to WD40 proteins but may reflect the presence of shared genetic components involved in both biological processes. Further investigation of these shared regulators may provide deeper insights into the mechanisms coordinating fungal development and pathogenicity in *F. graminearum*.

Beyond phenotypic characterization of WD40 proteins, we further investigated selected WD40 domain-containing proteins, Fgwd101, Fgwd133, and Fgwd048, to explore their roles as protein-protein interaction platforms and understand how WD40-mediated interaction networks contribute to their biological functions. AP-MS analysis identified Fgwd072 as a major interacting partner of Fgwd101. Fgwd101 and Fgwd072 are orthologs of *S. cerevisiae* Sec13 and Sec31, respectively, which form a conserved protein complex through direct interaction as components of the COPII vesicle coat (Novick *et al*., 1980; Salama *et al*., 1997; Nie & Ma, 2025). The identification of the Fgwd101-Fgwd072 interaction suggests that this conserved Sec13-Sec31 interaction module is maintained in *F. graminearum*. Interestingly, although *FgWD072* appears to be essential in *F. graminearum*, as deletion mutants could not be obtained, disruption of the Fgwd101-Fgwd072 interaction resulted in developmental defects rather than loss of viability. This observation suggests that the essential functions of Fgwd072 are not solely dependent on its interaction with Fgwd101. Instead, the Fgwd101-Fgwd072 interaction likely contributes to a subset of COPII-associated processes required for fungal development and pathogenicity, whereas other interaction partners or functions of Fgwd072 may be sufficient to support essential cellular processes.

In contrast to Fgwd101, whose conserved interaction with its orthologous partner was confirmed in this study, Fgwd133 exhibited a previously unrecognized interaction profile. Fgwd133, an ortholog of the WD40 repeat-containing Eed protein, has been reported as a component of the Polycomb Repressive Complex 2 (PRC2)(Connolly *et al*., 2013). Likewise, Kmt6, Suz12, and Eed have been shown to form a PRC2 complex in *F. graminearum* (Tang *et al*., 2021). However, our AP-MS analysis identified components of the septin complex rather than previously reported PRC2-associated proteins as major interaction partners of Fgwd133. Because the interaction between Fgwd133 and Cdc10 was further supported by both Co-IP and Y2H analyses, these findings are unlikely to reflect experimental artifacts. Instead, they likely reflect the context-dependent nature of protein interaction networks, which are influenced by developmental stage, cellular status, and environmental conditions. These observations further suggest that Fgwd133 may participate in distinct protein complexes depending on the biological context.

We further investigated whether the WD40 domain itself contributes to the observed phenotypes and biological functions of WD40 proteins using Fgwd101 and Fgwd048 as representative examples. Based on experimentally identified or previously reported protein interactions, we selected WD40 blades predicted to play important roles in protein-protein interactions. In both proteins, deletion of the selected blade markedly weakened protein interactions and abolished recovery of the corresponding mutant phenotypes, demonstrating that the WD40 domain contributes substantially to the biological functions of these proteins. However, the observed defects cannot be attributed solely to disruption of the representative protein interactions used to identify the candidate blades. Because WD40 proteins function as molecular scaffolds that coordinate interactions with multiple binding partners, disruption of a single interaction-associated blade is likely to affect additional protein interactions beyond those examined in this study. Although the specific protein interactions underlying the observed phenotypes remain to be identified, our findings support an important role for WD40-mediated protein interactions in the biological functions of WD40 proteins.

Interestingly, although deletion of the predicted interaction-associated blades markedly reduced the representative protein interactions in both Fgwd101 and Fgwd048, weak interactions between each WD40 protein and its representative interacting partner remained detectable. Nevertheless, neither blade-deletion strain restored the biological functions of the corresponding deletion mutant. These observations suggest that individual WD40 blades make major contributions to protein-protein interactions but are unlikely to function as independent interaction modules. Instead, stable protein interactions are likely maintained through the cooperative architecture of the WD40 β-propeller, in which multiple blades collectively contribute to interaction interfaces.

Collectively, our findings reveal the functional diversity of WD40 proteins in *F. graminearum* and highlight their potential roles as versatile platforms for protein complex formation. By combining genome-wide phenotypic profiling with protein interaction analysis and functional validation of WD40 interaction-associated blades, this study expands our understanding of how WD40 proteins contribute to diverse biological processes in filamentous fungi.

## Acknowledgments

This study was supported by the Korea Institute of Planning and Evaluation for Technology in Food, Agriculture, and Forestry (IPET) through the Agriculture and Food Convergence Technologies Program for Research Manpower Development, funded by the Ministry of Agriculture, Food and Rural Affairs (MAFRA) (RS-2024-00398300), the National Research Foundation of Korea (NRF) (RS-2025-00553624), and the intramural grant from the Korea Institute of Science and Technology (KIST) (Grand Challenge, 26Z0001), Republic of Korea.

## Author contributions

SC, NL, Y-WL, D-GS and HS conceived the study and designed the experiment; SC, NL, JP, HJ, SK, J-EK, JS, HM, KM, YC, AH, HK and GJC performed experiments and collected the data; SC, NL, JP and HJ analyzed the data; SC, NL, JP, HJ, D-GS and HS wrote the manuscript. SC and NL contributed equally to this work.

## Data availability

All data supporting the findings of this study are included in this article and its Supporting Information (Figs S1–S7; Tables S1–S7).

## Supporting Information Legends

**Table S1.** List of oligonucleotides used in this study

**Table S2.** List of 157 WD40 genes in *F. graminearum*

**Table S3.** Gene Ontology distribution of WD40 genes of *F. graminearum*

**Table S4.** Kyoto Encyclopedia of Genes and Genomes pathway enrichment of WD40 proteins in *F. graminearum*

**Table S5.** Phenotypes of 119 WD40 deletion mutants in *F. graminearum*.

**Table S6.** The result of affinity purification-mass spectrometry for Fgwd101.

**Table S7.** The result of affinity purification-mass spectrometry for Fgwd133.

**Fig. S1.** Gene Ontology classification and Kyoto Encyclopedia of Genes and Genomes pathway enrichment analysis of WD40 proteins in *F. graminearum*.

**Fig. S2.** Southern blot validation of WD40 gene deletion mutants and the GFP-Atg8 strain in *F. graminearum*.

**Fig. S3.** Perithecial cirrhus formation on carrot agar medium at 9 days post-inoculation.

**Fig. S4.** Assessment of male and female fertility in WD40 gene deletion mutants of *F. graminearum*.

**Fig. S5.** Virulence of *F. graminearum* WD40 deletion mutants on wheat heads.

**Fig. S6.** Phenotypic correlation analysis of *F. graminearum* WD40 gene deletion mutants.

**Fig. S7.** Immunoprecipitation of GFP-tagged Fgwd101 and Fgwd133 for affinity purification-mass spectrometry analysis and yeast two-hybrid assays examining interactions between Fgwd133 and septin complex proteins.

## References

Bowden RL, Leslie JF. 1999. Sexual recombination in *Gibberella zeae*. Phytopathology 89(2): 182–188.

Chao J, Li Z, Sun Y, Aluko OO, Wu X, Wang Q, Liu G. 2021. MG2C: a user-friendly online tool for drawing genetic maps. Molecular Horticulture 1(1): 16.

Choi S, Yang JW, Kim JE, Jeon H, Shin S, Wui D, Kim LS, Kim BJ, Son H, Min K. 2023. Infectivity and stress tolerance traits affect community assembly of plant pathogenic fungi. Frontiers in Microbiology 14: 1234724.

Connolly LR, Smith KM, Freitag M. 2013. The *Fusarium graminearum* histone H3 K27 methyltransferase KMT6 regulates development and expression of secondary metabolite gene clusters. PLoS Genetics 9(10): e1003916.

Desjardins AE. 2006. Fusarium mycotoxins : chemistry, genetics, and biology. St. Paul, MN: APS Press, American Phytopathological Society.

Desjardins AE, Proctor RH. 2007. Molecular biology of *Fusarium* mycotoxins. Int. J. Food Microbiol. 119(1): 47–50.

Edgar RC. 2004. MUSCLE: multiple sequence alignment with high accuracy and high throughput. Nucleic Acids Research 32(5): 1792–1797.

Evans R, O’Neill M, Pritzel A, Antropova N, Senior A, Green T, Žídek A, Bates R, Blackwell S, Yim J et al. 2022.Protein complex prediction with AlphaFold-Multimer. bioRxiv: doi: 10.1101/2021.1110.1104.463034.

Goswami RS, Kistler HC. 2004. Heading for disaster: *Fusarium graminearum* on cereal crops. Molecular Plant Pathology 5(6): 515–525.

Götz S, García-Gómez JM, Terol J, Williams TD, Nagaraj SH, Nueda MJ, Robles M, Talón M, Dopazo J, Conesa A. 2008. High-throughput functional annotation and data mining with the Blast2GO suite. Nucleic Acids Research 36(10): 3420–3435.

Guenther JC, Trail F. 2005. The development and differentiation of *Gibberella zeae* (anamorph: *Fusarium graminearum*) during colonization of wheat. Mycologia 97(1): 229– 237.

Hu R, Xiao J, Gu T, Yu X, Zhang Y, Chang J, Yang G, He G. 2018. Genome-wide identification and analysis of WD40 proteins in wheat (*Triticum aestivum* L.). BMC Genomics 19(1): 803.

Horwitz BA, Sharon A, Lu S-W, Ritter V, Sandrock TM, Yoder OC, Turgeon BG. 1999. A G protein alpha subunit from *Cochliobolus heterostrophus* involved in mating and appressorium formation. Fungal Genetics Biology 26(1): 19–32.

Kim YJ, Yoo JE, Jeon Y, Chong JU, Choi GH, Song DG, Jung SH, Oh BK, Park YN. 2018. Suppression of PROX1-mediated TERT expression in hepatitis B viral hepatocellular carcinoma. International Journal of Cancer 143(12): 3155–3168.

Kim YT, Lee YR, Jin J, Han KH, Kim H, Kim JC, Lee T, Yun SH, Lee YW. 2005. Two different polyketide synthase genes are required for synthesis of zearalenone in *Gibberella zeae*. Molecular Microbiology 58(4): 1102–1113.

King R, Urban M, Hammond-Kosack MC, Hassani-Pak K, Hammond-Kosack KE. 2015. The completed genome sequence of the pathogenic ascomycete fungus *Fusarium graminearum*. BMC Genomics 16(1): 544.

Lambright DG, Sondek J, Bohm A, Skiba NP, Hamm HE, Sigler PB. 1996. The 2.0 Å crystal structure of a heterotrimeric G protein. Nature 379(6563): 311–319.

Lee J, Lee T, Lee YW, Yun SH, Turgeon BG. 2003. Shifting fungal reproductive mode by manipulation of mating type genes: obligatory heterothallism of *Gibberella zeae*. Molecular Microbiology 50(1): 145–152.

Leplat J, Friberg H, Abid M, Steinberg C. 2013. Survival of *Fusarium graminearum*, the causal agent of Fusarium head blight. A review. Agronomy for Sustainable Development 33(1): 97–111.

Leslie JF, Summerell BA. 2006. The Fusarium laboratory manual. Ames, Iowa: Blackwell Publighing.

Letunic I, Bork P. 2019. Interactive Tree Of Life (iTOL) v4: recent updates and new developments. Nucleic Acids Research 47(W1): W256–W259.

Meng EC, Goddard TD, Pettersen EF, Couch GS, Pearson ZJ, Morris JH, Ferrin TE. 2023. UCSF ChimeraX: Tools for structure building and analysis. Protein Science 32(11): e4792.

Mirdita M, Schütze K, Moriwaki Y, Heo L, Ovchinnikov S, Steinegger M. 2022. ColabFold: making protein folding accessible to all. Nature Methods 19(6): 679–682.

Neer EJ, Schmidt CJ, Nambudripad R, Smith TF. 1994. The ancient regulatory-protein family of WD-repeat proteins. Nature 371(6495): 297–300.

Nie J, Ma J. 2025.Unveiling Sec31: The regulator of COPII vesicle-dependent secretion and beyond. iScience 28(10).

Novick P, Field C, Schekman R. 1980. Identification of 23 complementation groups required for post-translational events in the yeast secretory pathway. Cell 21(1): 205–215.

Obara K, Sekito T, Niimi K, Ohsumi Y. 2008. The Atg18-Atg2 complex is recruited to autophagic membranes via phosphatidylinositol 3-phosphate and exerts an essential function. Journal of Biological Chemistry 283(35): 23972–23980.

Ok HE, Lee SY, Chun HS. 2018. Occurrence and simultaneous determination of nivalenol and deoxynivalenol in rice and bran by HPLC-UV detection and immunoaffinity cleanup. Food Control 87: 53–59.

Ouyang Y, Huang X, Lu Z, Yao J. 2012. Genomic survey, expression profile and co- expression network analysis of *OsWD40* family in rice. BMC Genomics 13: 100.

Salama NR, Chuang JS, Schekman RW. 1997. *Sec31* encodes an essential component of the COPII coat required for transport vesicle budding from the endoplasmic reticulum. Molecular Biology of the Cell 8(2): 205–217.

Sambrook J, Russell DW. 2001. Molecular cloning: a laboratory manual: Cold Spring Harbor Laboratory Press.

Smith TF, Gaitatzes C, Saxena K, Neer EJ. 1999. The WD repeat: a common architecture for diverse functions. Trends in Biochemical Sciences 24(5): 181–185.

Son H, Lee J, Park AR, Lee Y-W. 2011a. ATP citrate lyase is required for normal sexual and asexual development in *Gibberella zeae*. Fungal Genetics and Biology 48(4): 408–417.

Son H, Seo Y-S, Min K, Park AR, Lee J, Jin J-M, Lin Y, Cao P, Hong S-Y, Kim E-K et al. 2011b. A phenome-based functional analysis of transcription factors in the cereal head blight fungus, *Fusarium graminearum*. PLoS Pathogens 7(10): e1002310.

Sondek J, Bohm A, Lambright DG, Hamm HE, Sigler PB. 1996. Crystal structure of a GA protein βγdimer at 2.1 Å resolution. Nature 379(6563): 369–374.

Stajich JE, Harris T, Brunk BP, Brestelli J, Fischer S, Harb OS, Kissinger JC, Li W, Nayak V, Pinney DF et al. 2012. FungiDB: an integrated functional genomics database for fungi. Nucleic Acids Research 40(D1): D675–681.

Szklarczyk D, Kirsch R, Koutrouli M, Nastou K, Mehryary F, Hachilif R, Gable AL, Fang T, Doncheva NT, Pyysalo S et al. 2023. The STRING database in 2023: protein- protein association networks and functional enrichment analyses for any sequenced genome of interest. Nucleic Acids Research 51(D1): D638–646.

Tamura K, Stecher G, Kumar S. 2021. MEGA11: molecular evolutionary genetics analysis version 11. Molecular Biology and Evolution 38(7): 3022–3027.

Tang G, Yuan J, Wang J, Zhang YZ, Xie SS, Wang H, Tao Z, Liu H, Kistler HC, Zhao Y et al. 2021. *Fusarium* BP1 is a reader of H3K27 methylation. Nucleic Acids Research 49(18): 10448–10464.

Thompson JD, Higgins DG, Gibson TJ. 1994. CLUSTAL W: improving the sensitivity of progressive multiple sequence alignment through sequence weighting, position-specific gap penalties and weight matrix choice. Nucleic Acids Research 22(22): 4673–4680.

van Nocker S, Ludwig P. 2003. The WD-repeat protein superfamily in *Arabidopsis*: conservation and divergence in structure and function. BMC Genomics 4(1): 50.

Wall MA, Coleman DE, Lee E, Iñiguez-Lluhi JA, Posner BA, Gilman AG, Sprang SR. 1995. The structure of the G protein heterotrimer Giα1β1γ2. Cell 83(6): 1047–1058.

Wang C, Zhang S, Hou R, Zhao Z, Zheng Q, Xu Q, Zheng D, Wang G, Liu H, Gao X. 2011. Functional analysis of the kinome of the wheat scab fungus *Fusarium graminearum*. PLoS Pathogens 7(12): e1002460.

Wang Y, Hu XJ, Zou XD, Wu XH, Ye ZQ, Wu YD. 2015. WDSPdb: a database for WD40- repeat proteins. Nucleic Acids Research 43(D1): D339–344.

Windels CE. 2000. Economic and social impacts of Fusarium head blight: changing farms and rural communities in the northern great plains. Phytopathology 90(1): 17–21.

Wu XH, Wang Y, Zhuo Z, Jiang F, Wu YD. 2012. Identifying the hotspots on the top faces of WD40-repeat proteins from their primary sequences by beta-bulges and DHSW tetrads. PLoS One 7(8): e43005.

Xie Y, Li H, Luo X, Li H, Gao Q, Zhang L, Teng Y, Zhao Q, Zuo Z, Ren J. 2022. IBS 2.0: an upgraded illustrator for the visualization of biological sequences. Nucleic Acids Research 50(W1): W420–426.

Xu C, Min J. 2011. Structure and function of WD40 domain proteins. Protein & Cell 2(3): 202–214.

Ye J, Zhang Y, Cui H, Liu J, Wu Y, Cheng Y, Xu H, Huang X, Li S, Zhou A et al. 2018. WEGO 2.0: a web tool for analyzing and plotting GO annotations, 2018 update. Nucleic Acids Research 46(W1): W71–75.

Yu J-H, Hamari Z, Han K-H, Seo J-A, Reyes-Domínguez Y, Scazzocchio C. 2004. Double-joint PCR: a PCR-based molecular tool for gene manipulations in filamentous fungi. Fungal Genetics and Biology 41(11): 973–981.

Yun Y, Liu Z, Yin Y, Jiang J, Chen Y, Xu JR, Ma Z. 2015. Functional analysis of the *Fusarium graminearum* phosphatome. New Phytologist 207(1): 119–134.

Zhang C, Zhang F. 2015. The multifunctions of WD40 proteins in genome integrity and cell cycle progression. Journal of Genomics 3: 40–50.

Zheng H, Miao P, Lin X, Li L, Wu C, Chen X, Abubakar YS, Norvienyeku J, Li G, Zhou J et al. 2018. Small GTPase Rab7-mediated FgAtg9 trafficking is essential for autophagy-dependent development and pathogenicity in *Fusarium graminearum*. PLoS Genetics 14(7): e1007546.

Zhou X, Li G, Xu JR 2011. Efficient approaches for generating GFP fusion and epitope- tagging constructs in filamentous fungi. In: Xu J-R, Bluhm BH eds. Fungal Genomics: Methods and Protocols. New York, NY: Humana Press, 199–212.

Zou X-D, Hu X-J, Ma J, Li T, Ye Z-Q, Wu Y-D. 2016. Genome-wide analysis of WD40 protein family in human. Scientific Reports 6(1): 39262.

