## Supplementary Figures for "High-variance phenome database reveals important roles of WD40 proteins in the plant pathogenic fungus *Fusarium graminearum*"

**The following Supporting Information is available for this article:**

**Table S1.** List of oligonucleotides used in this study

**Table S2.** List of 157 WD40 genes in *F. graminearum*

**Table S3.** Gene Ontology distribution of WD40 genes of *F. graminearum*

**Table S4.** Kyoto Encyclopedia of Genes and Genomes pathway enrichment of WD40 proteins in *F. graminearum*

**Table S5.** Phenotypes of 119 WD40 deletion mutants in *F. graminearum*.

**Table S6.** The result of affinity purification-mass spectrometry for Fgwd101.

**Table S7.** The result of affinity purification-mass spectrometry for Fgwd133.

**Fig. S1.** Gene Ontology classification and Kyoto Encyclopedia of Genes and Genomes pathway enrichment analysis of WD40 proteins in *F. graminearum*.

**Fig. S2.** Southern blot validation of WD40 gene deletion mutants and the GFP-Atg8 strain in *F. graminearum*.

**Fig. S3.** Perithecial cirrhus formation on carrot agar medium at 9 days post-inoculation.

**Fig. S4.** Assessment of male and female fertility in WD40 gene deletion mutants of *F. graminearum*.

**Fig. S5.** Virulence of *F. graminearum* WD40 deletion mutants on wheat heads.

**Fig. S6.** Phenotypic correlation analysis of *F. graminearum* WD40 gene deletion mutants.

**Fig. S7.** Immunoprecipitation of GFP-tagged Fgwd101 and Fgwd133 for affinity purification-mass spectrometry analysis and yeast two-hybrid confirmation of the interaction between Fgwd133 and septin complex proteins.

**Note: Tables S1–S7 are provided in a separate Excel file.**

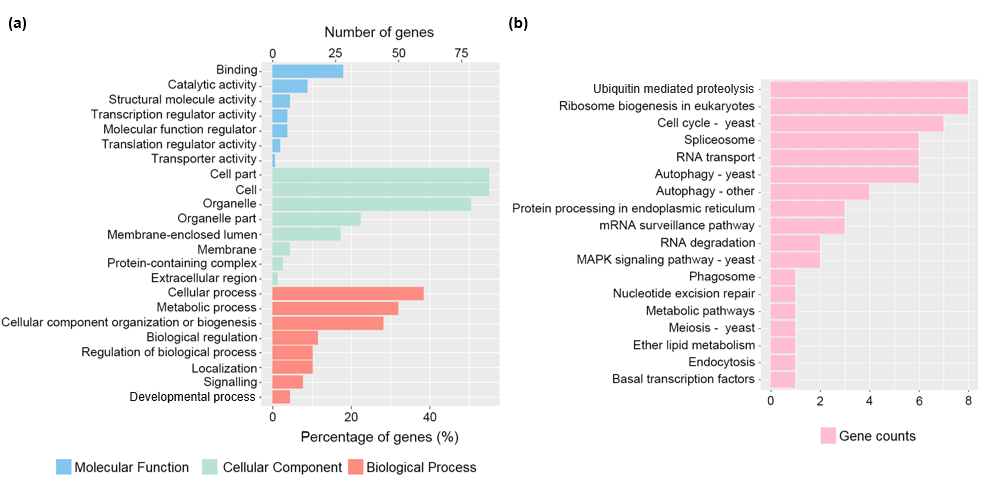

**Fig. S1. Gene Ontology classification and Kyoto Encyclopedia of Genes and Genomes pathway enrichment analysis of WD40 proteins in *F. graminearum*. (a)** Gene Ontology (GO) classification of WD40 proteins based on functional annotation using OmicsBox and visualized with WEGO and R. Detailed GO annotation results are provided in Table S3. **(b)** Kyoto Encyclopedia of Genes and Genomes pathway enrichment analysis of WD40 proteins. Bar graphs indicate the gene count for each enriched pathway. Detailed results are provided in Table S4.

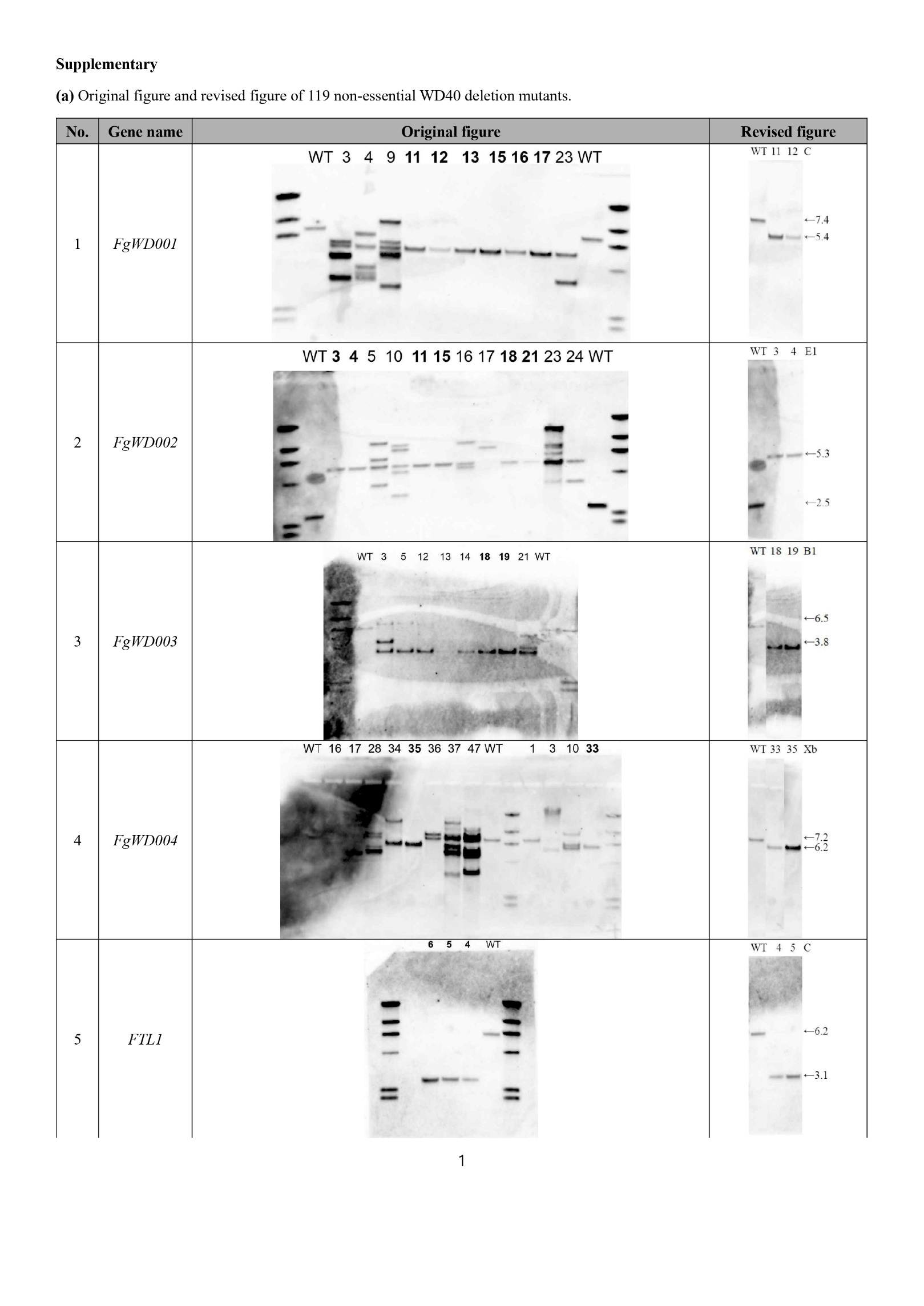

Continued on the following page.

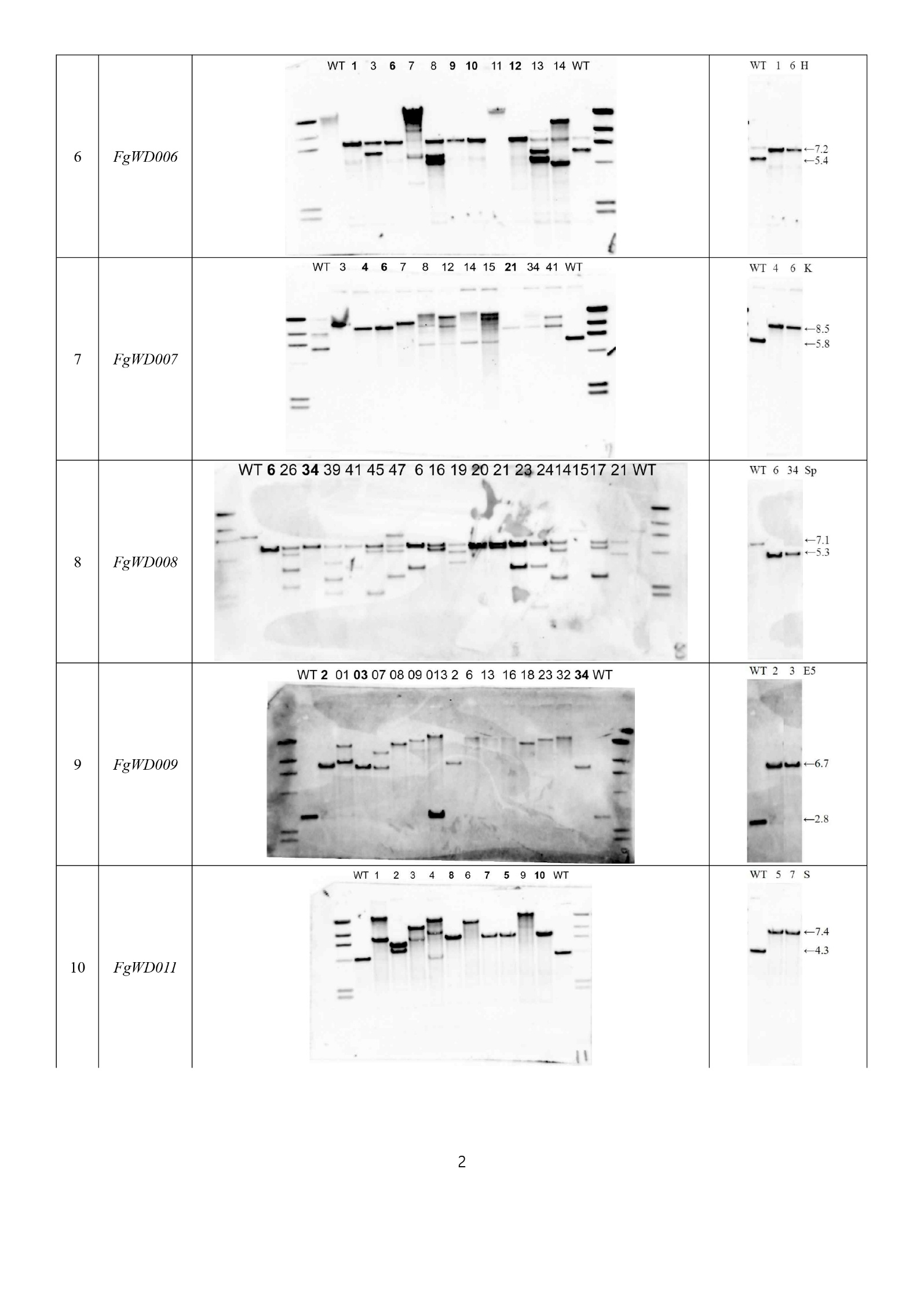

Continued on the following page.
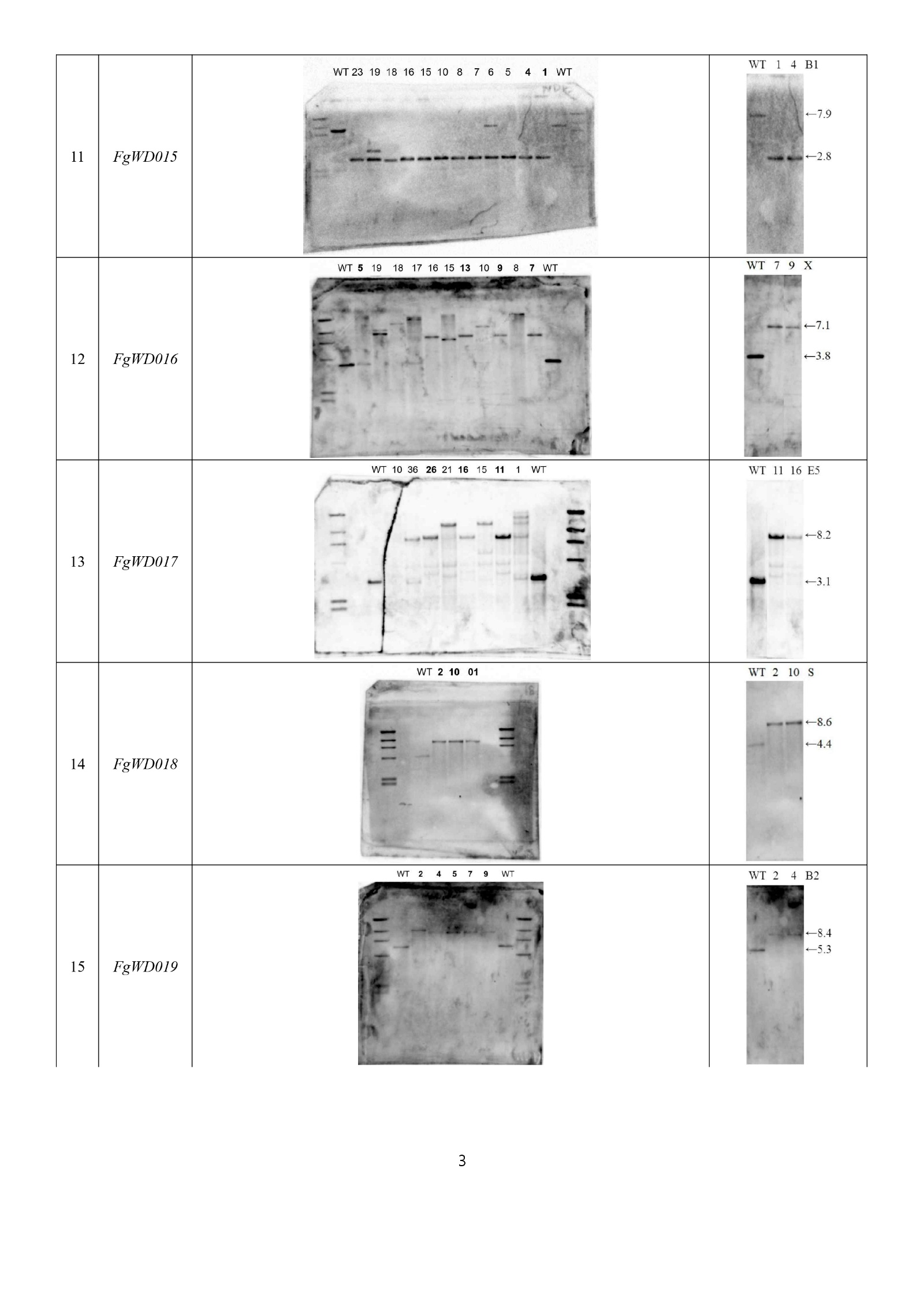

Continued on the following page.

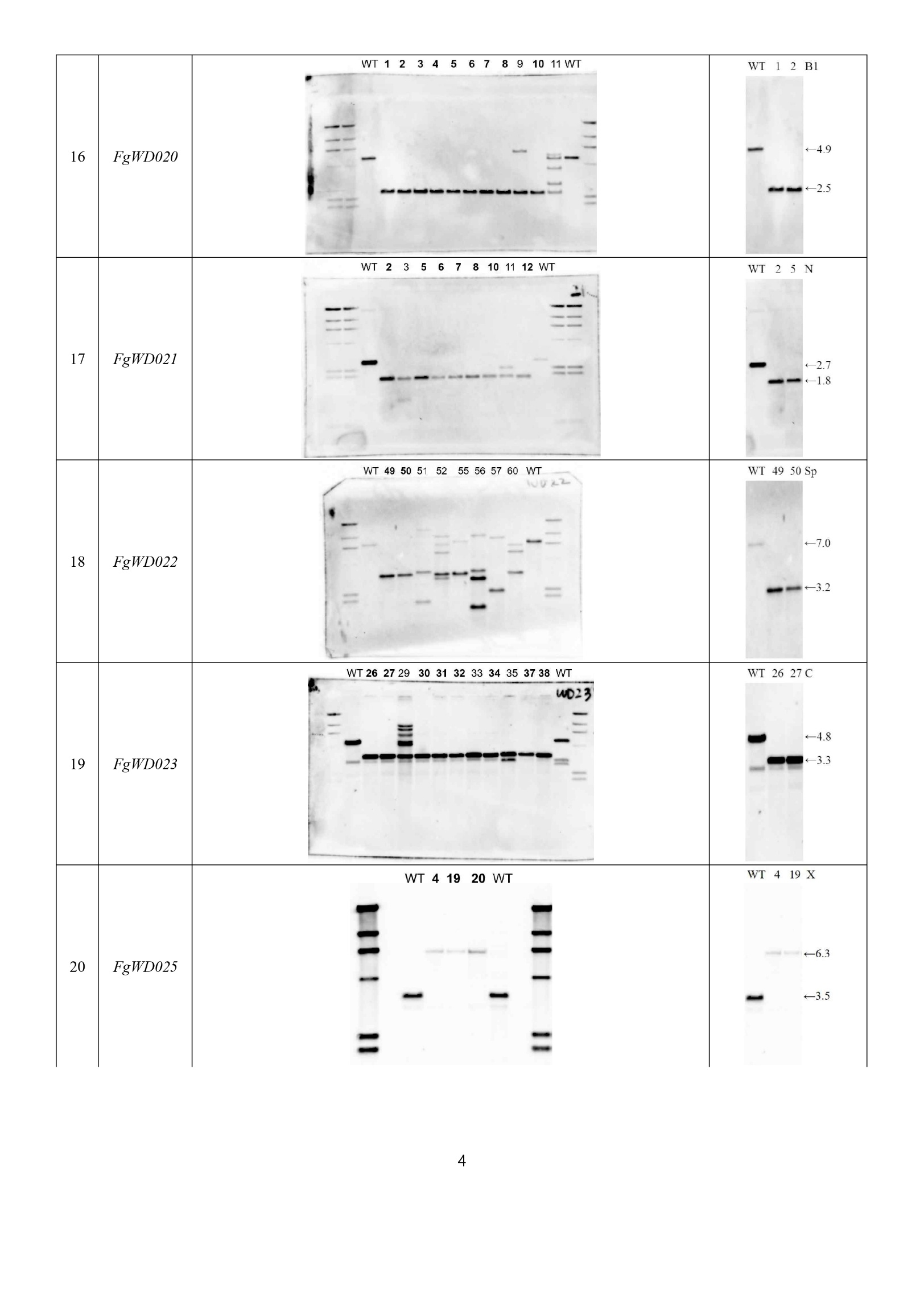

Continued on the following page.

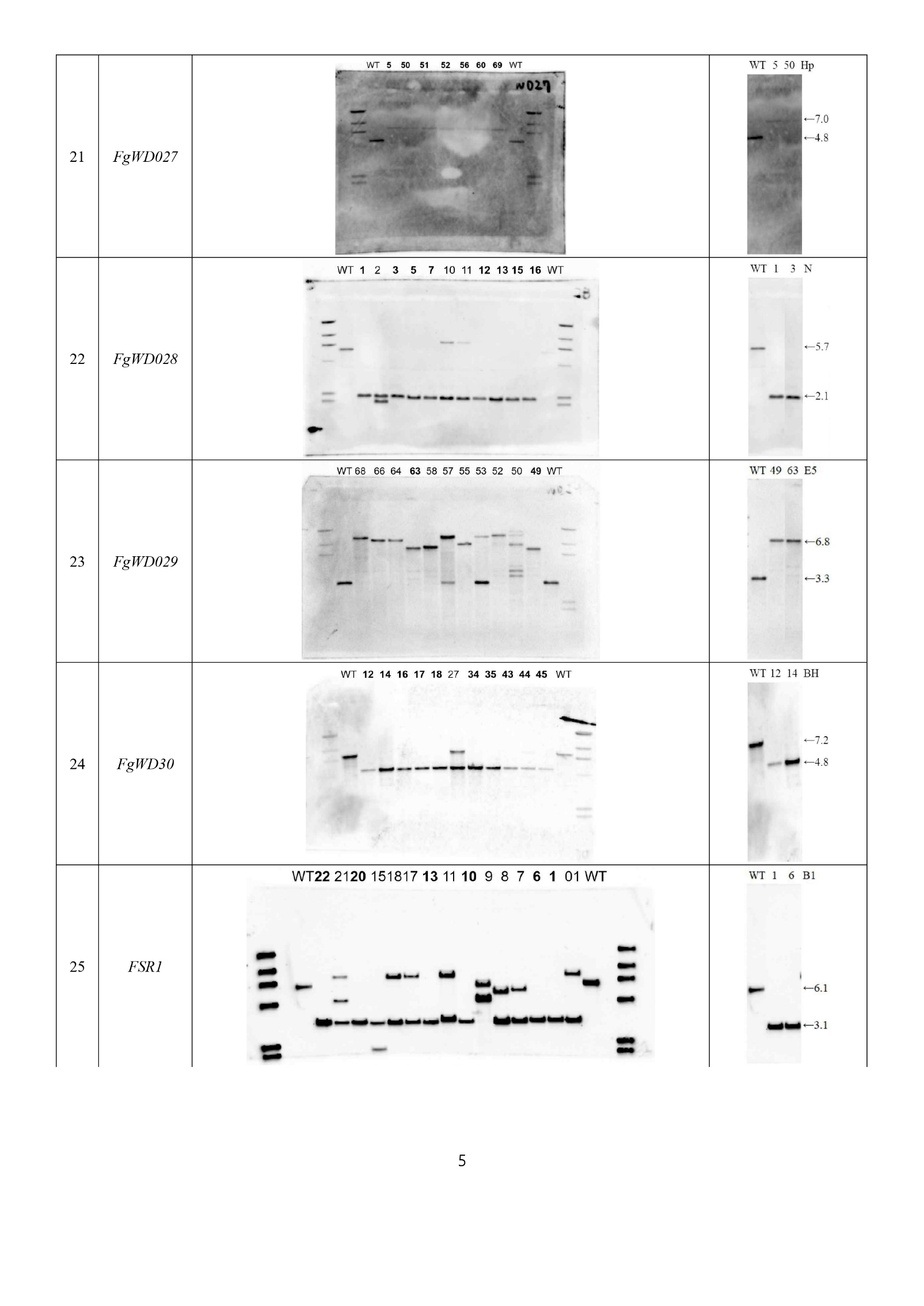

Continued on the following page.

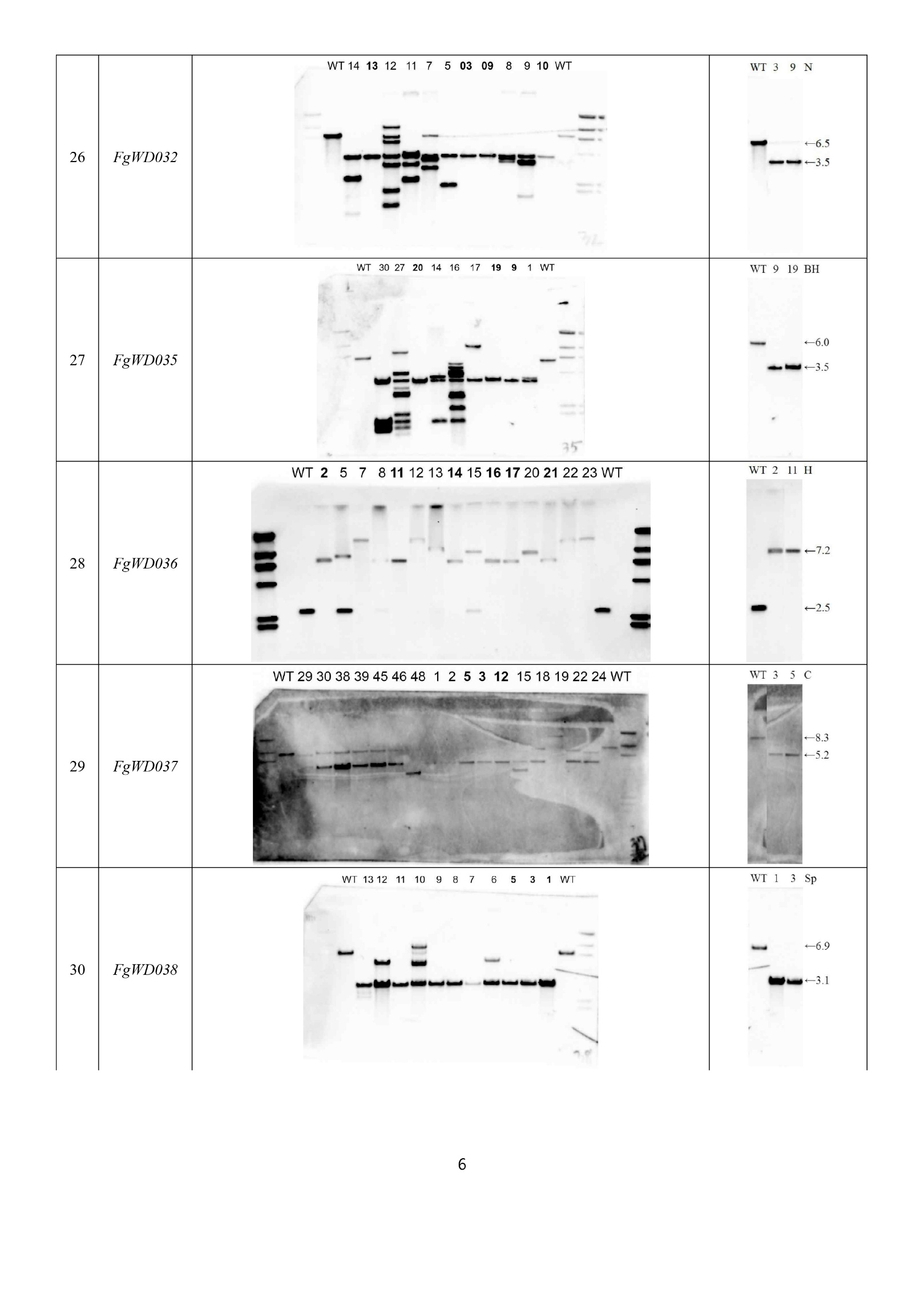

Continued on the following page.

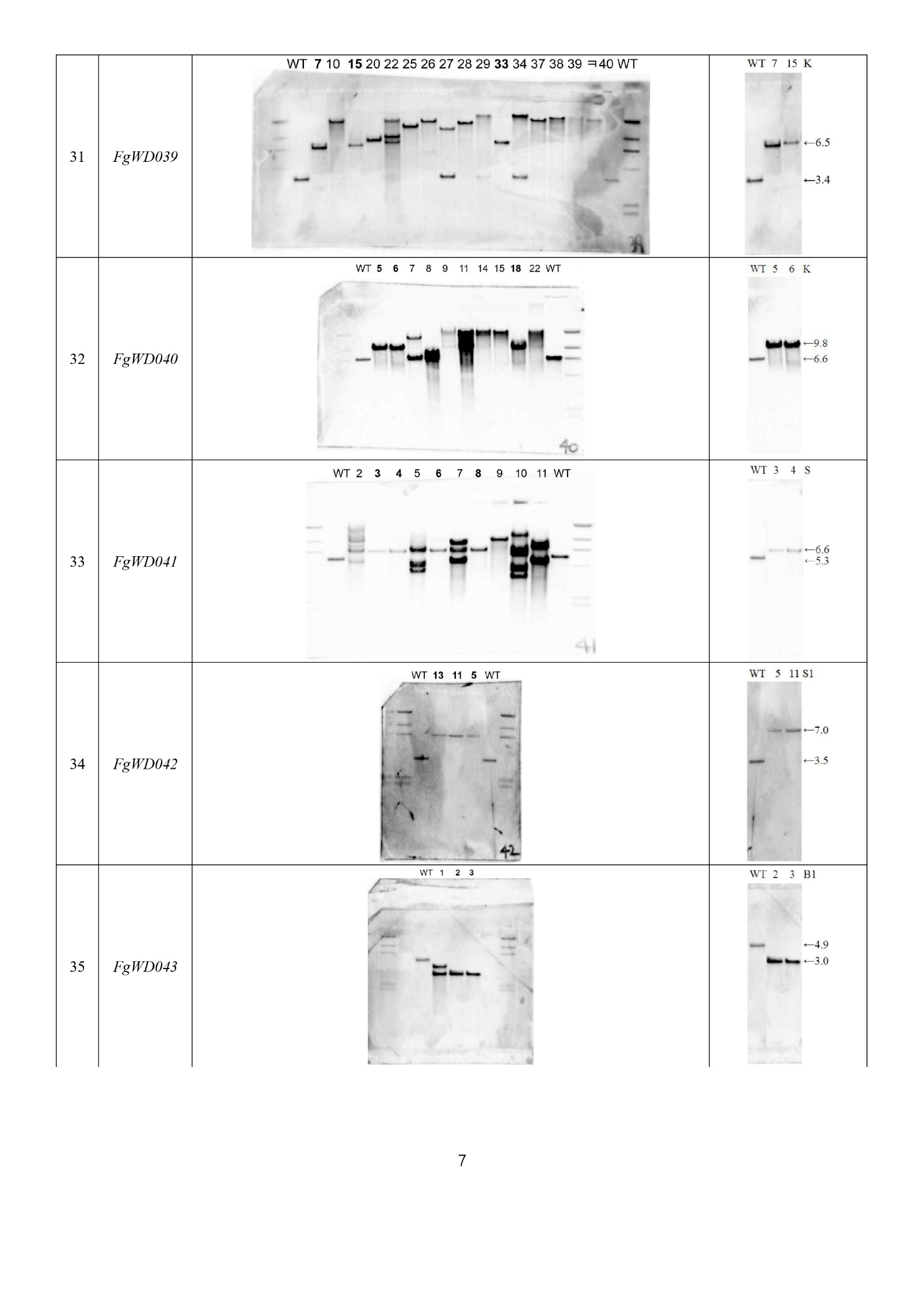

Continued on the following page.

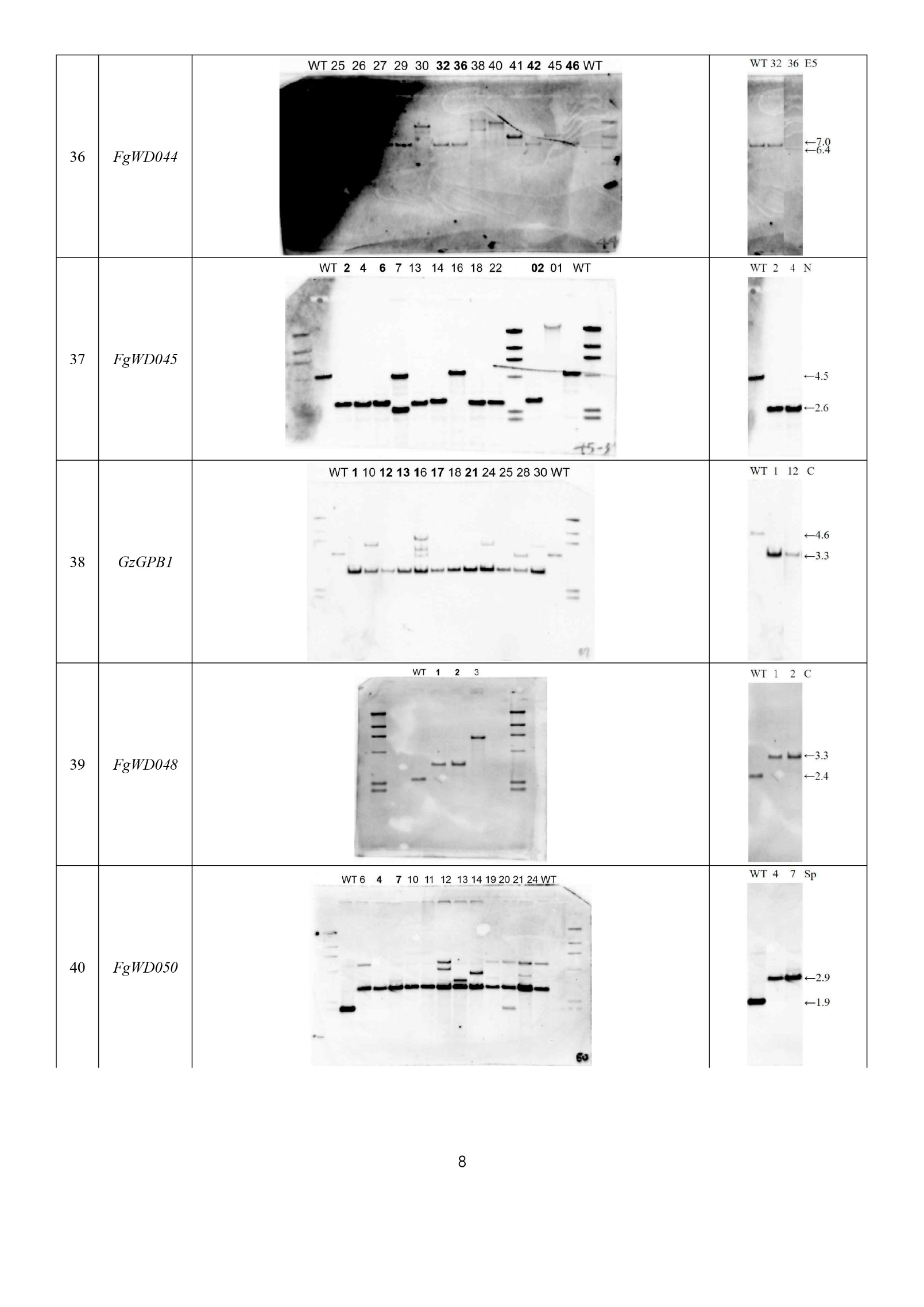

Continued on the following page.

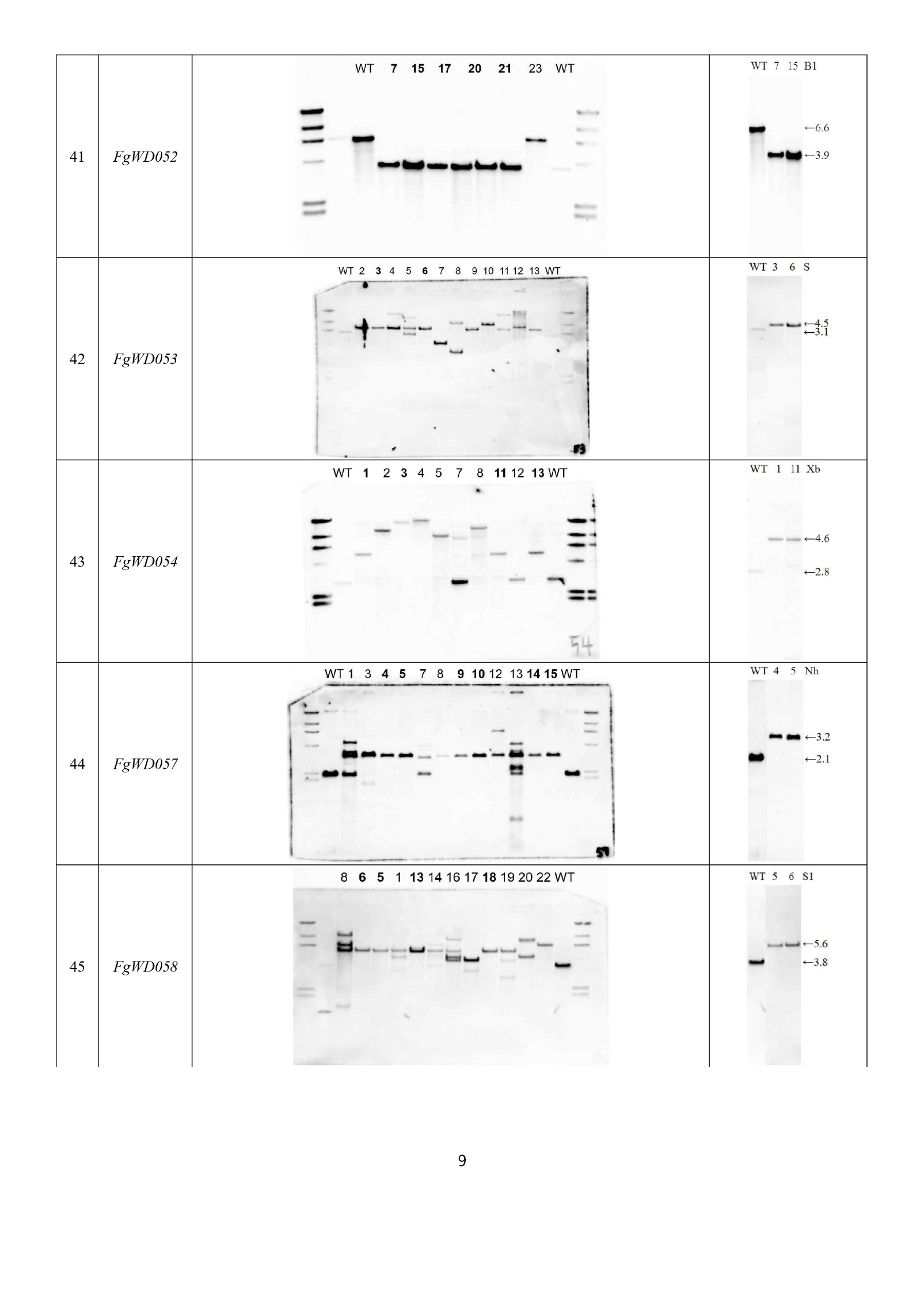

Continued on the following page.

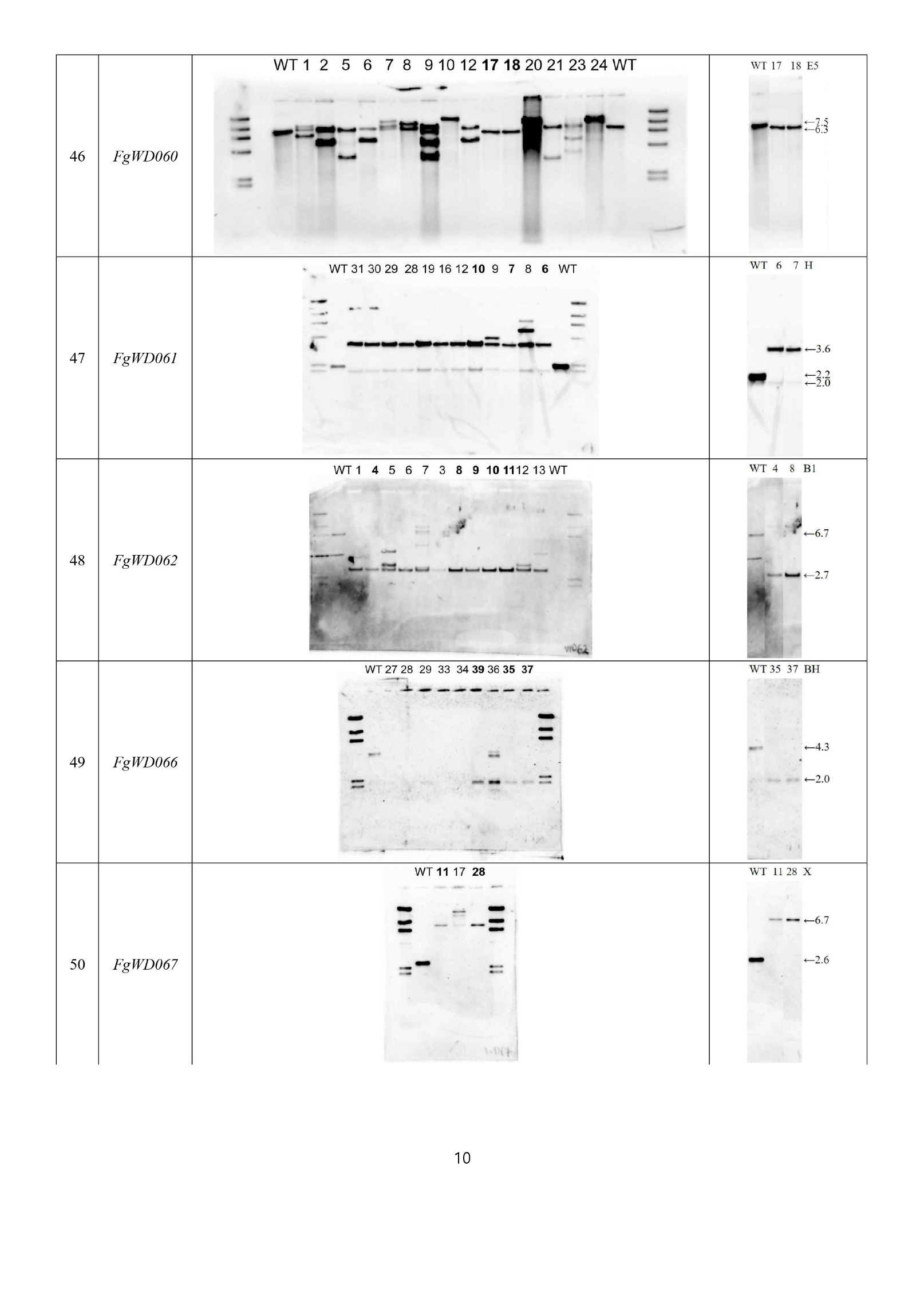

Continued on the following page.

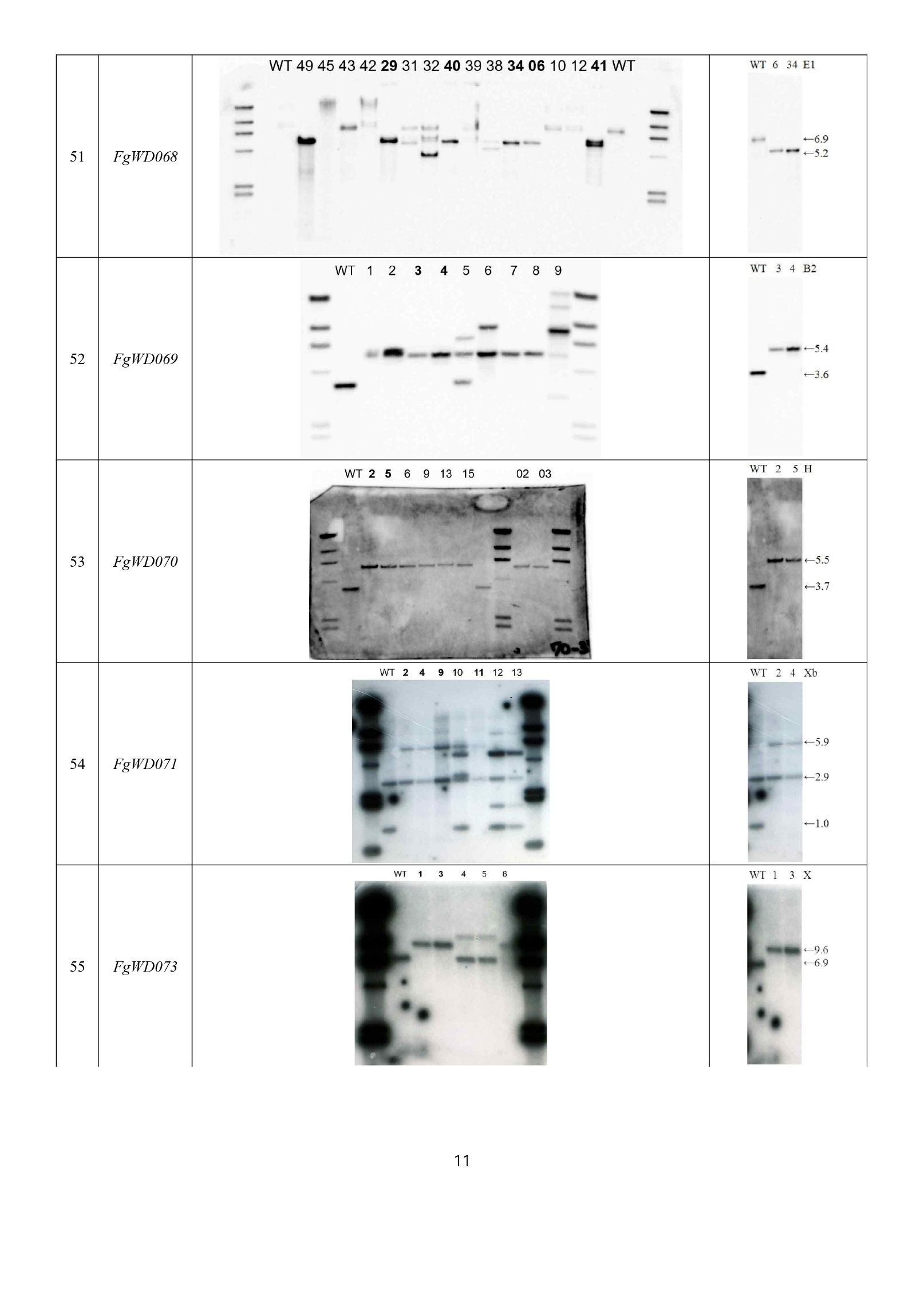

Continued on the following page.

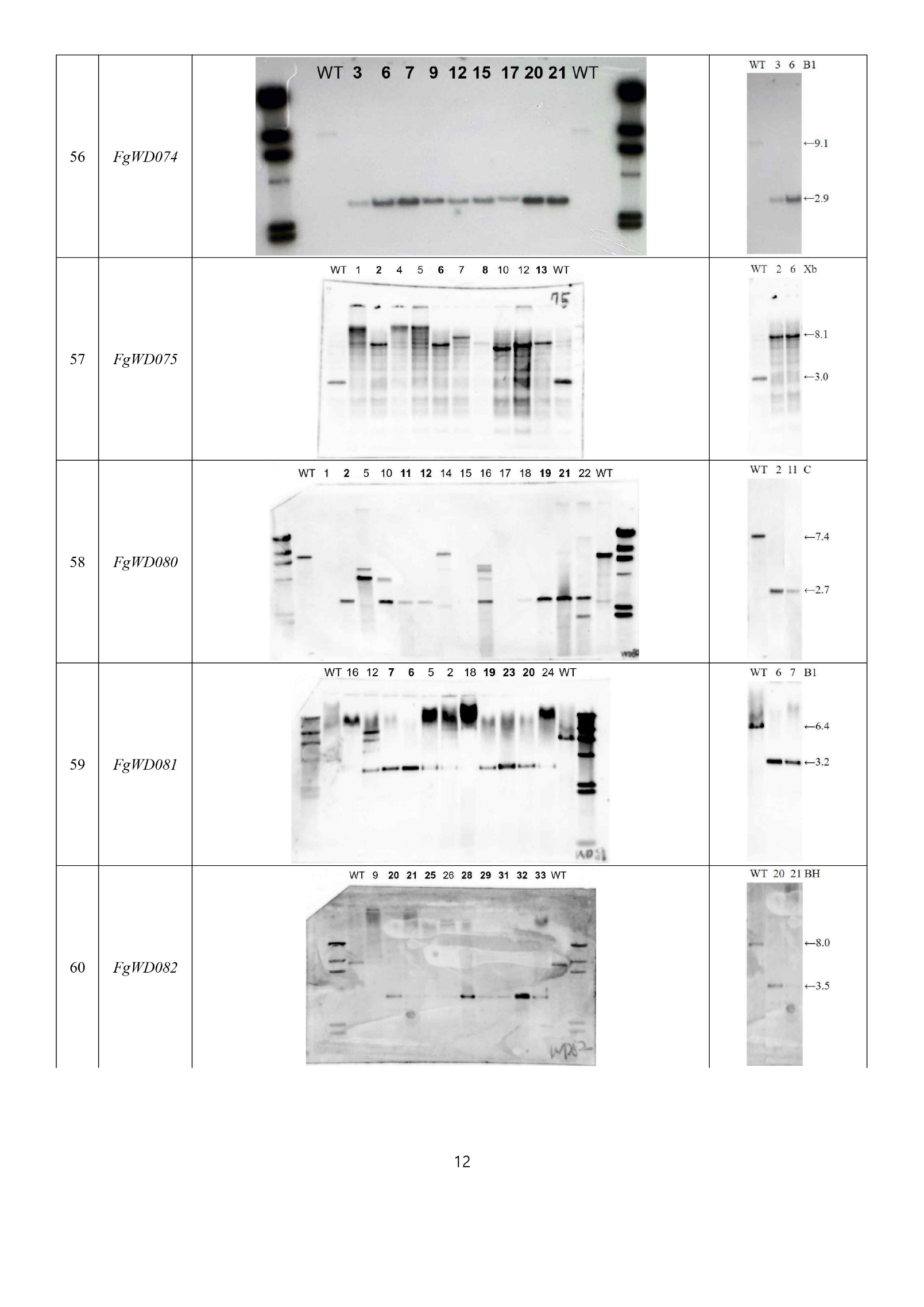

Continued on the following page.

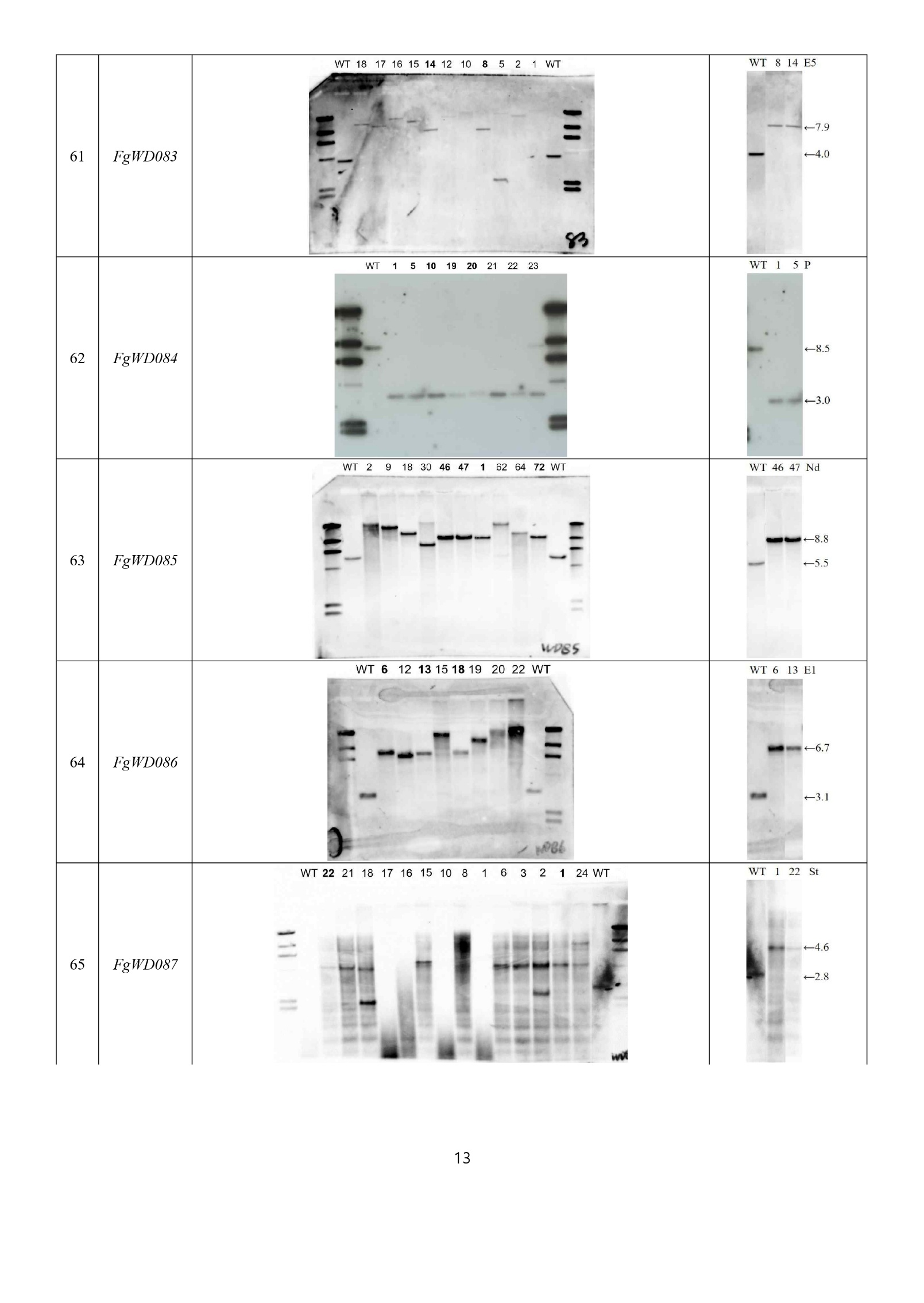

Continued on the following page.

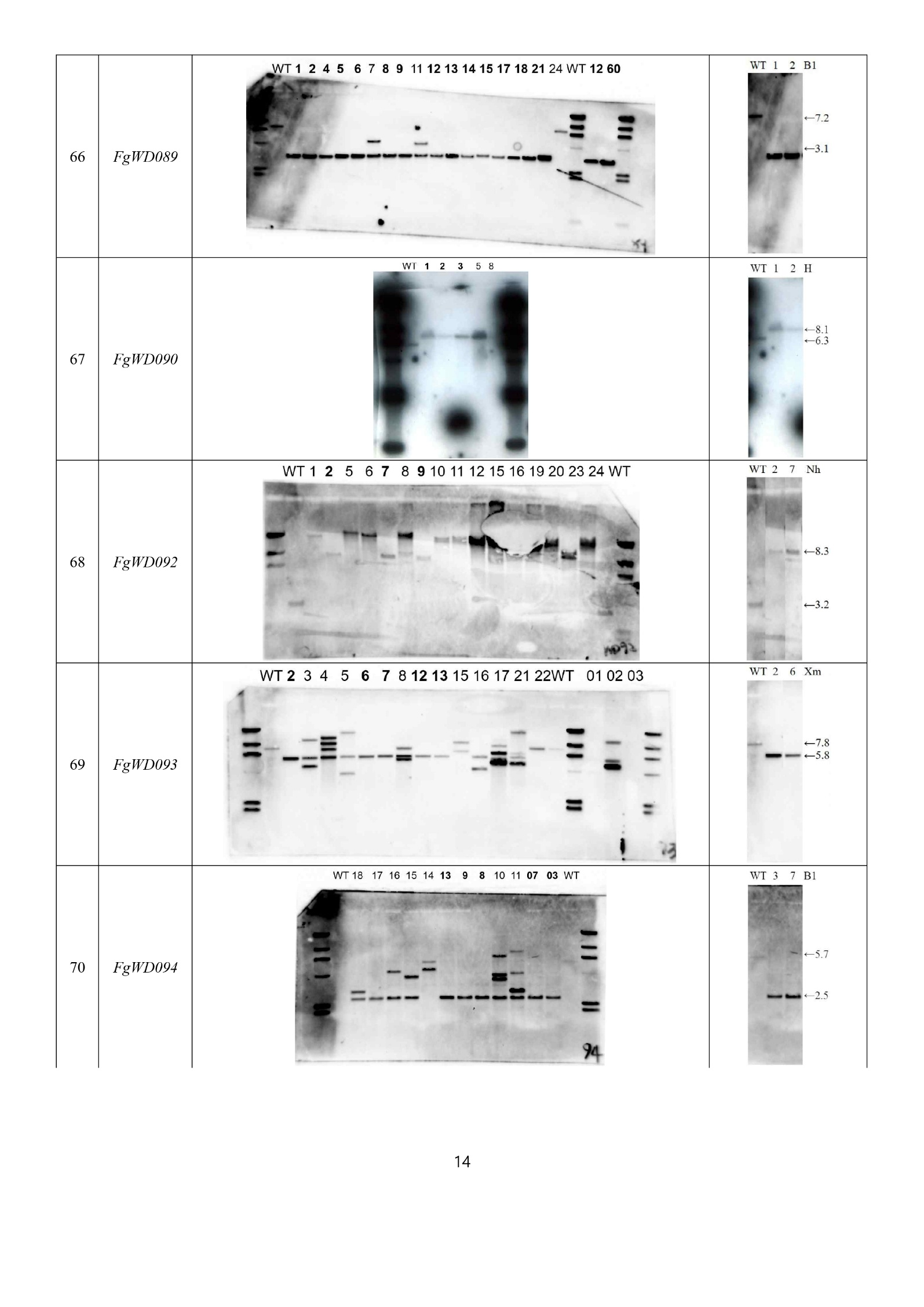

Continued on the following page.

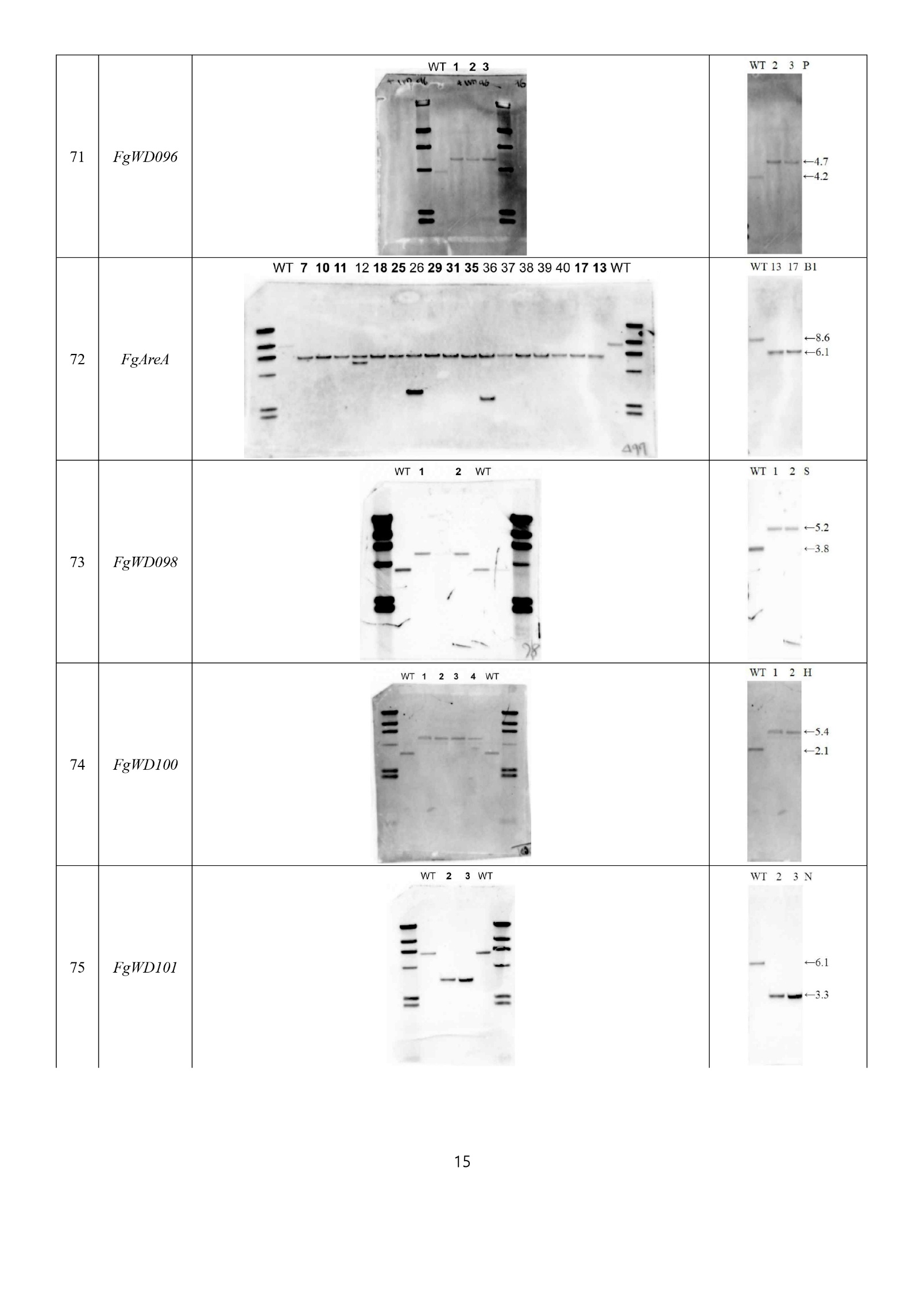

Continued on the following page.

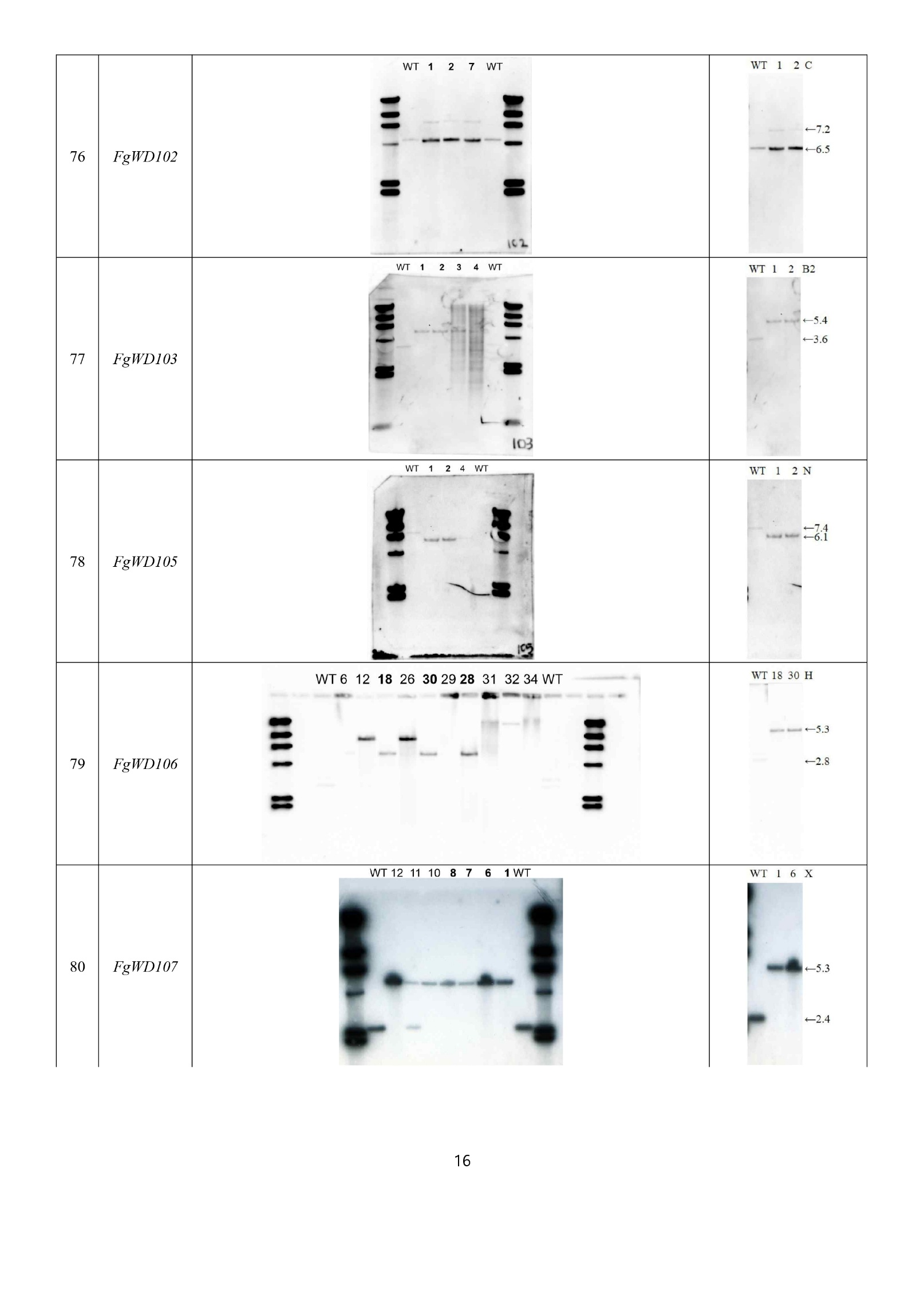

Continued on the following page.

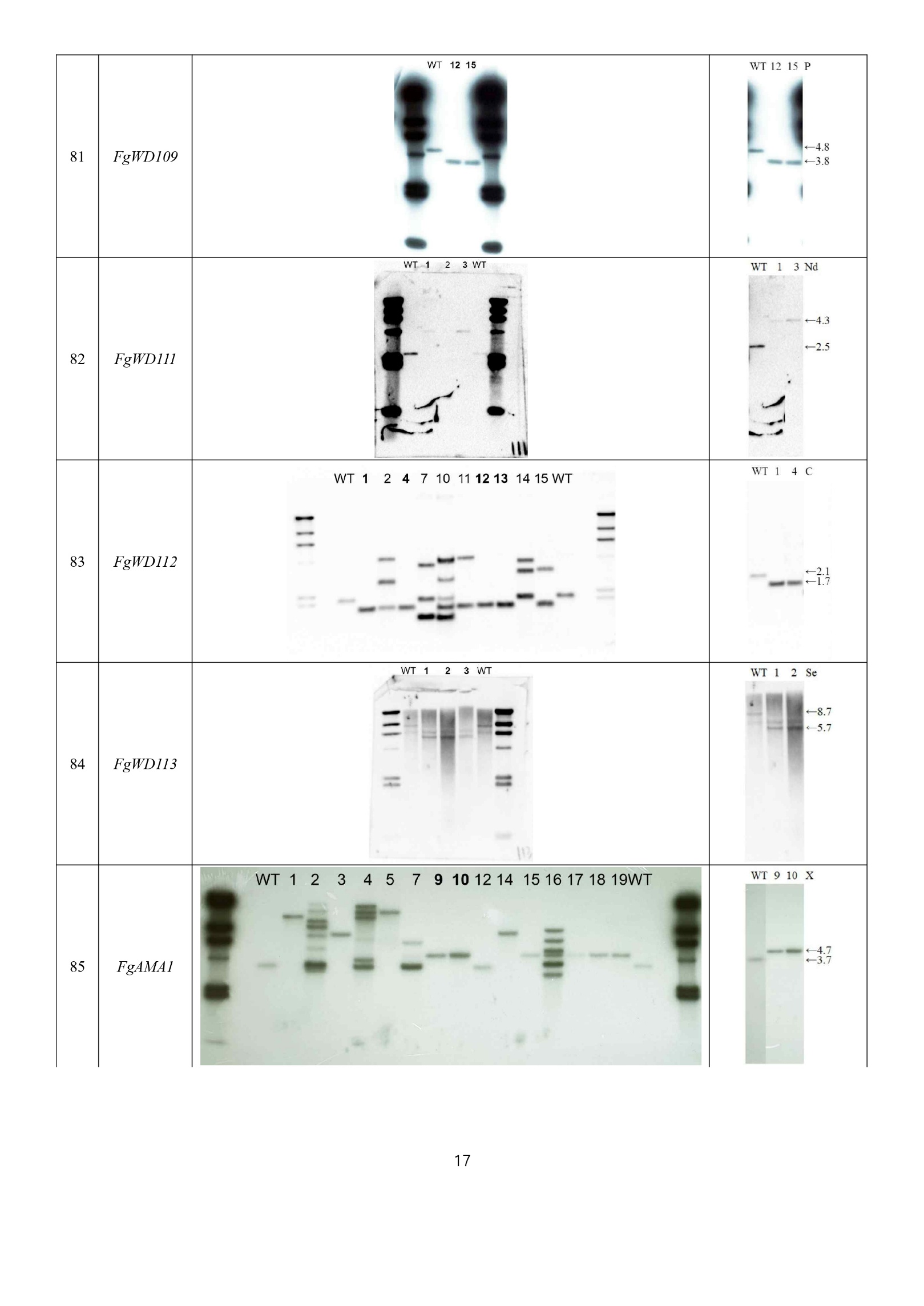

Continued on the following page.

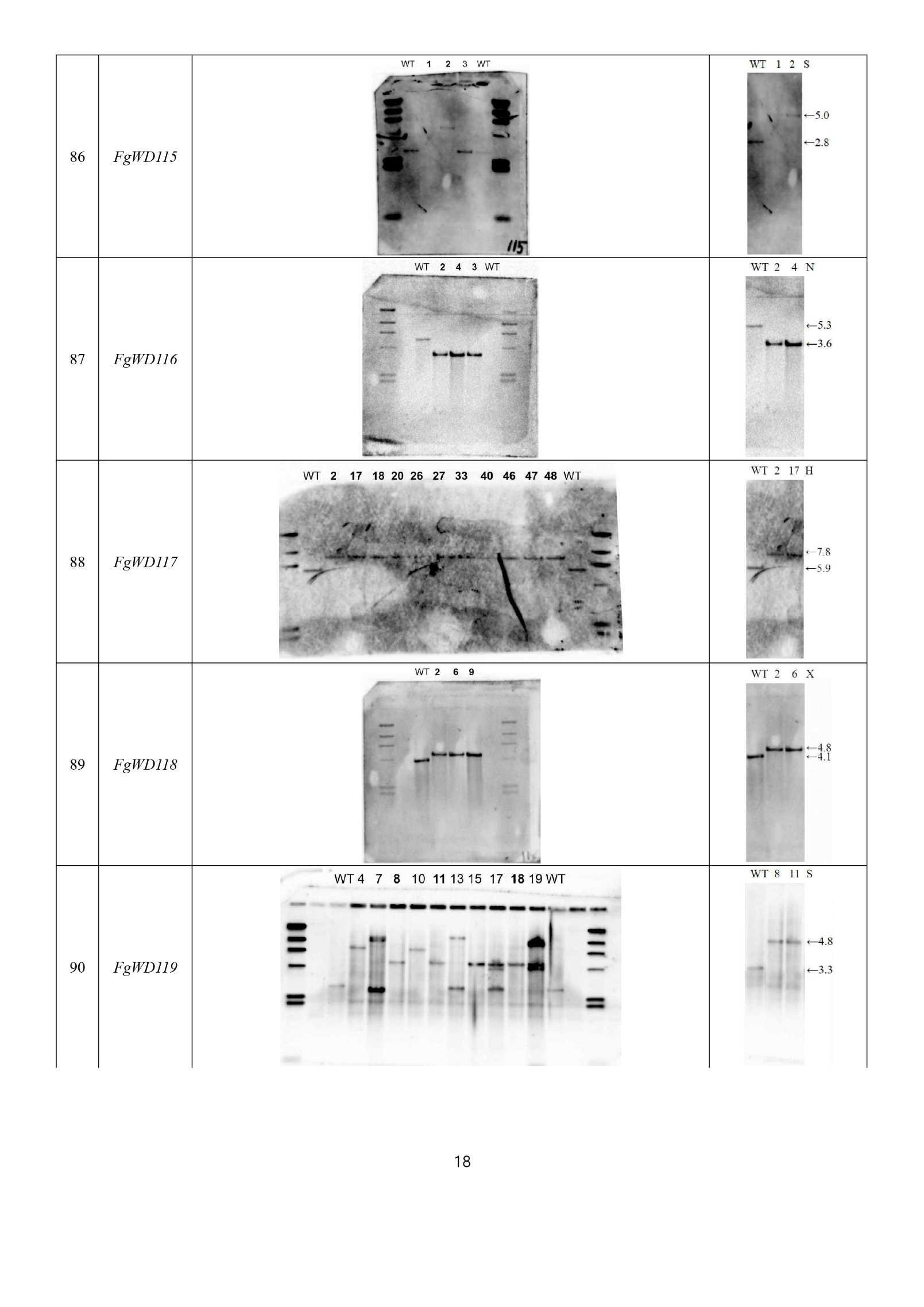

Continued on the following page.

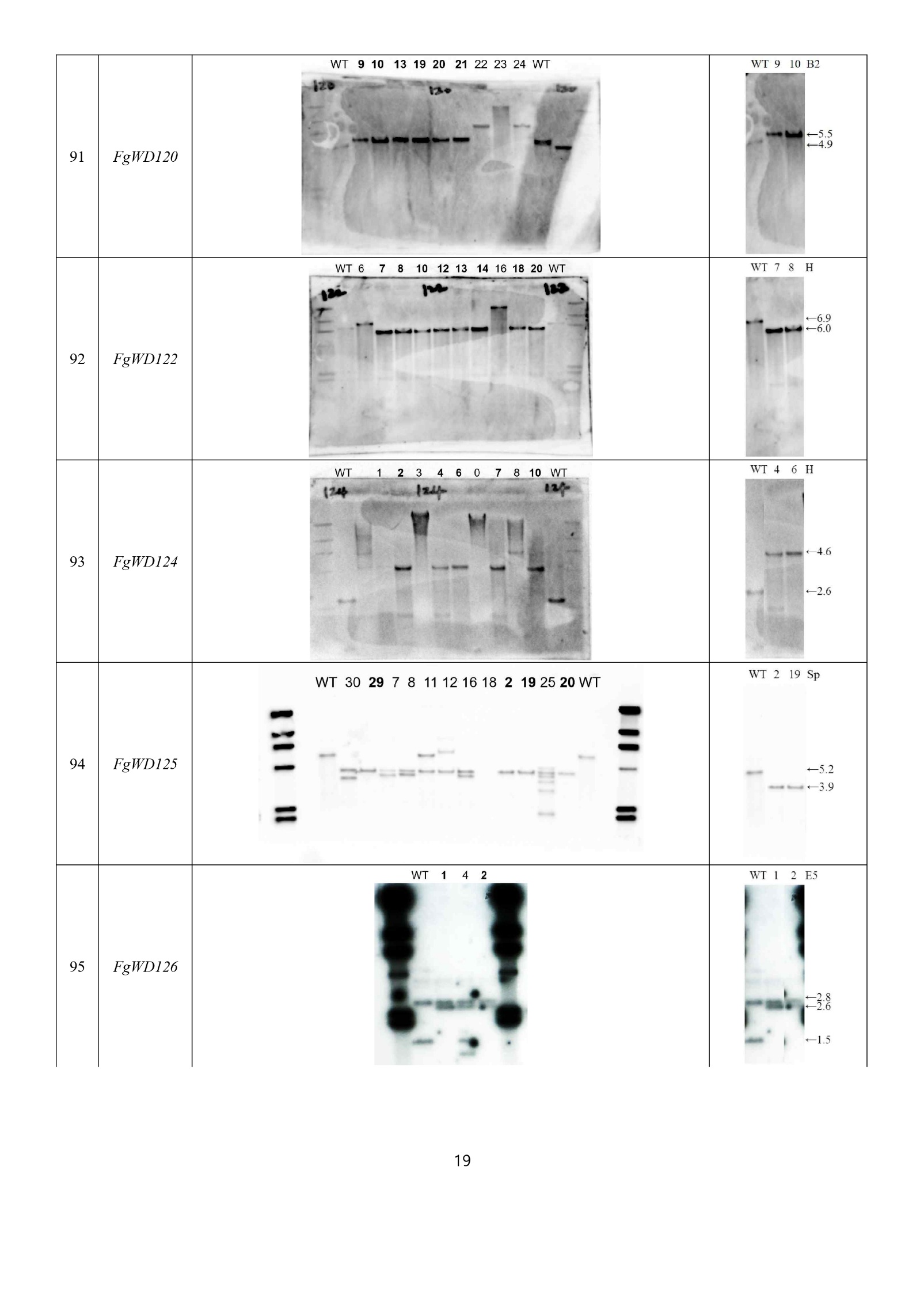

Continued on the following page.

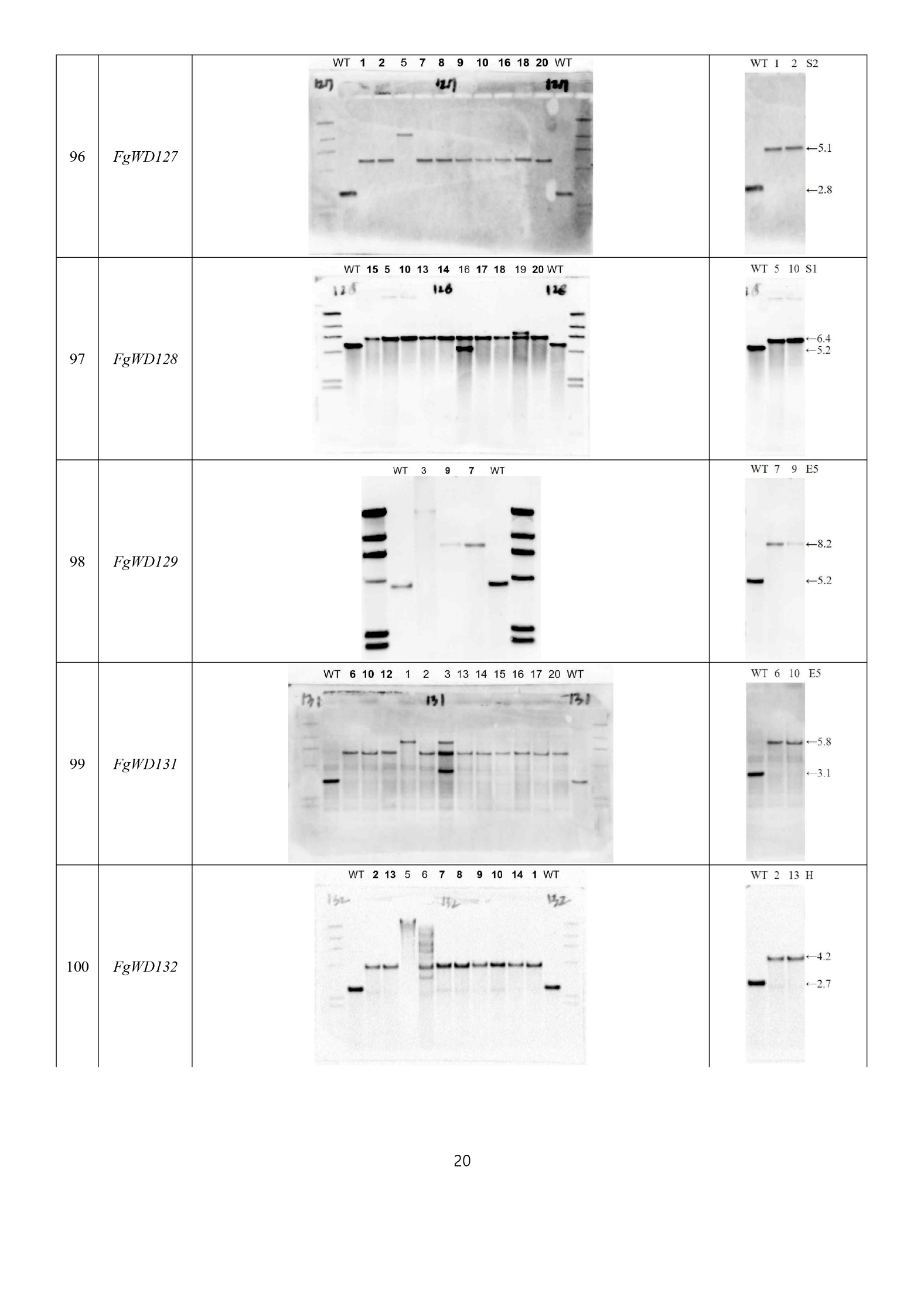

Continued on the following page.

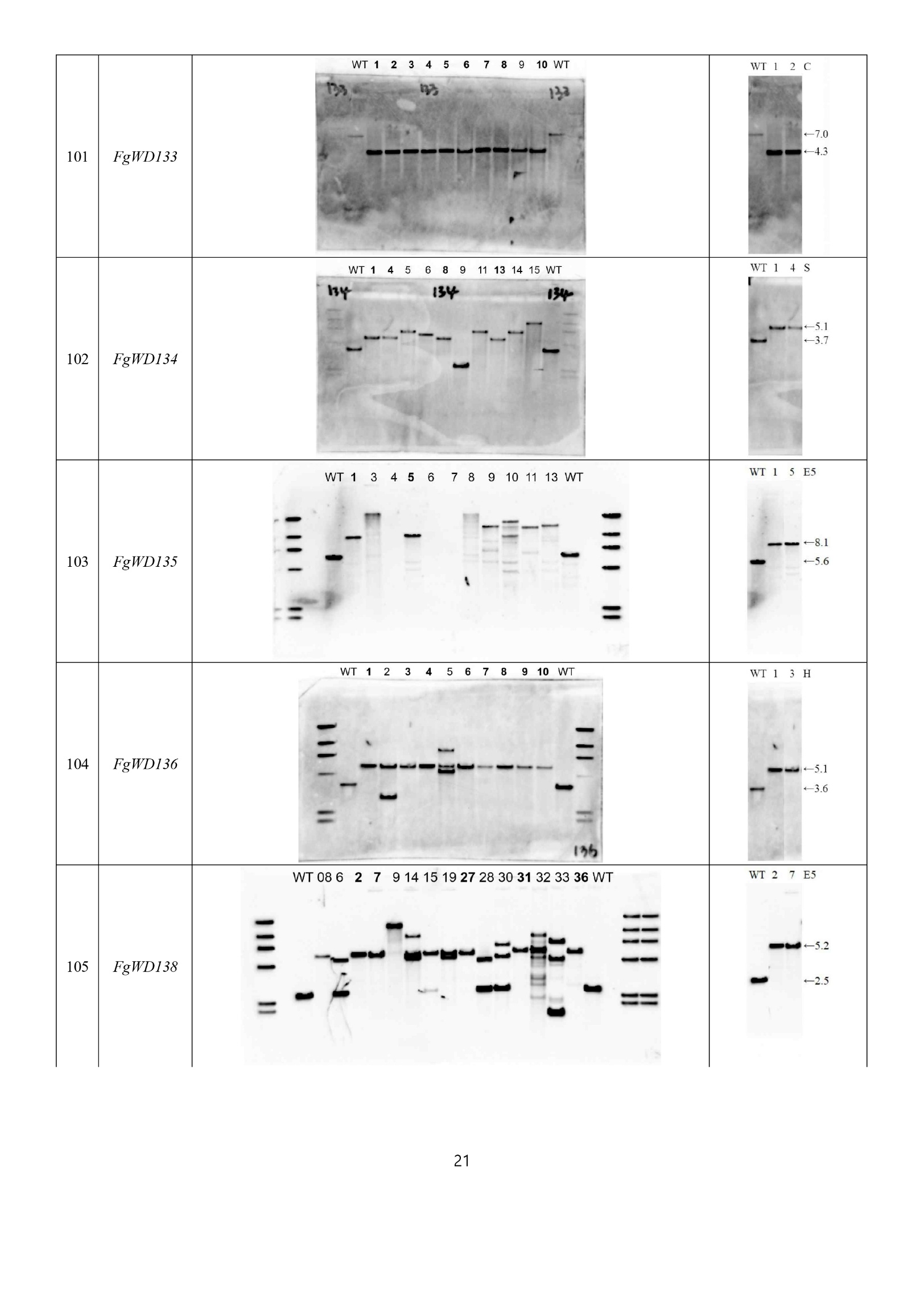

Continued on the following page.

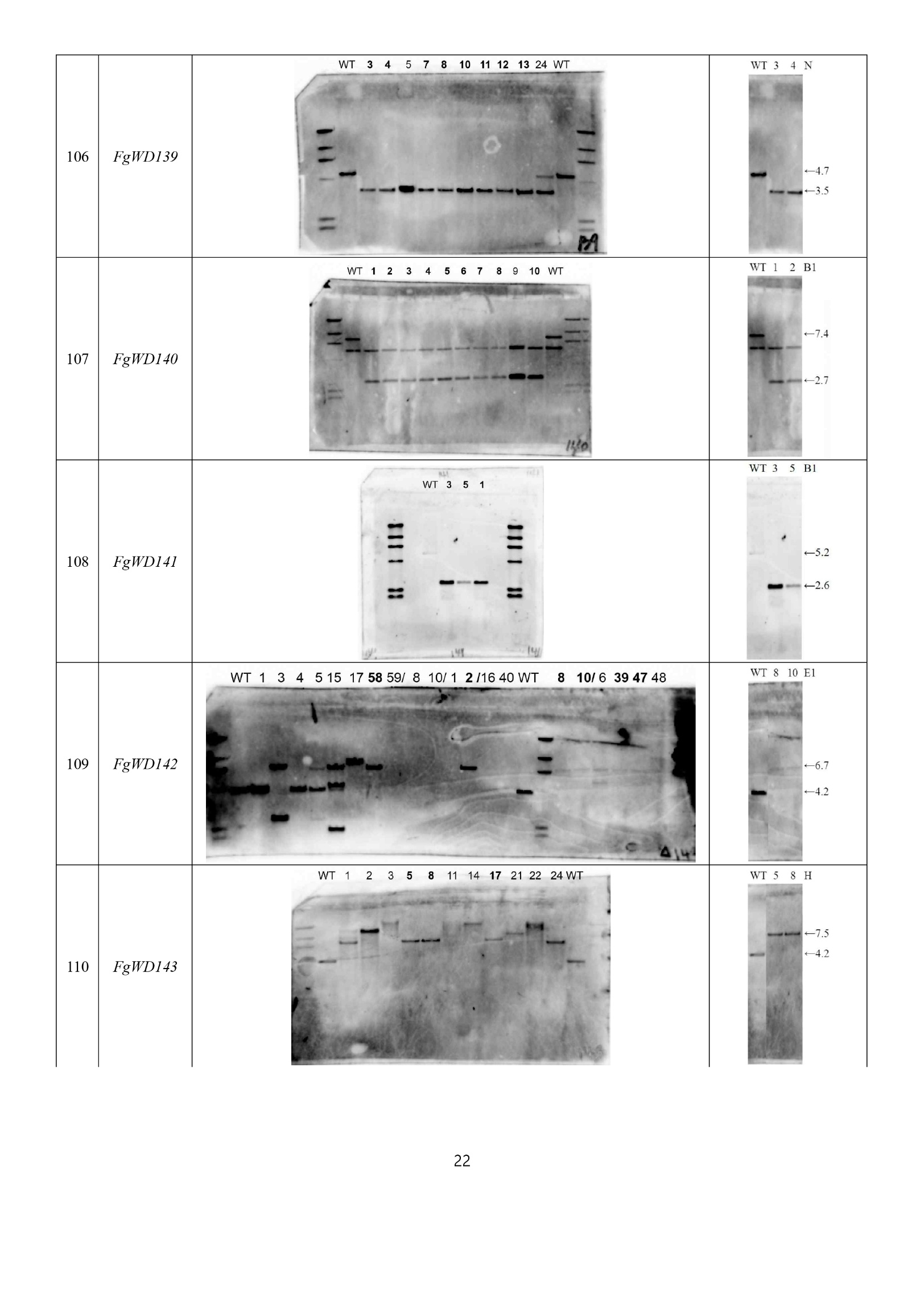

Continued on the following page.

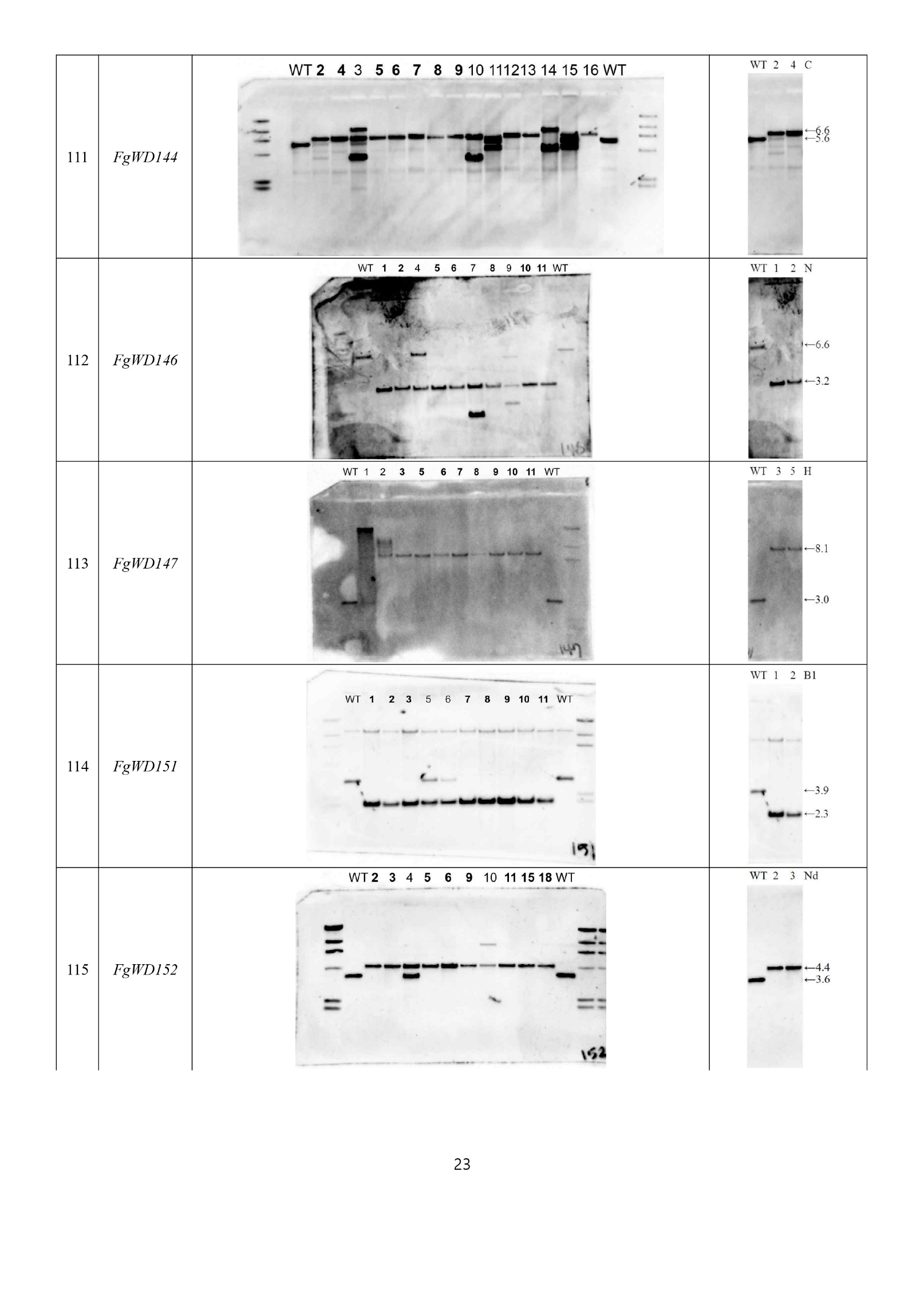

Continued on the following page.

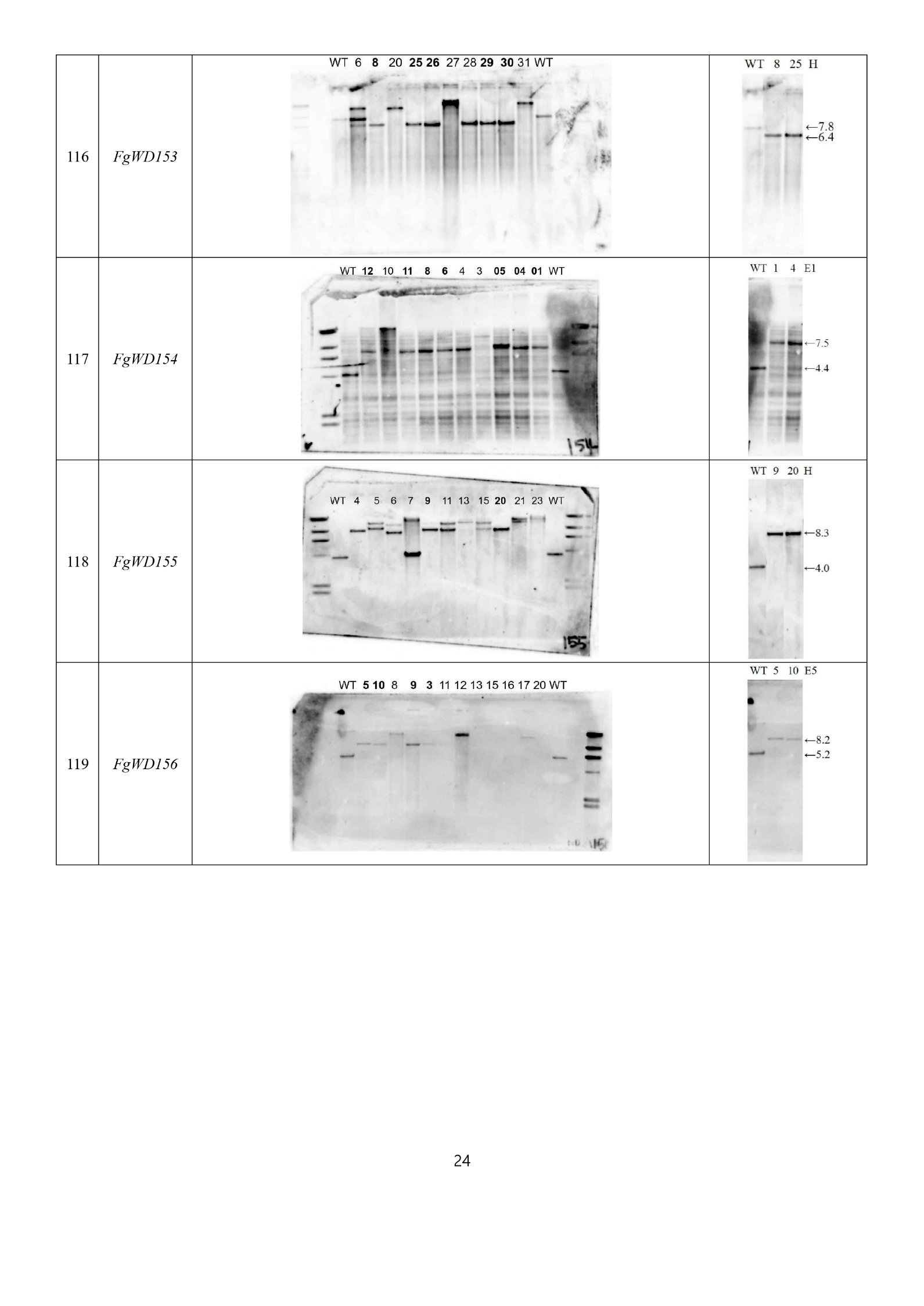

| **Gene name** | **Original figure** | **Revised figure** |
| --- | --- | --- |
| *GFP-Atg8* | 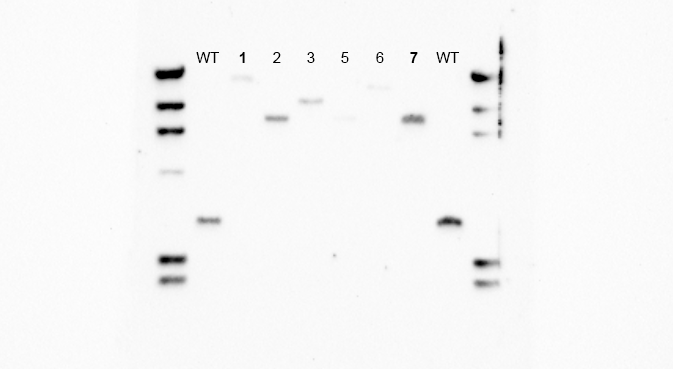 | 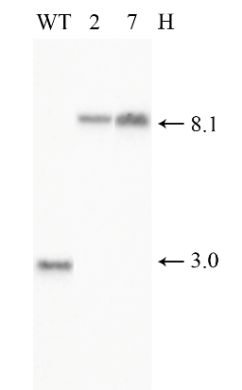 |

**Fig. S2. Southern blot validation of WD40 gene deletion mutants and the GFP-Atg8 strain in *F. graminearum*.** Southern blot analysis confirming the successful generation of 119 non-essential WD40 deletion mutants using gene-specific flanking probes. For each mutant, the uncropped original Southern blots are shown on the left, and the cropped blots containing only the lanes corresponding to the strains used in this study are shown on the right. The restriction enzymes used for genomic DNA digestion and the molecular size markers (kb) are indicated on the right of each blot. A, ApaI; B1, BglI; B2, BglII; BH, BamHI; C, ClaI; E1, EcoRI; E5, EcoRV; H, HindⅢ; Hp, HpaI; K, KpnI; N, NcoI; Nd, NdeI; Nh, NheI; P, PstI; S, SalI; S1, SacI; S2, SacII; Se, SpeI; Sp, SphI; St, StuI; X, XhoI; Xb, XbaI; Xc, XcmI; Xm, XmaI; WT, *F. graminearum* wild-type strain Z-3639; kb, kilobases.

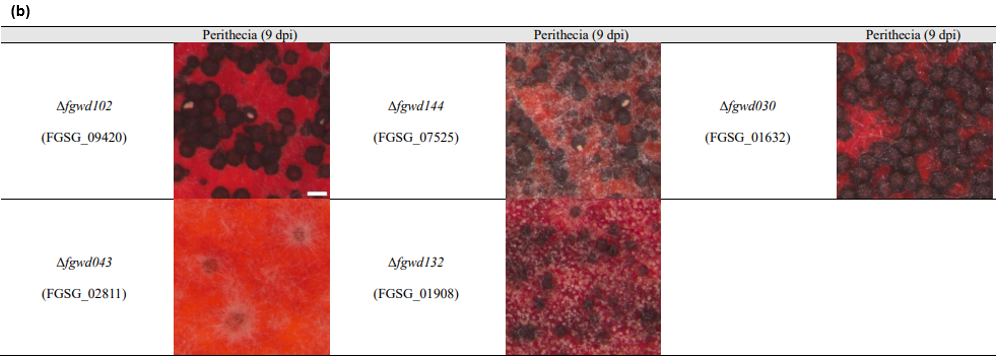

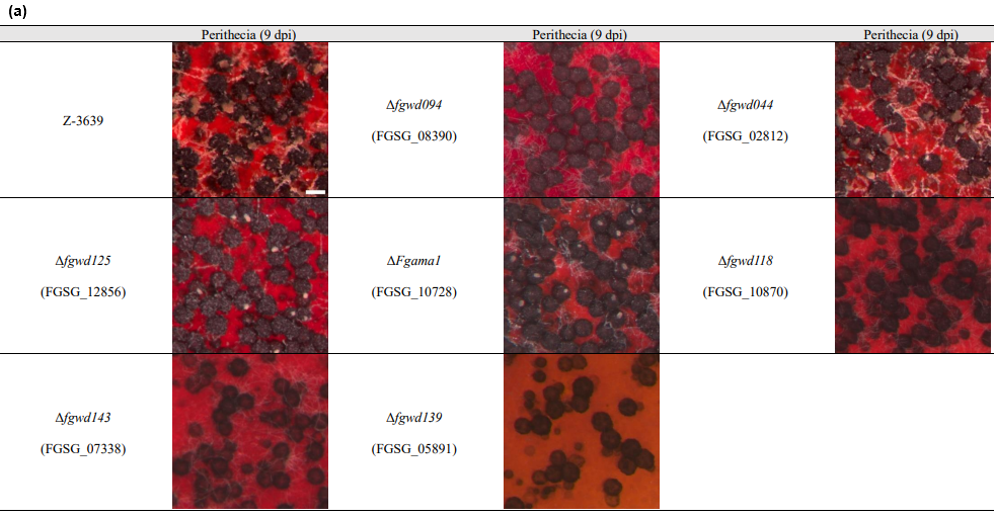

**Fig. S3. Perithecial cirrhus formation on carrot agar medium at 9 days post-inoculation. (a)** WD40 gene deletion mutants belonging to the second sexual development (SD) group. The wild-type strain Z-3639 is shown for comparison. **(b)** WD40 gene deletion mutants belonging to the third SD group. Scale bar = 200 µm. dpi, days post-inoculation.

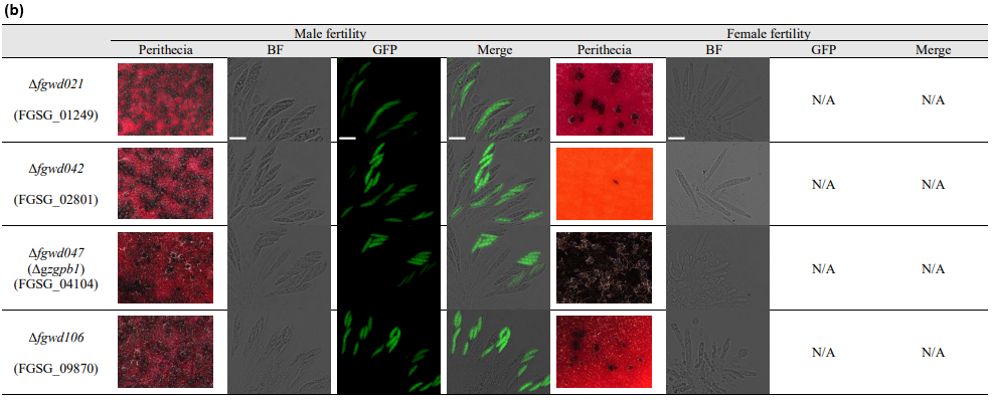

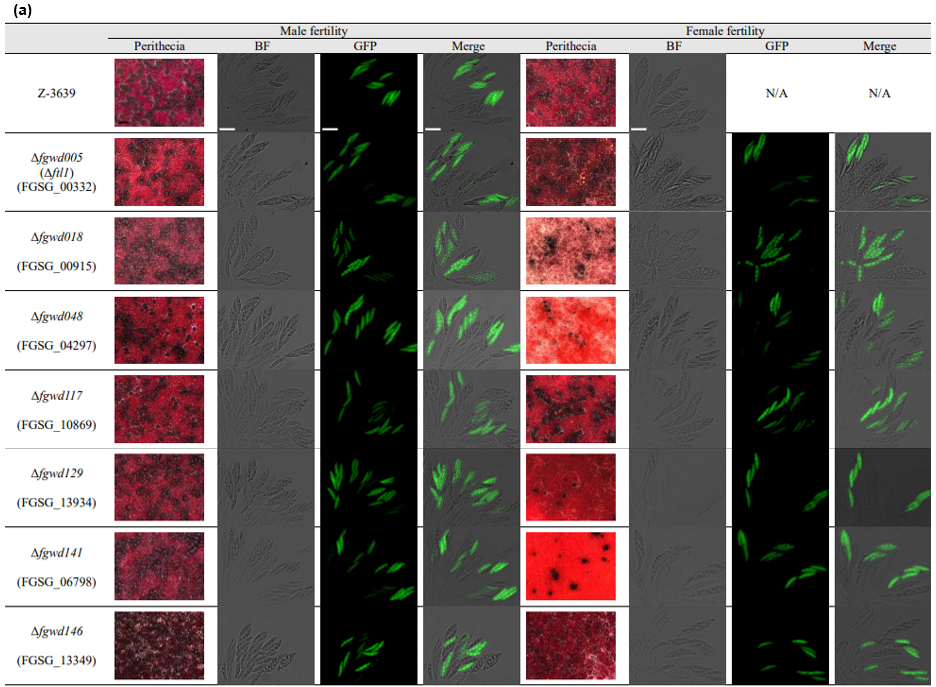
Continued on the following page.

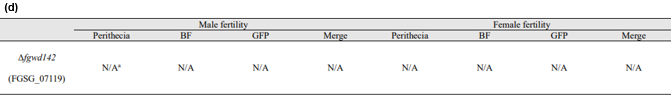

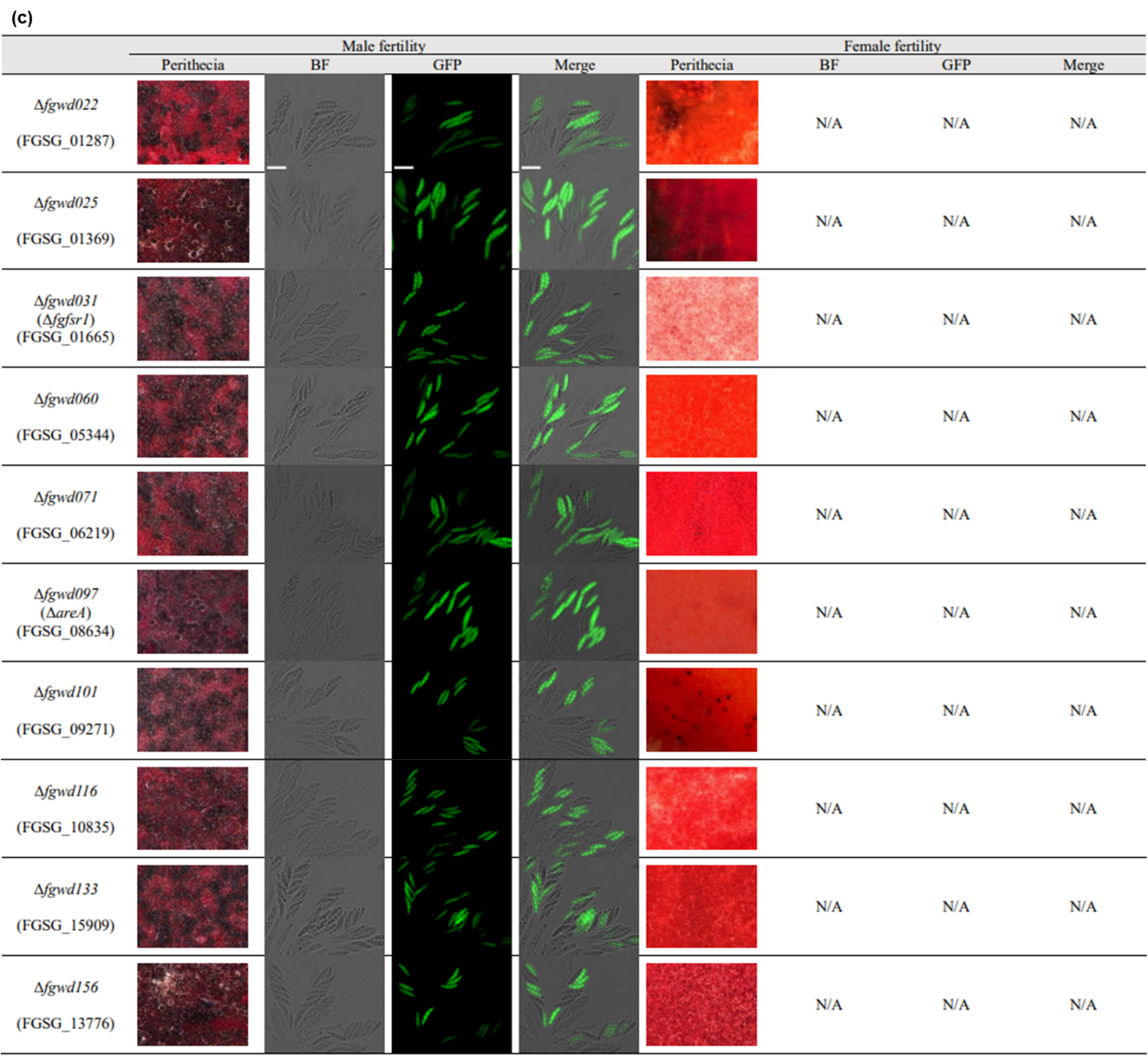

**Fig. S4. Assessment of male and female fertility in WD40 gene deletion mutants of *F. graminearum*.** Each strain was inoculated on carrot agar and crossed with ∆*mat1-1*/pIGPAPA-GFP. Photographs were taken 7–9 days after sexual induction. In normal asci, the eight ascospores exhibited a 1:1 segregation of GFP-positive and GFP-negative spores. Scale bar: black = 500 µm; white = 20 µm. **(a)** Mutants with normal male and female fertility. **(b)** Mutants producing partially normal perithecia but defective in ascospore formation in release. **(c)** Mutants exhibiting female-specific infertility. **(d)** Fgwd142 exhibited inadequate male and female fertility. ^a^ Not applicable. BF, bright field.

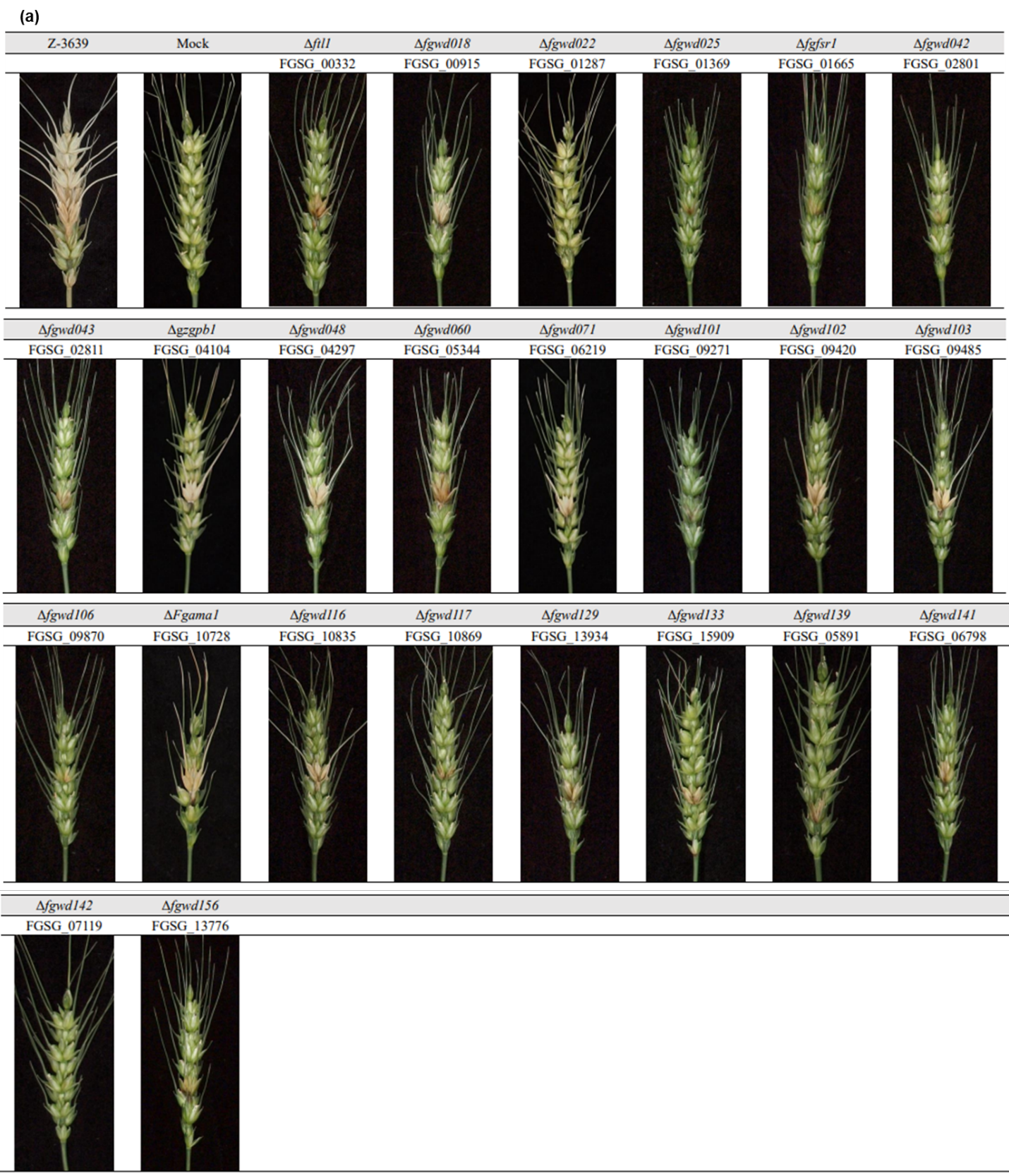

Continued on the following page.

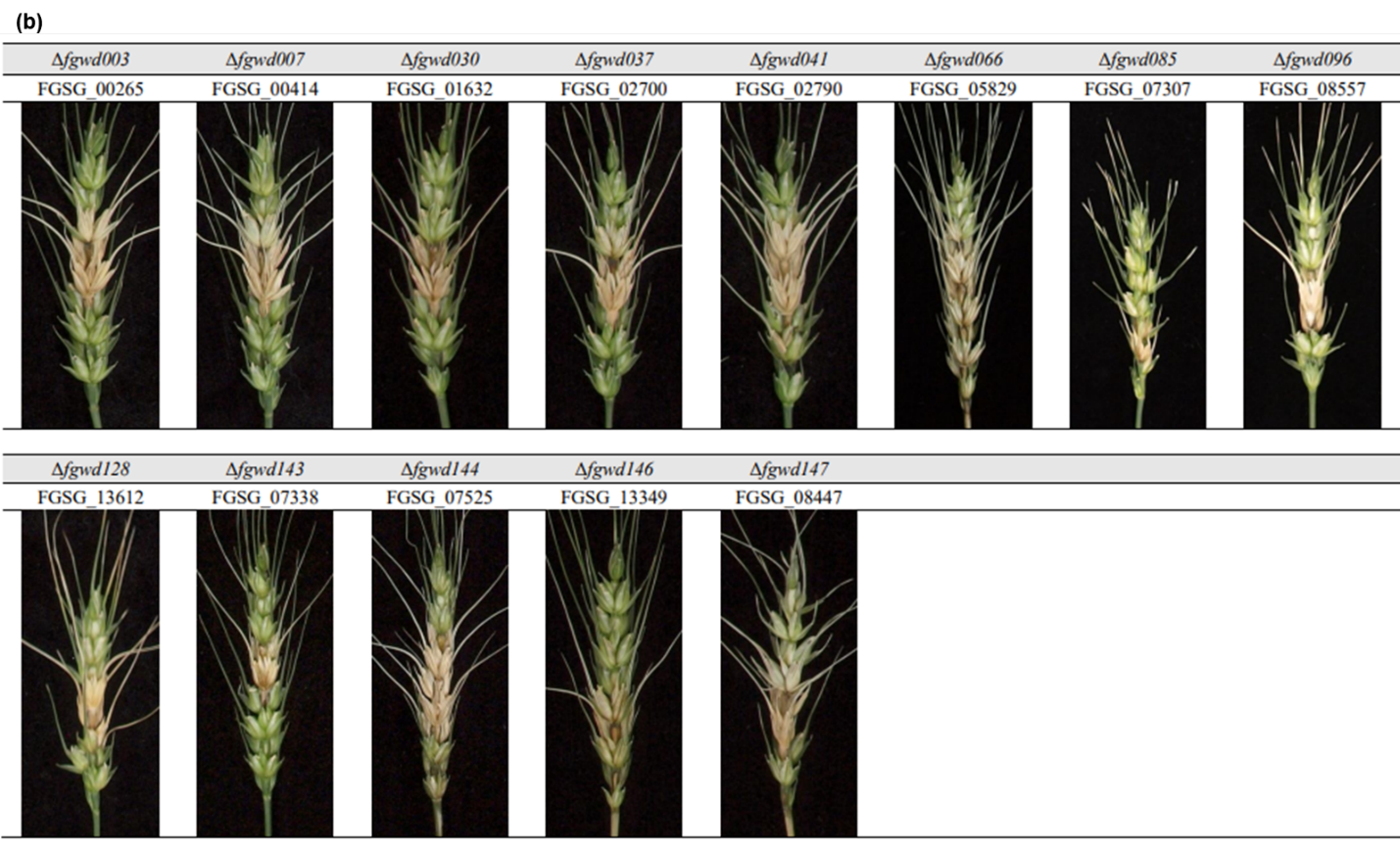

**Fig. S5. Virulence of *F. graminearum* WD40 deletion mutants on wheat heads.** Wheat heads were inoculated with the indicated strains and photographed at 21 days post-inoculation. The wild-type strain Z-3639 and mock treatment are shown for comparison. **(a)** WD40 gene deletion mutants in virulence group 1, showing colonization restricted to the inoculated spikelet and failure to spread to adjacent spikelets. **(b)** WD40 gene deletion mutants in virulence group 2, showing markedly reduced virulence compared with the wild-type strain.

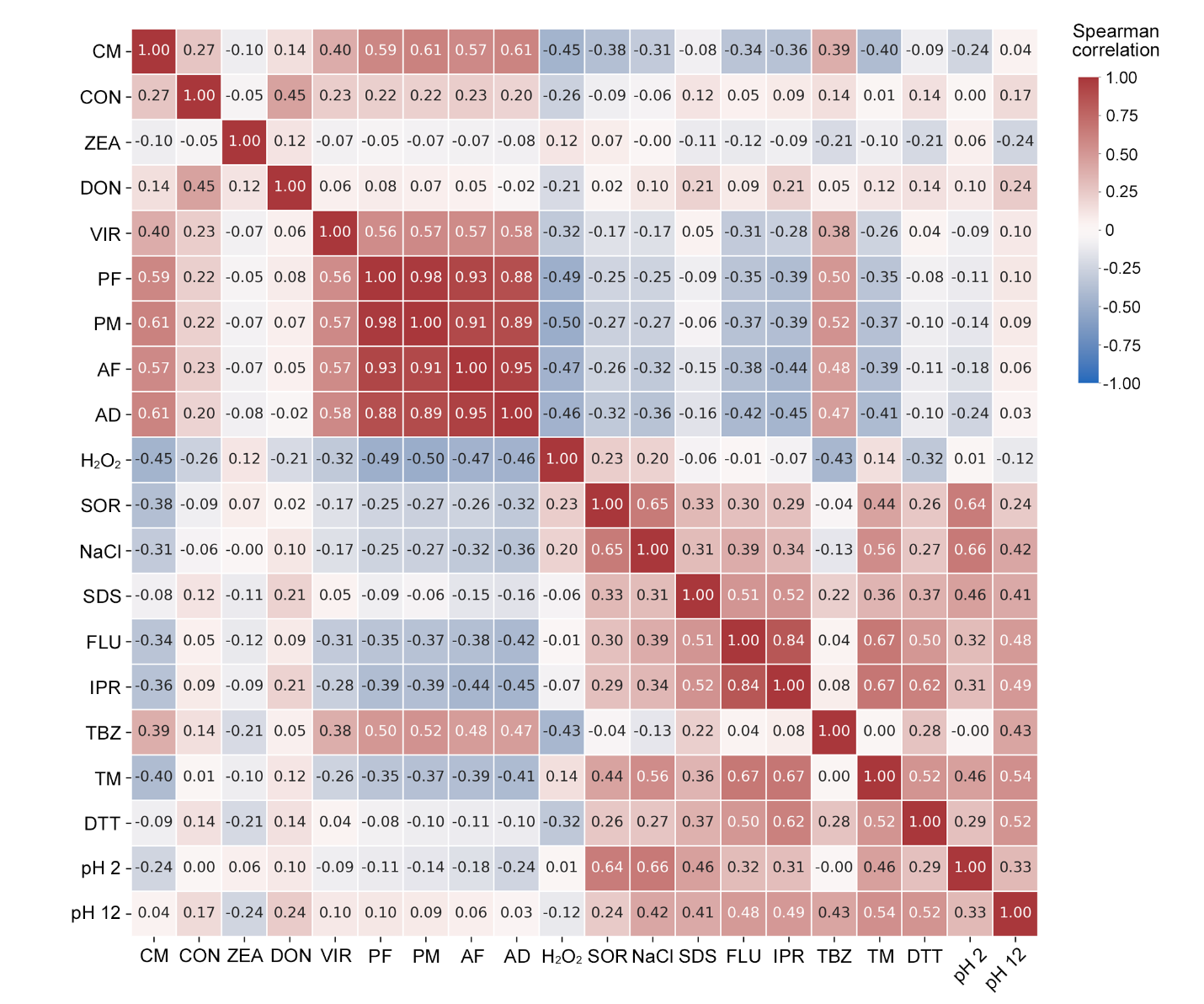

**Fig. S6. Phenotypic correlation analysis of *F. graminearum* WD40 gene deletion mutants.** Heat map showing Spearman’s correlation coefficients among phenotypic traits of WD40 gene deletion mutants. Red and blue indicate positive and negative correlations, respectively, with greater color intensity indicating stronger correlations. CM, complete medium; CON, conidiation; ZEA, zearalenone; DON, deoxynivalenol; VIR, virulence; PF, perithecia formation; PM, perithecia maturation; AF, ascospore formation; AD, ascospore discharge; H_2_O_2_, 0.31 mM hydrogen peroxide; SOR, 1 M sorbitol; NaCl, 1 M NaCl; SDS, 0.1 mg mL−1 sodium dodecyl sulfate; FLU, 0.05 μg mL−1 fludioxonil; IPR, 125 μg mL−1 iprodione; TBZ,0.625 μg mL−1 tebuconazole; TM, 2 μg mL−1 tunicamycin; DTT, 1.25 mM dithiothreitol.

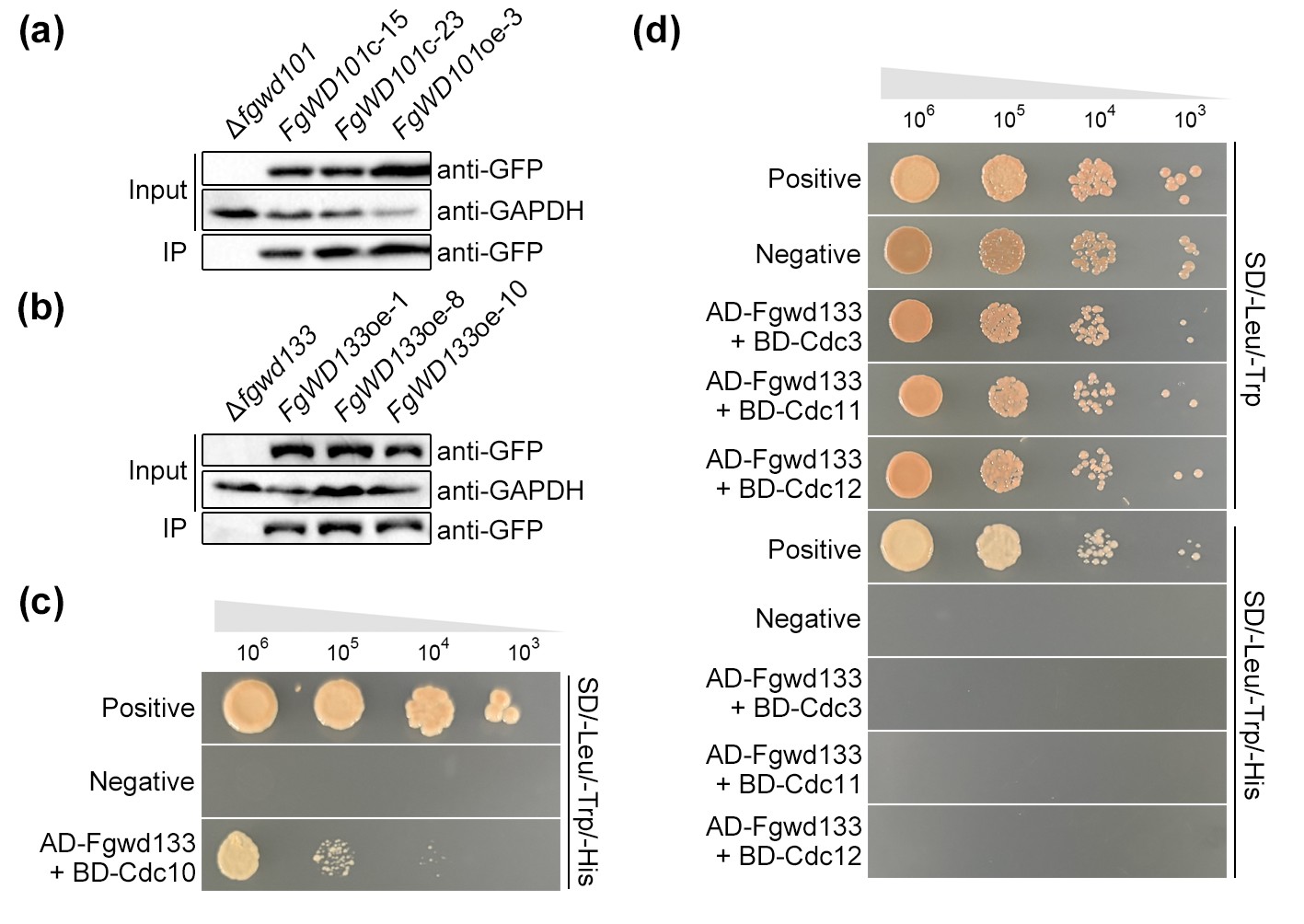

**Fig. S7. Immunoprecipitation of GFP-tagged Fgwd101 and Fgwd133 for affinity purification-mass spectrometry analysis and yeast two-hybrid confirmation of the interaction between Fgwd133 and septin complex proteins. (a,b)** Total protein extracts (Input) and proteins immunoprecipitated with anti-GFP magnetic beads (IP) from the GFP-tagged Fgwd101- and Fgwd133- expressing strains. GFP-tagged proteins were detected using an anti-GFP antibody. GAPDH was detected in the input samples using an anti-GAPDH antibody as a loading control. **(c)** Yeast two-hybrid (Y2H) interaction between Fgwd133 and Cdc10. Images were acquired after one additional day of incubation compared with Fig. 6i. **(d)** Y2H interactions between Fgwd133 and Cdc3, Cdc11, and Cdc12. The positive and negative control images are the same as those shown in Fig. 6i because the assays were performed simultaneously. AD, activation domain; BD, DNA-binding domain; SD, synthetic dropout medium.
